# Architectonic Spandrels in the Origin of Enzymatic Function

**DOI:** 10.64898/2026.02.09.704872

**Authors:** Miriam Poley-Gil, Miguel Fernandez-Martin, Alin Banka, Michael Heinzinger, Burkhard Rost, Alfonso Valencia, R. Gonzalo Parra

## Abstract

How new molecular functions emerge during protein evolution remains a fundamental question in molecular biology. The energy landscape theory states that proteins are minimally frustrated, i.e. they have minimized their internal conflicts, to allow robust folding. Yet, not all energetic conflicts are eliminated, with functional regions such as catalytic residues and ligand-binding sites being often enriched in frustrated interactions, trading localized stability for biological activity. However, it is still uncertain whether this functional frustration is an evolutionary adaptation, positively selected despite its energetic cost or an inevitable physical byproduct of the fold architecture. Here, we combine reverse folding, structure prediction, and sequence analysis with local frustration profiling to address this long-standing question. Unexpectedly, we found that reverse folding algorithms are unable to energetically minimize evolutionary conserved frustration at specific residues, even when detrimental to overall structural stability. We propose that these frustration hotspots act as architectural spandrels, inherent physical constraints of the fold that evolution subsequently co-opts for function. Our findings connect biophysical constraints and evolutionary selection, providing a new framework to understand how functional specificity emerges in protein landscapes.

---

For over half a century, protein biophysicists have sought to understand how protein sequences encode their structures, dynamics, and function(s) (1–4). Proteins must satisfy multiple, and often conflicting, constraints to fold into stable structures while remaining flexible and specific enough for binding, catalysis, and molecular interactions (5, 6). Dissecting how sequence features differentially contribute to structural integrity versus functional specificity remains a central challenge in molecular biology. Historically, this challenge has sparked debate over the relative roles of evolutionary selection and physical laws in shaping protein architecture. While the classical Darwinian view emphasizes natural selection as the primary driver of functional innovation, alternative perspectives suggest that protein folds are intrinsically constrained by the physical and chemical properties of polypeptides (7–10). Rather than opposing forces, physics defines the allowable structural and energetic landscape within which evolution operates. According to this view, many functional features are not sculpted *de novo* by natural selection, but emerge as intrinsic physical consequences of packing secondary structural elements (11). Evolutionary adaptation subsequently fine-tunes these physically determined structural features (12). However, the specific biophysical mechanisms linking protein fold architecture to the emergence of local functional features remain unknown.

Progress in disentangling these views has been hindered by a long-standing asymmetry between available sequences and experimentally determined protein structures (13). Fortunately, recent machine learning methods have offered new opportunities to revisit these questions. Structure predictions methods (14, 15) now provide high-quality models for most known proteins, narrowing the sequence–structure gap (16); and inverse-folding methods (17, 18) generate sequences compatible with a given backbone, enabling to explore how large is its sequence attractor (19, 20). Combined with the Energy Landscape Theory, these advances provide a powerful framework to investigate how physical constraints shape sequence variation, structural stability, and ultimately molecular function (21). This theory states that, according to the “minimum frustration principle”, proteins fold through funneled energy landscapes that have a strong bias toward the native state, minimizing conflicts among residues. Nonetheless, *≃* 10% of native interactions exhibit high frustration, as conflicting with the local structure (22).

Since function and stability often conflict (25), these frustrated interactions are thought to have been positively selected for functional and dynamic reasons. Previous studies have also shown that frustrated residues co-localized with functionally important regions (26–28). Here, we extend this paradigm by proposing that local frustration at certain sites can constitute an unavoidable physical constraint of the fold itself, independent of functionality. We explore the biophysical boundaries of well-characterized folds analyzing frustration conservation patterns (29) across reverse-folded sequence libraries, linking sequence variability to energetic constraints (Fig.1). We show that while some residues tolerate broad sequence and frustration variability, others remain constrained and persistently highly frustrated across designed sequences even when reverse-folding algorithms attempt to maximize structural stability. Some of these persistent constraints coincide with known functional sites. Thus, we propose that some local frustration does not arise through adaptive evolution but instead emerges unavoidably from the protein fold architecture, constituting a type of *spandrel-like* constraint in analogy to *architectural spandrels* (30–32) (Fig.5). These *spandrels* may predate biological functions and were subsequently co-opted via evolutionary exaptation (33). Our findings therefore suggest that protein folds do not merely constrain sequence evolution but actively shape the landscape of evolutionary possibilities by generating intrinsic energetic features upon which natural selection can act.

**Fig. 1.**
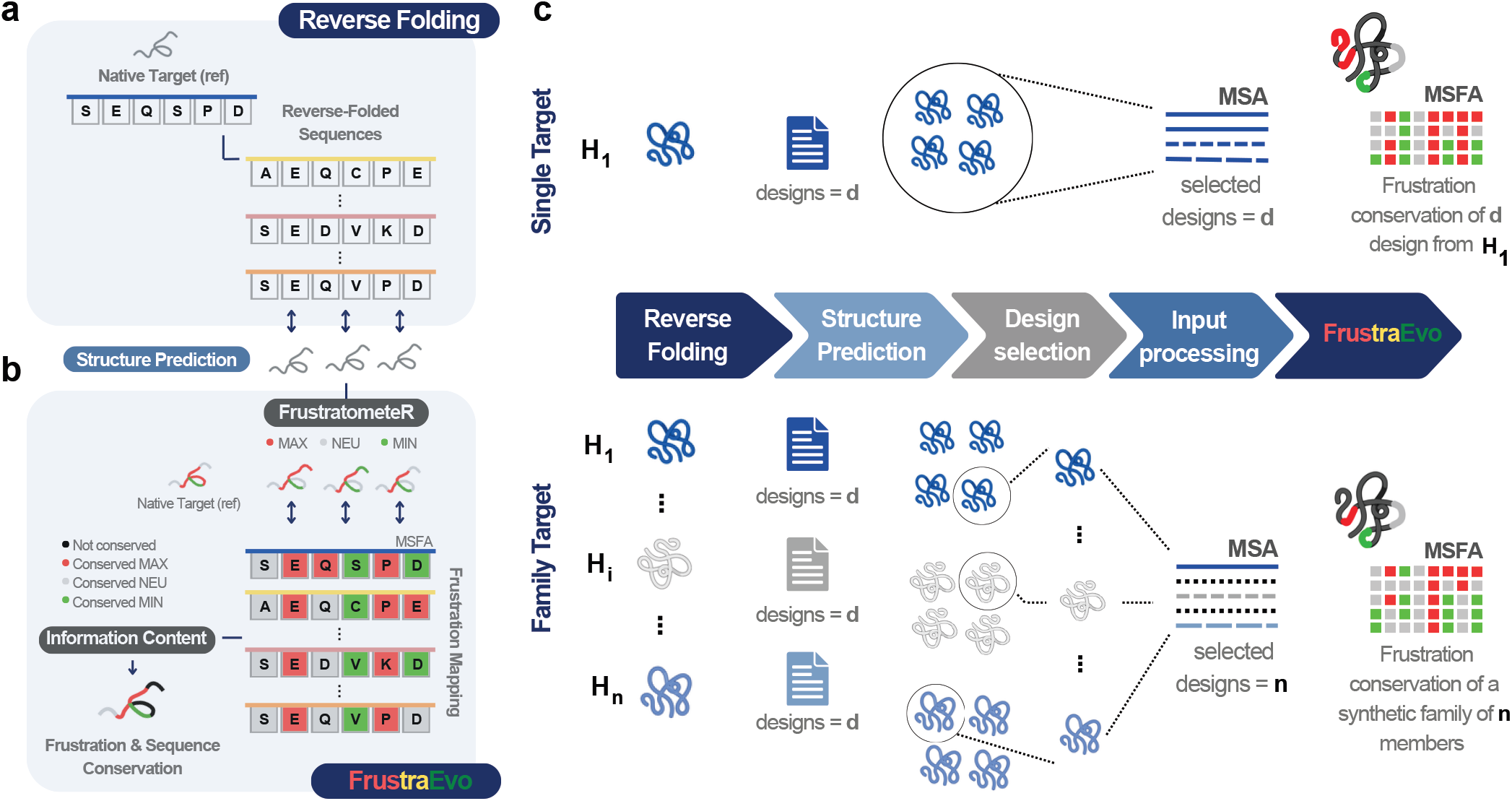
Exploration of frustration patterns and biophysical constraints of reverse-folded sequences. (a) Reverse-folded sequences are generated and structurally predicted to confirm matching with a given backbone. (b) Local frustration is computed for each design using FrustratometeR (23) and integrated across all designs with FrustraEvo (24), measuring frustration conservation in the columns of a MSA. Because residue identities may vary among designs, the frustration state at each position may also differ across models. By combining frustration information from all sequences and their MSA, FrustraEvo highlights conserved and variable frustration patterns, revealing which positions share similar energetic behavior and which can tolerate alternative residues. Comparing remodelled and invariant positions helps identify regions constrained by folding, stability, or function. (c) Overview of the proposed workflow in any target or protein family. Combining (a) and (b) we implement two complementary strategies. (1) Single Target approach: designs multiple sequences for one target, their predicted models are then analysed collectively to assess frustration conservation across the entire design set. (2) Family Target approach: evaluates a reverse-folding algorithm on a set of homologous structures. For each homologous or family member, a best-scoring design is selected, building a synthetic designed family. FrustraEvo then quantifies frustration conservation across this designed family, providing deeper coverage of the sequence space.

## Results

### Catalytic Residues in the *β*-lactamase Fold Resist Energetic Optimization with ProteinMPNN

The *β*-lactamase family consists of catalytic proteins that play a key role in antibiotic resistance in bacteria (34). We used a non-redundant dataset of experimentally determined native *β*-lactamase structures (n=31) to generate a Multiple Sequence and Frustration Alignment (MSFA) with FrustraEvo (24) and analysed the energetic diversity across alignable residues from the different family members. Our previous work (28) reported that five out of the six annotated catalytic residues in native *β*-lactamases are energetically conserved (MSFA columns that consistently exhibit a majoritary frustration state with respect to the background distribution, FrustIC*>*0.5), three as highly frustrated and two as neutral (Fig.2A). The native MSFA contains 256 positions (*SI Appendix, Fig*.*S1A*) from which 120 (47%) are energetically conserved. When classifying these conserved positions according to the frustration state that contributes the most to the energetic conservation we get 51 minimally frustrated (20%), 62 neutral (24%) and seven highly frustrated positions (3%) (*SI Appendix, Fig*.*S3D*). The remaining 136 positions (53%) are not energetically conserved, i.e., the distribution of frustration states for these residues is indistinguishable from the background distribution. To better understand the sequence, structure, and frustration relationships in this family we used ProteinMPNN (17) to reverse fold sequences that fit the 3D-coordinates of the different members of the *β*-lactamases family, and explored how much frustration variation their native structures can harbour (Fig.1). In a first approach that we refer to as Single Target we reverse folded multiple sequence designs (n=20) for a single protein target. In this case the reference used is the *β*-lactamase of *Bacillus licheniformis* (PDB ID: 4blm-A) (35), from which we took the residue numbering used in what follows. We predicted their structures with ESMFold and used them together with their corresponding MSA as inputs for FrustraEvo (Fig.2B). In a second approach that we refer to as Family Target we reverse folded multiple sequence designs (n=20) for each family member (n=31) and predicted their structures. Then, within each member design set, we selected the structural model that maximizes the pLDDT score to build a synthetic *β*-lactamases protein family composed of 31 designs, one from each family member (Fig.2C). In Fig.2D we show a full-sequence consensus MSFA that facilitates the comparison between the native *β*-lactamases (Fig.2A), the Single Target (Fig.2B) and the Family Target (Fig.2C) designs (*SI Appendix, Fig*.*S1*, full MSFA). Surprisingly, the native identities of two catalytic residues, K73 and K234, are maintained in all ProteinMPNN designs even when such identities introduce high frustration at the predicted structures. The catalytic residues S70 and S130 appear as conserved and neutral at the structure level as in the native family. For the remaining two catalytic residues, ProteinMPNN frequently assigns alternative identities, most notably for A237, which is substituted in the majority of designs. In the case of E166, a subset of designs incorporate variants that reduce local frustration (Fig.2B,C).

**Fig. 2.**
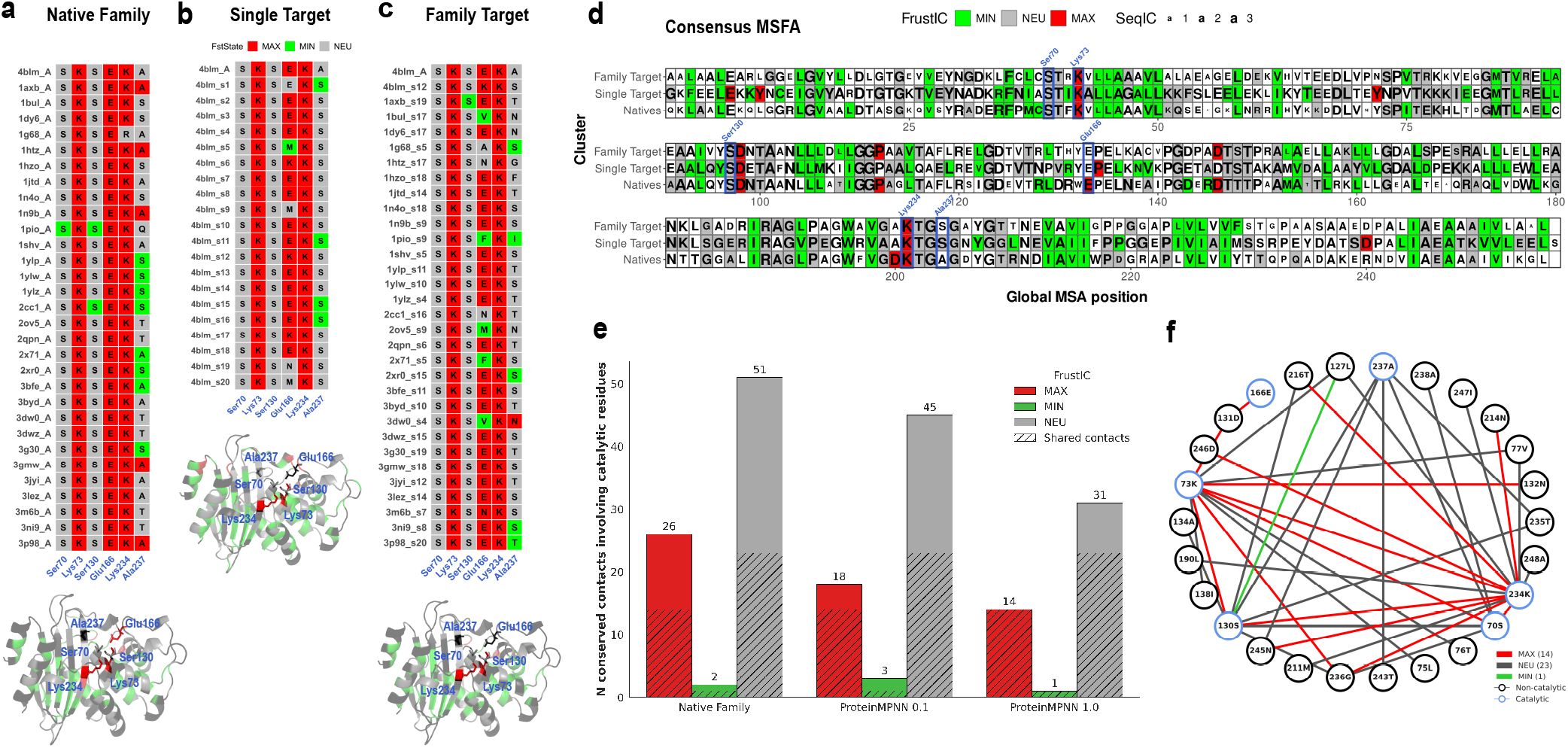
Frustration patterns in native and ProteinMPNN *β*-lactamases. (a) Subset from the native MSFA including the catalytic residues (S70, K73, S130, E166, K234, A237) annotated (numbering corresponding to the reference 4blm-A), including the SRFI results when using *β*-lactamases native structures (n=31). The MSFA represents the MSA on top of which the corresponding SRFI based states (i.e. minimally, green; neutral, gray; highly frustrated, red) are mapped. The same subset is shown for both (b) the 4blm-A Single Target approach (using 20 designs and the reference 4blm-A, n=21) and (c) the Family Target approach (using 1 design per native *β*-lactamase and the reference 4blm-A, n=32) generated at T=0.1. The designs chosen per native target (s number in y axis labels refers to sample number) are those that maximize the ESMFold pLDDT score in each case, among all those designed. Below, frustration conservation is displayed for each MSFA position, mapped onto the reference 4blm-A. Regions in black are positions with a non-conserved frustration state (FrustIC *≤* 0.5), the conserved ones (FrustIC *>* 0.5) are coloured according to the frustration state that contributes the most to the overall FrustIC value. (d) Consensus multiple MSFA full-sequence comparison, where the letter size reflects the Sequence Information Content (sequence conservation) and only positions with a conserved frustration state are colored accordingly, while non-conserved positions are shown in white. Catalytic residues are labeled in blue. Each cluster row summarises the FrustraEvo results for a given MSFA, containing SRFI data for the described datasets. The reference 4blm-A is included in each case to facilitate pattern comparison among runs. (e) MutFI results considering only the conserved contacts (i.e. non-conserved contacts not shown) involving the catalytic residues within the Native family, and within the Family Target approach at 0.1 and 1.0. The hatched regions refers to shared conserved contacts across the three datasets, which are plotted in the attached network (f). Catalytic residues are highlighted (blue nodes) differentially from the non-catalytics (black nodes). The contacts (edges) are colored according to the three frustration states.

All previous designs were generated with default parameters. The sampling temperature hyperparameter (T), in ProteinMPNN, promotes higher sequence variability, therefore we repeated the analyses varying T from 0.1 (default) to 1.0. Increasing T also results in higher frustration states variability for alignable positions in the MSFAs across designs (*SI Appendix, Fig*.*S2, S3C, S4C*). We observed a decrease in the overall percentage of conserved positions as T increases (*SI Appendix, Fig*.*S3D, S4D*). However, certain residues such as K73 and K234, remain conserved and highly frustrated even when sampling at maximum T. Although S70 and S130 have a neutral FrstIC at the Single Frustration Index (SRFI), we also previously reported (28) that these residues participate in a dense network of conserved and highly frustrated interactions (FrustIC based on the Mutational Frustration Index, MutFI, see Methods) involving the other catalytic residues: mainly K73 and K234, and, to a smaller degree E166. A237 is the only catalytic residue that is not part of the highly frustrated network of interactions and only displays neutral contacts including both serines. We next computed frustration conservation using MutFI to determine whether the persistent high-frustration signal identified by SRFI in the ProteinMPNN designs was also conserved at the interaction level. We further compared these contact-level conservation patterns of the native family with those observed in the Family Target approach generated at the lowest (T = 0.1) and highest (T = 1.0) tested sampling temperatures. As in the native proteins, all catalytic residues except A237 participated in a conserved network of highly frustrated interactions across the ProteinMPNN designs, partially preserving the native frustration network (Fig. 2E,F).

To further investigate the tendency of ProteinMPNN to maintain highly frustrated residues, we tested whether this effect persists when the target structures are modified to explicitly reduce local frustration. We used AlphaFold2 models where we systematically introduced frustration-minimizing mutations at key catalytic positions across the entire *β*-lactamases family. We also tested this effect in experimentally solved structures carrying frustration decreasing mutations. In both cases, ProteinMPNN designs consistently reintroduced native and highly frustrated identities at catalytic sites, confirming the robustness of this behavior (*SI Appendix, Note S1*).

### Persistence of Local Frustration in the *β*-lactamase Fold with Alternative Inverse Folding Strategies

Because catalytic sites are highly conserved and restricted in amino acid composition (36), ProteinMPNN may have memorized these architectures, imprinting expected catalytic identities onto enzyme-like backbones. We tested this possibility by retraining ProteinMPNN on training sets excluding either (1) all annotated hydrolases or (2) all annotated enzymes in the PDB (37). We used the contrastive learning-based CLEAN algorithm (38) to identify potentially unannotated enzymes in the ProteinMPNN training set. Applying this additional annotation layer, we obtained four retrained versions of ProteinMPNN: PDB-Hydrolases, PDB-AllEnzymes, CLEAN-Hydrolases, and CLEAN-AllEnzymes (*SI Appendix, Figs*.*S10, S11, Table S1*). Then we compared the original and retrained versions using the Family Target approach on the natural *β*-lactamases set at minimum and maximum sampling temperatures.

Designs from retrained models showed pLDDT distributions similar to the original ProteinMPNN at each temperature, with lower scores at higher temperatures (*SI Appendix, Fig*.*S12A*). Consistent with previous results, the percentage of residues with a non-conserved frustration state increases with temperature, while the conserved residues across the three frustration states remains consistent across models, being more native-like at lower temperatures in all cases (*SI Appendix, Fig*.*S12C*).

Of the six catalytic residues, S70, K73 and K234 are consistently recovered in the designs from all models in both sampling temperatures (*SI Appendix, Fig*.*S13A*,*B*). S130 is recovered in all cases except for the CLEAN-AllEnzymes where it gets substituted by a glycine at both temperatures. E166 and A237 are largely remodeled in agreement with results from the previous section. However, results obtained with the CLEAN-AllEnzymes model should be interpreted cautiously as the training dataset was reduced over 50% and therefore its ability to generate biophysically sounding proteins is not guaranteed.

Despite that, native identities are mostly recovered at catalytic sites even when they are not always highly frustrated as a consequence of their interacting residues being highly remodelled (*SI Appendix, Fig*.*S13*).

Notably, the non-catalytic residue D131 is also consistently recovered by ProteinMPNN as conserved and highly frustrated across all models at both temperatures. We additionally generated the 4blm-A D131V mutant, predicted its structure, and performed Single Target, consistently recovering D131 as conserved and highly frustrated across the ProteinMPNN designs (*SI Appendix, Fig. S14*). Thus, the recovery of this non-catalytic residue by ProteinMPNN, despite its location within the SDN loop and its side chain pointing away from the active site (39), is unlikely to be a memorization artifact.

However, the way ProteinMPNN was trained introduces a potential limitation: its objective function maximizes Native Sequence Recovery (NSR) for a given backbone. Because natural sequences evolve under multiple constraints (40), not only functional ones but also those ensuring efficient and accurate folding, it is possible that some frustration patterns observed in native proteins originally arose to assist the folding process (41). Once folding is complete, these residues may no longer play an active structural role, although they could later be co-opted for additional functions over evolutionary time. ProteinMPNN may have learned these folding constraints, incorporating them into its designs.

To further rule out memorization artifacts by Protein-MPNN we have repeated some analyses using Caliby (18), a Potts-based reverse-folding model that optimizes the sequence-structure fit rather than NSR (*SI Appendix, Note S4*). We also used alternative frustration parameter values that are better aligned with folding simulation analysis (*SI Appendix, Note S5*). Results showed that although some frustration retained by ProteinMPNN disappears, the catalytic K73 and the supporting residue D131 in the SDN loop retain high frustration across all design approaches, including Caliby. High frustration at K234 is conserved in ProteinMPNN, whereas *≃* 20% of Caliby designs substitute this position, reducing its frustration. A summary of how the conserved and highly frustrated residues in the native family behave across design strategies is provided (*SI Appendix, Table S2*). A similar pattern was observed for K117 in the Ras family. Although this residue interacts with the nucleotide substrate and is retained by ProteinMPNN as highly frustrated, the signal is not preserved in Caliby designs (*SI Appendix, Note S3*).

### Decoupling Function from Fold Architecture: Functional Residues in the *α*-globin Fold can be Energetically Minimized by ProteinMPNN

As a counterexample to our findings in the catalytic *β*-lactamases fold, we analyzed the non-enzymatic *α*-globins family (*SI Appendix, Note S2*), where highly frustrated native residues are predominantly conserved to mediate functional protein-protein interactions (PPIs). Using both ProteinMPNN and Caliby reverse-folding approaches, we found that these functional sites are systematically remodeled toward lower frustration states (*SI Appendix, Fig*.*S17-S19, Table S3)*, with almost all native frustration removed at higher sampling temperatures (*SI Appendix, Fig*.*S20-S22)*. This differential behavior suggest that functional and structural constraints can be effectively decoupled in *α*-globins, and that their native frustration most likely reflects adaptive, functional demands that can be minimized during stability-driven redesign.

### Persistent Frustration in Redesigned Proteins Folds

Early *de novo* designed proteins lack evolutionary history and were optimized for having stable, minimally frustrated energy landscapes without specific functional constraints.

Top7, a 93-residue *α*/*β* protein with a novel topology (PDB ID: 1qys (42)), was the first successful globular *de novo* design. Despite closely matching its predicted structure (1.2 Å RMSD), Top7 exhibits non-cooperative folding with multiple intermediates, being such behaviour a direct consequence of the target structure (43). Local frustration analysis identified five highly frustrated residues in native Top7 (*≃* 5% of the protein: Y21, N34, Y39, D78, D84) using the SRFI (Fig.3A), while MutFI revealed a few highly frustrated surface contacts (Fig.3B). ProteinMPNN redesigns showed high structural confidence (mean pLDDT 87.91 vs. 81.01 for native Top7 if predicted) and 0.44 mean sequence recovery (Fig.3C-E), but still all retained at least five highly frustrated residues with no positional overlap with original design hotspots. Some frustrated positions were recurrent across designs, mainly positions 22 (D/E) and 43 (consistently P) (T24 and A45 from original Top7 sequence numbering) (*SI Appendix, Fig*.*S23*). Additionally, all designs include S49 and Y74 (originally S51 and Y76), which, while predominantly highly frustrated, can also adopt other energetic states due to differences in the local interacting neighborhood either in terms of conformation or identities.

**Fig. 3.**
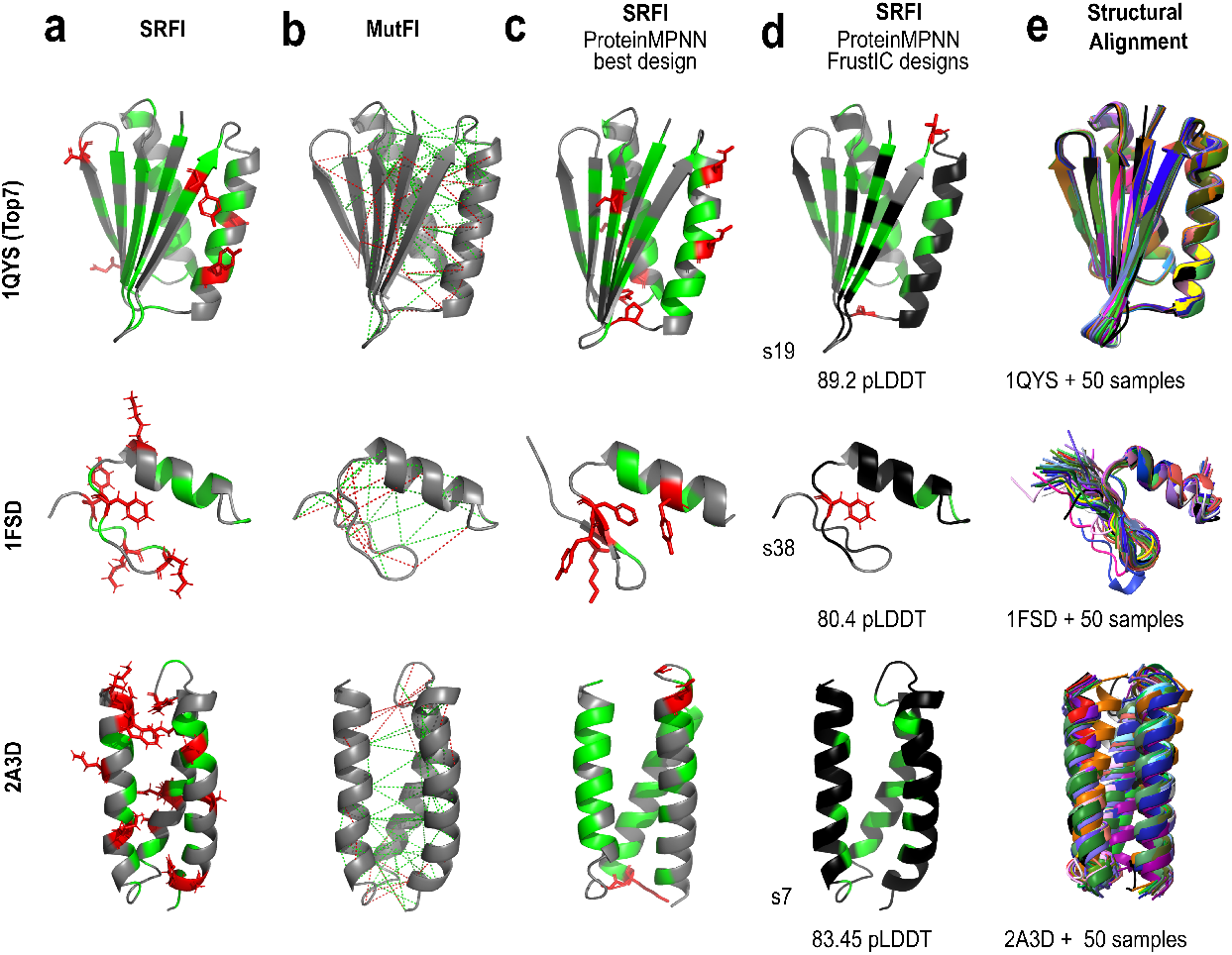
Evaluating frustration in *de novo* proteins re-designed by ProteinMPNN. For each *de novo* protein we show: (a) SRFI frustration pattern for *de novo* target, where positions are colored according to the three frustration states. (b) MutFI frustration pattern for *de novo* target, just showing short and long contacts colored according to if they are highly or minimally frustrated. (c) SRFI frustration pattern in the best structural design (s refers to sample number, corresponding to the design with higher ESMFold pLDDT score) proposed by the Single Target approach. (d) SRFI frustration conservation pattern resulting for the 50 derived designs from the Single Target approach, where regions in black are positions with a non-conserved frustration state (FrustIC *≤*0.5) and the conserved ones (FrustIC *>*0.5) are colored according to the frustration state that contributes the most to the overall FrustIC value. (e) Structural alignment considering 50 folded designs target-like. In any case and even if not visible because of overlapping, the reference structure is colored in black within the structural alignment.

Similar behavior was observed in other *de novo* proteins. The 28-residue long *ββα* zinc-finger-like motif (PDB ID: 1fsd (44)) showed at least one highly frustrated and conserved position in both native and redesigned structures (Fig.3D), while *>*70% of positions showing non-conserved frustration (*SI Appendix, Fig*.*S24A*). Mean-while, in the complex 73-residue 3-helix bundle *de novo* protein, *α*3D (PDB ID: 2a3d (45)), only a subset of ProteinMPNN redesigns adopted the target fold (Fig.3E). Highly frustrated positions were not fully alignable but still scattered across redesigns in a non-conserved way with a minimum of two positions per design (*SI Appendix, Fig*.*S24B*). This suggests that many alternative sequences can be accommodated within a scaffold that require the occurrence of non-adaptive frustration to satisfy the structural constraints imposed by that fold. Reverse folding with Caliby only confirmed that two residues remain consistently highly frustrated in Top7, matching ProteinMPNN proposals and supporting a structural rather than functional origin of local frustration (*SI Appendix, Fig*.*S29A, Note S4, Note S5*).

### Frustration as an Intrinsic Feature of Native Protein Folds

Evaluating reverse-folded sequences from the *β*-lactamase fold showed that certain local frustration, often involving functional residues, cannot be minimized. Similarly, some frustration sistematically persists in *de novo* designed proteins such as Top7. To test how general this effect is, we analyzed a dataset of 488 enzymatic functional families (FunFams), containing 15% of mainly *α*, 16% of mainly *β*, and 69% of *α*/*β* folds according to the CATH classes (49). By comparing native frustration conservation profiles with those of ProteinMPNN and Caliby designs (Fig.4A), we found that 74 (*≃* 3%) of the 2,524 positions natively conserved and highly frustrated remain consistently constrained in the same way by consensus across all design configurations (Fig.4B). This estimate considered a conservative lower bound for the recurrent frustration in native folds, based on an initial, albeit limited, protein dataset. Considering a less restrictive filtering criteria would increase this number to 137 (*≃* 5%), based solely on the intersection of ProteinMPNN 0.1 and Caliby with the native set (*SI Appendix, Table S4*).

**Fig. 4.**
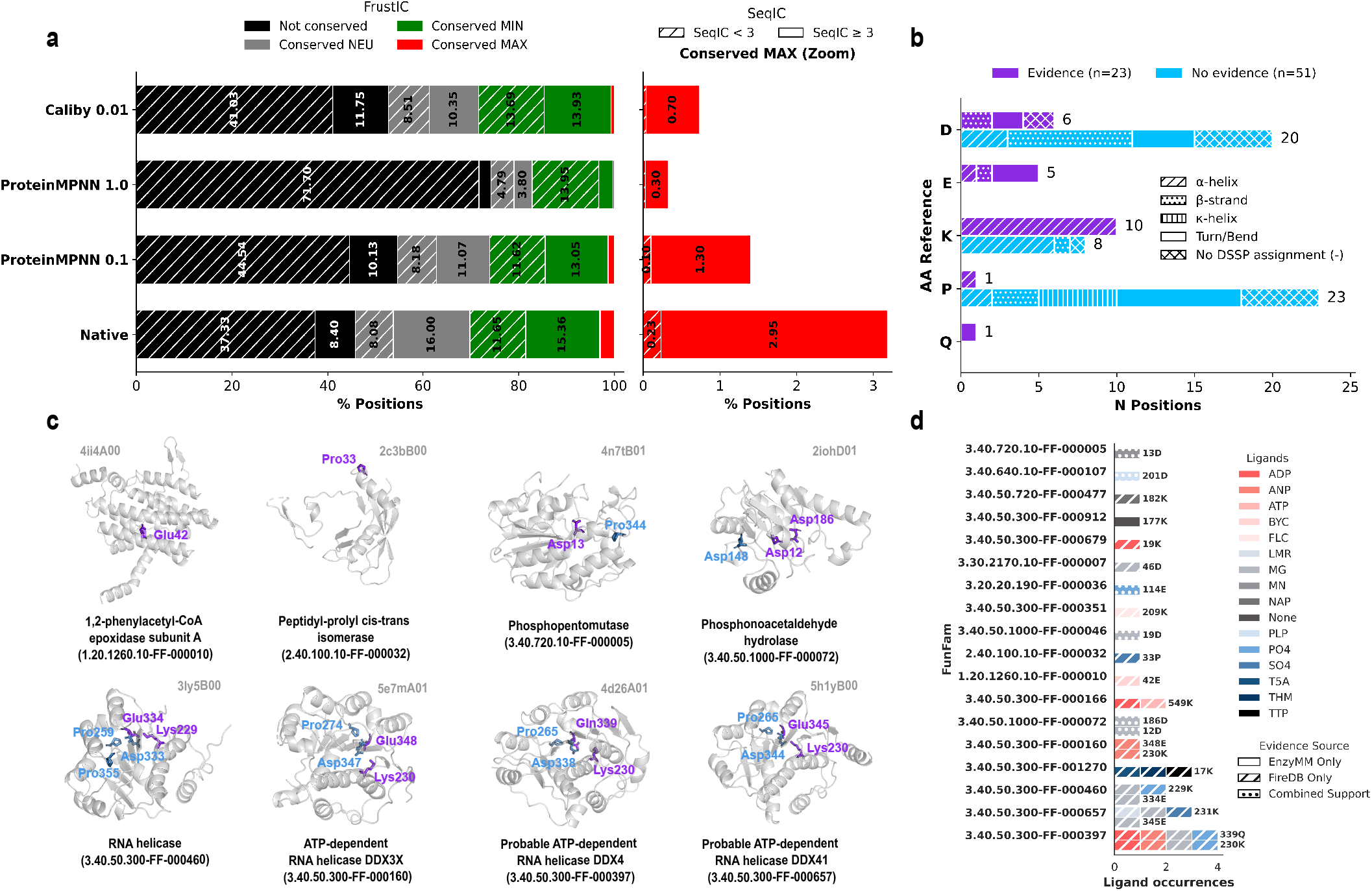
Frustration conservation across FunFams and design configurations. (a) Frustration (FrustIC) and sequence (SeqIC) conservation over MSA positions from 488 FunFams across design configurations using FrustraEvo (SeqDist=3). Colors correspond to the conserved FrustIC states (green: conserved minimally frustrated; grey: conserved neutral; red: conserved highly frustrated; black: not conserved). Hatched regions represent sequence conservation (SeqIC). Conservation thresholds are set at FrustIC *>* 0.5 and SeqIC *≥* 3. MSA positions with zero or non-available SeqIC, as reported by the sequence-logo analysis, were excluded. (b) Distribution of consensus positions conserved as highly frustrated across designs configurations, with (purple, n=23) and without (blue, n=51) functional evidence. Data are stratified by amino acid identity (using identities from the native reference proteins) and secondary structure (using consensus DSSP (46) across MSA positions from each native FunFam). Note that unstructured positions (turns and bends) have been collapsed for the sake of simplicity. (c) Structural mapping of representative consensus positions on the native reference proteins, showcasing examples from the three main CATH structural classes, derived from secondary structure content. (d) Functional annotation layer for the 23 consensus positions with evidence, classified according to EnzyMM (47) and FireDB (48) functional sources. These positions are distributed across 18 FunFams.

Among those 74 persistent hotspots, 23 residues are identified as catalytic by the EnzyMM geometric template matching (47) and/or annotated as functionally relevant in FireDB (48), whereas the remaining 51 residues showed the same conserved high frustration pattern despite not having functional evidence. Functionally annotated sites are restricted to just five identities (K, Q, E, D, P) mainly within rigid *α*-helices across 18 FunFams, while those lacking functional evidence map to an even narrower chemical set (D, K, P) within more diverse motifs across 44 FunFams, six of which also harbor a conserved, functionally annotated site (Fig.4B).

Given that highly frustrated catalytic residues in *β*-lactamases are predominantly buried, whereas the PPIs in *α*-globins are more solvent exposed, we asked whether solvent accessibility could explain the persistence of frustration upon reverse folding. We therefore examined the relative solvent accessibility of such frustrated hotspots identified across protein families. These positions spanned both solvent-exposed and buried environments, indicating that solvent accessibility alone does not determine whether native frustration can be energetically minimized through reverse folding (*SI Appendix, Fig*.*S31*).

In total, 56 of 488 FunFams (11.47%) contain at least one natively conserved, highly frustrated position that resists energetic minimization by reverse-folding, retaining examples from *α, β* and *α*/*β* CATH classes (49). Notably, the six FunFams retaining both annotated and unannotated frustration hotspots converge on a single evolutionary lineage: all belong to the 3-layer *αβα*-sandwich architecture (3.40), five share the Rossmann fold topology (3.40.50), and four specifically map to P-loop NTPases (3.40.50.300) (Fig.4C). Furthermore, functional annotations for these sites are enriched in co-factor and ligand binding, particularly nucleoside phosphates (e.g., ATP, ADP), divalent cations (e.g., Mg^2+^) and metabolic co-factors such as NADP and PLP (Fig.4D) (50). More interestingly, 112 positions neither frustrated nor conserved in native FunFams often became so by consensus across ProteinMPNN 0.1 and Caliby, mostly at P/D-rich sites without functional annotation (*SI Appendix, Fig*.*S31B, Table S4)*. Considering ProteinMPNN 0.1 or Caliby individually, would increase this number to 528 or 417 respectively. These results, as shown in non-alignable frustrated positions of *de novo* proteins, suggest that some native scaffolds would also require non-adaptive frustration to satisfy fold-specific structural constraints even when a slightly different sequence is accommodated.

## Discussion

The existence of protein families sharing a common fold but exhibiting extensive sequence diversity implies that different regions of a fold are subject to fundamentally different evolutionary constraints. We combined inverse-folding algorithms with local energetic frustration analysis to identify which positions tolerate sequence variation and which remain constrained by the underlying biophysics across the sequence space compatible with a given fold. We found a remarkably consistent pattern: while many positions within a fold tolerate broad sequence diversity, certain residues remain energetically constrained. Some are conserved and minimally frustrated residues that act as structural anchors required for stability and foldability (22, 51). Unexpectedly, other positions, frequently associated to known functional sites, remain persistently highly frustrated across designs, even when this compromises local stability. Additionally, we identified residues that remain highly frustrated in sequences reverse-folded for *de novo* protein backbones, even though these proteins lack any known biological function other than folding. This observation extends the prevailing interpretation of local frustration as a predominantly adaptive feature selected for function (26–29).

We observed two distinct behaviors among highly frustrated sites previously associated with function. In some cases, frustration at functional sites could be decreased through reverse folding, as exemplified by *α*-globins. This indicates that functional and structural constraints can be decoupled, allowing stability-driven redesign to remove energetic conflicts associated with function. This is consistent with the known trade-off between function and stability (6), and with reports that ProteinMPNN can remodel functional sites and impair activity unless key residues are explicitly fixed (52). In contrast, in catalytic folds such as *β*-lactamases, several catalytic and non-catalytic positions could not be minimized by reverse folding, including the structurally constrained D131 in the SDN loop. This persistence was observed even when redesign was performed from stabilizing *in silico* or experimentally determined mutant structures. Since some residual local frustration has been proposed to contribute to robust protein folding (41), ProteinMPNN could recover these frustration signals as a consequence of its NSR objective rather than because they reflect architectonic constraints. However, retrained ProteinMPNN models depleted of enzymes still retained similar local frustration patterns. Moreover, Caliby, an orthogonal reverse-folding method, also recovered comparable frustration, arguing against an explanation based solely on information learned by ProteinMPNN or NSR inherent optimization biases.

Even with these controls, it is not possible to completely rule out the possibility that persistent local frustration reflects evolutionary information learned from natural proteins. We therefore turned to *de novo* proteins, which lack any evolutionary history and were designed solely to achieve stable folds. Remarkably, both human-engineered and machine-generated proteins retained highly frustrated regions after redesign, despite the absence of evolved biological functions. Although the specific frustrated positions differed among human and machine designs, persistent local frustration consistently re-emerged. These observations further support that local energetic conflicts can arise as an intrinsic consequence of the protein fold architecture (53).

Extending this analysis to a dataset of 488 enzymatic FunFams, we identified a broader pattern of persistent frustration affecting both positions with and without known functional evidence. These sites were restricted to a few amino acid identities (K, Q, E, D, P) and mostly found within P-loop NTPases adopting the Rossmann fold. This suggests that, in some cases, particular sequence identities at specific positions represent structural compromises imposed by protein fold architecture rather than independently selected functional features (54–57). These architectonic constraints may contribute to widespread protein frustration despite involving residues that are neither catalytically prominent (36, 58) nor prebiotically abundant (59–61). Furthermore, the enrichment of these frustration hotspots in ancient and abundant protein lineages (62–64) further raises the possibility that structural constraints inherent to ancient folds contributed to the emergence and diversification of early functional proteins (50, 61, 65). Recent work proposed that these versatile folds drove early metabolic expansion, followed by adaptation of existing folds rather than the emergence of new ones (66).

Taken together, these observations suggest that local protein frustration can arise through two distinct mechanisms (Fig.5B.3). *Adaptive frustration* consists of localized energetic conflicts that are retained evolutionarily because they provide a selective advantage, presumably of a functional nature; however, if such functional constraints are relaxed or masked, this frustration can be alleviated (Fig.5B1, C–E). In contrast, *architectural frustration* arises as an unavoidable consequence of the geometric and packing constraints required to build a particular protein fold (Fig.5B2, C–E). By analogy with *architectural spandrels* (30, 31), i.e. the unavoidable triangular spaces formed when domes are supported by arches (Fig.5E), we refer to this class of energetic conflicts as *spandrel-like frustration*. These constraints need not be fixed at identical residue positions across all sequences compatible with the same fold. Instead, different sequence solutions can satisfy the same architectural requirements while localizing unavoidable energetic conflicts at distinct positions. Consequently, a protein fold defines not a single optimal sequence, but a constrained ensemble of compatible sequence solutions, each containing its own pattern of *spandrel-like frustration*. Once these architectonic frustration hotspots arise, they become byproducts upon which natural selection can subsequently act, with a subset eventually becoming exapted into new molecular functions (32).

**Fig. 5.**
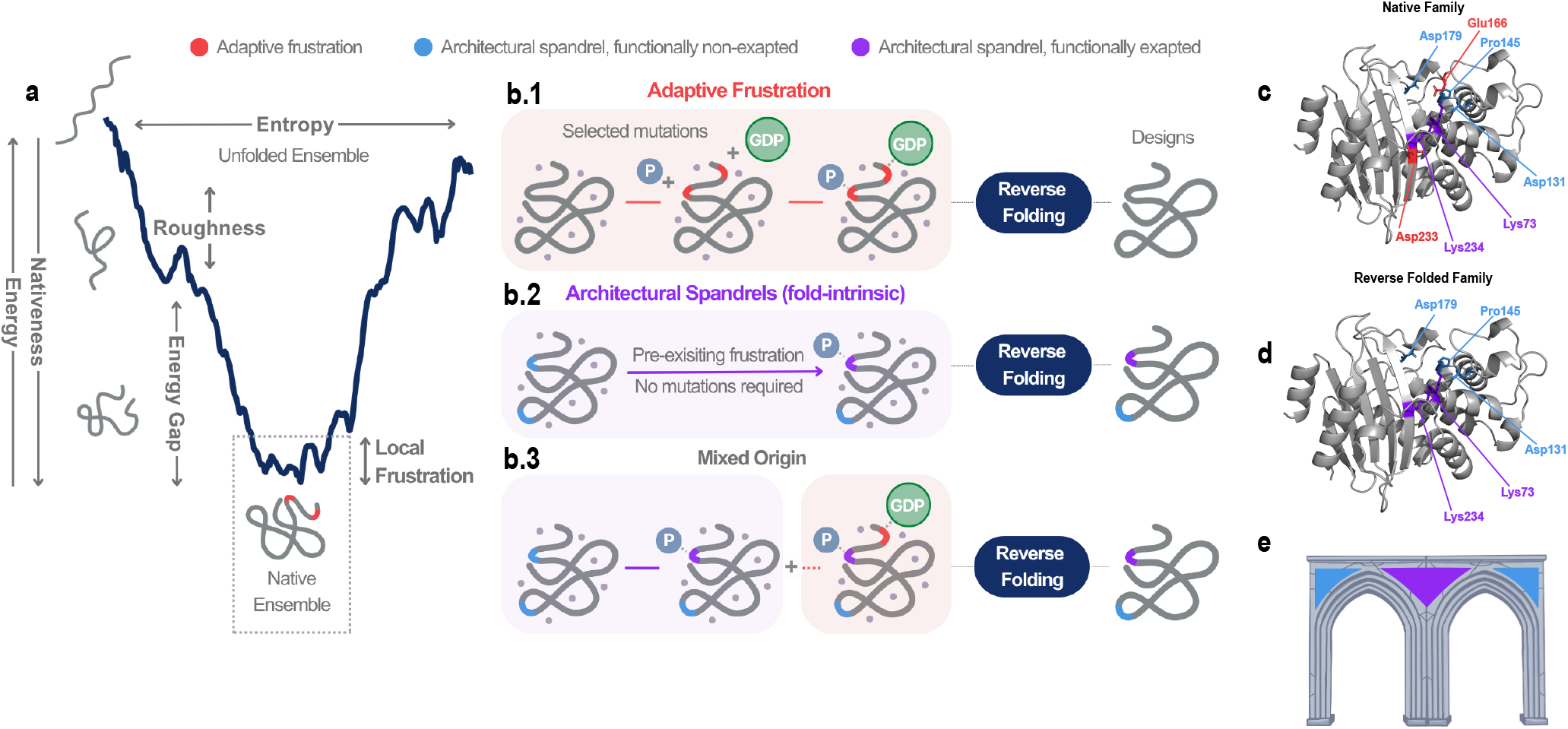
How reverse-folding algorithms handle conserved structural and functional constraints. (a) Schematic representation of the protein folding energy landscape. Proteins fold by minimizing internal energetic conflicts according to the minimum frustration principle, producing a funneled landscape biased toward the native state, while residual frustrated interactions generate local roughness. (b.1) Classical view in which frustration mainly reflects adaptive functional constraints (red patches). (b.2) Alternative view in which frustration arises from unavoidable structural fold constraints (architectural spandrels, blue patches), which may later become functionally exapted (purple patches). (b.3) Mixed model supporting 1) the shared origin of frustration and functional sites in folds and 2) that both adaptation and structural constraints contribute to protein functional diversification. (c,d) Example from the *β*-lactamase family comparing native proteins (top) and ProteinMPNN reverse-folded designs (bottom). Alleviated frustrated residues are proposed to be functionally adaptive (E166, D233). Hardwired frustrated residues are proposed to be structurally constrained (K73, K234, D131, P145, D179) and sometimes functionally exapted (K73, K234). (e) Analogy to architectural spandrels, triangular spaces that emerge inevitably when domes are built on arches (30, 31).

Consistent with this interpretation, the *spandrel-like* constraints retained in our designs can be grouped into distinct classes. Some *spandrel-like* positions coincide with residues that are already conserved and highly frustrated in native proteins and have known functional roles. We interpret these as *exapted native spandrels*: architectonic constraints that have been retained and co-opted for function during evolution. Other spandrels coincide with positions that are also conserved and highly frustrated in native proteins but lack known functional annotations, potentially representing *non-exapted native spandrels* or foldability requirements that lack a function in the folded state. Finally, we also identified positions that became conserved and highly frustrated across reverse-folded sequences but were neither conserved nor highly frustrated in the corresponding native families. These *non-native spandrels* represent alternative energetic compromises arising from different sequence solutions compatible with the same fold, which may have been explored and subsequently lost during evolution or may correspond to regions of sequence space not sampled yet by natural proteins. *Non-native spandrels* are particularly relevant because their absence from natural families makes native motif memorization less likely.

Thus, we propose that structural constraints can precede functional adaptation by providing latent opportunities for protein innovation through locally unstable environments, that can subsequently be exploited for function (11, 32). This view resonates with the concept of *Platonic folds*, which posits that the repertoire of protein structures is constrained by universal physical laws (8– 10, 67), as reflected in the conservation of structure despite extensive sequence divergence (2, 68, 69) and the uneven distribution of folds across structural space (70). But more importantly, it reframes the relationship between fold and function: rather than functional sites imposing an independent energetic penalty on an otherwise stable structure, this cost may be largely “pre-paid” by the fold itself. Within the metastable regime of proteins, evolution can therefore recruit structurally constrained, energetically costly positions for new capabilities without incurring the full cost of creating such environments *de novo*. In this sense, physical constraints imposed by the fold may act as latent substrates for functional innovation (11, 71, 72).

## Materials and Methods

### ProteinMPNN Structure-Conditioned Protein Sequence Design

Sequences designed to explore differential frustration patterns were generated by ProteinMPNN (17) using the full trained protein backbone vanilla model *v 48 020*.*pt*, which was originally trained with default flags num neighbors=48, backbone noise=0.20 and num epochs=150. Default parameters were also considered for the generation (without setting backbone noise, fixed positions or amino acids bias as the main considerations). To explore how the generation varies according to the sampling temperature hyperparameter, we cover a range from 0.1 (default) to 1.0, generating multiple sequences per target. Note that the minimum sampling temperature tested in our analyses coincides with the ProteinMPNN default.

### ProteinMPNN Re-training and Enzyme Function Prediction

We retrieved the complete ProteinMPNN training dataset, a multi-chain training data (16.5 GB, PDB bio units, 2021 August 2) containing 555,720 entries, of which 163,529 are unique PDBs. Then, we re-trained the model customising such a dataset by (1) removing all the experimental entries that were annotated as hydrolases (2) removing all the experimental entries that were annotated as any class of enzymes. The annotation related to the Enzyme Commission (E.C) number was retrieved from the PDB database (by July 2024) by using the advanced search with *Structure Determination Methodology* set to *experimental* and *Enzyme Classification Number* to *any of 3* (40,222 structures classified as hydrolases) or not empty (101,556 structures classified as enzymes) for filtering only hydrolases and all the enzymes respectively. Not all the structures annotated in the PDB were present in the ProteinMPNN training dataset. Once filtered, the new training dataset without hydrolases contains 463,204 entries (131,929 unique PDBs) assuming a total of *≃* 19% of PDBs removed from the original dataset. Similarly, the new training dataset without any enzyme contains 314,016 entries (81,502 unique PDBs) assuming a total of *≃* 50% of PDBs removed from the original dataset. However, in order to complete the annotation provided by the PDB database for the E.C number classification, we used the CLEAN software (38). CLEAN makes a function prediction per protein sequence according to the pairwise distances between the query sequence and all the functional cluster centres from the ESM-1b embeddings (esm1b t33 650M UR50S). We follow the max separation score (no hyperparameters to tune) for all the sequences contained in the original dataset, considering the CLEAN predictions with a confidence score *>* 0.5 and not categorized as enzymes in the PDB database before. Applying these filters result in 448,599 entries when considering only hydrolases (127,111 unique PDB IDs, *≃* 77% from the original training dataset, so assuming a total of *≃* 23% of PDBs removed from the original dataset), and 280,689 entries when filtering across all enzyme classes (70,343 unique PDB IDs, *≃* 43% from the original training dataset, assuming a total of *≃*57% of PDBs removed from the original dataset). Specifically, 11,159 unique PDB IDs are filtered out in addition to the PDB annotation through CLEAN, with 4,818 of them predicted as hydrolases. In total, four new ProteinMPNN models have been re-trained. Training curves and evaluation details are provided for these models. In all cases we run the ProteinMPNN retraining with the flags num neighbors=48, backbone noise=0.20, num epochs=120 saving the last models at the best epoch (lowest validation loss). Such models were then evaluated in a range of sampling temperatures from 0.1 to 1.0, taking into account the average sequence recovery and the average sequence perplexity. Once evaluated, their respective weights have been used for the generation of *β*-lactamases and Ras protein families, with default parameters and covering minimum and maximum sampling temperatures as before in the case of the pre-trained ProteinMPNN.

### Caliby Ensemble-Conditioned Protein Sequence Design

Alternatively to ProteinMPNN, we use Caliby (18), a Potts model-based sequence design method capable of conditioning on an ensemble of structures. At the time of submission, only the initial public release of Caliby was available (by November 2025) including a [later fix](https://github.com/ProteinDesignLab/caliby/blob/main/examples/scripts/clean_pdbs.sh) to clean the inputs PDBs before running the ensemble generation. Default parameters were considered for both the ensembles generation (32 conformers per target = 1 ensemble) and sequence design ensemble-conditioned (30 samples per target, conditioned on its previous single ensemble composed by 32 conformers). Caliby provides the global energy (U) of the designed sequences computed by the Potts model. Energy values are used as a scoring method to select the best designs (lower energy) per target for further analyses on the main examples exposed. In any case the lowest-energy designs correspond to the highest-quality structural models, and the correlation between pLDDT and U was notably weak (not shown).

### Structure Prediction

Protein structure of each sequence generated by ProteinMPNN or Caliby was modelled using ESMFold (*esm*.*pretrained*.*esmfold v1*) (15) with default parameters. ESMFold generates a single structure prediction per sequence, followed by no relaxation protocol. Such structural predictions are primarily needed to quantify energetic frustration conservation (24) calculated by FrustratometeR (23). Because frustration is calculated in a coarse-grain fashion at the level of whole residues, the structural accuracy of ESMFold is considered appropriate. The ESMFold predicted pLDDT values are only considered as a scoring method to select the best designs (higher values) per target for the ProteinMPNN related analyses. AlphaFold2 (v2.3.2) (14) predictions were used in some cases to compare pLDDT score ranges with ESMFold and additional controls. Moreover, because the ProteiMPNN algorithm appears to be highly sensitive to angstrom-level structural differences, AlphaFold2 was also used to predict the structure of non-available protein targets (input structures) on the Protein Data Bank (37). The max template date was set to *2023-12-31*. As AlphaFold2 generates five structure predictions per sequence, the one predicted with higher pLDDT once relaxed (Amber protocol) was the structure used as target. Structural alignments versus native references were performed using TM-align (V.20190822) (73) to derive both TM and RMSD scores. Structural visualizations were performed by PyMOL 3.0.3 and ChimeraX 1.8.

### Conservation of Local Energetic Frustration

Conservation analysis of local energy frustration of 3D sets of protein structures were performed in all the cases by FrustraEvo (24, 29), using [FrustraEvo source code](https://github.com/proteinphysiologylab/FrustraEvo) or [FrustraEvo Docker container](https://hub.docker.com/r/proteinphysiologylab/frustraevo). The input consists of (1) a MSA in FASTA format with sequences composed solely by the standard 20 amino acids code (other characters accepted by FASTA are replaced by a gap), and (2) a set of protein structures (either experimentals or models) in PDB format corresponding to the same set of sequences contained in the MSA.

For each protein structure, local energetic frustration is computed by FrustratometeR (23) by deriving decoys, already integrated in FrustraEvo. The Electrostatics K and SeqDist parameters for the calculate frustration() function remain setted by default, being NULL and 12 respectively. NULL Electrostatics K will not take into account electrostatic interactions for calculating frustration. SeqDist (sequence separation) is used to calculate the local densities of the amino acids. A distance equal to 12 is back compatible with the one used in the original FrustratometeR algorithm (74). Alternatively, users may set SeqDist=3 which aligns more closely with findings from molecular dynamics simulations performed with AWSEM-MD (75). Most results presented in the main text are made using the default SeqDist=12 since also most published analyses are based on static protein structures. For completeness we have reproduced the main examples also with SeqDist=3, showing consistent results primarily for the residues of interest.

According to the way the decoys are constructed the analyses can be carried out using any of the three different Frustration Indices (FIs): Single Residue FI (SRFI) (measured by residue), mutational FI, and configurational FI (both measured by contacts). The SRFI aggregates the energetic description of all interactions established by a given residue when mutating its identity. The mutational FI reflects how frustration changes based on the amino acid identities at a specific pair of positions. In contrast, the configurational FI considers frustration changes not only due to amino acid identities but also due to shifts in conformational state and solvent exposure. While these indices are correlated, the mutational FI is particularly useful for identifying active or ligand-binding sites, whereas the configurational FI is more suitable for studying protein-protein interactions and conformational changes (24). For these reasons, only the SRFI was evaluated for all case studies while mutational FI was also included in some of them. Configurational FI has not been explicitly considered in these case studies. For all the FIs, 2,000 decoys are generated. The native energy is compared against the mean and standard deviation of such distribution by computing a Z-score and therefore, the FIs are expressed in standard deviation units, typically in the [-4, 4] range. Then, the native residues or contacts are classified as highly, neutrally, or minimally frustrated according to how distant the native energy is from the mean value of the energy distribution of the decoys. The configurational and mutational FIs have the following thresholds to define the different frustration states for the interactions, as proposed by Ferreiro et al. (26, 75): if FI*<*-1 then the interaction is highly frustrated. If FI *>* 0.78 then the interaction is minimally frustrated. If −1*<*FI*<*0.78 then the interaction is neutral. In the case of the SRFI thresholds for single residues, if SRFI *<*-1 then the residue is highly frustrated. If SRFI*>*0.55 then the residue is minimally frustrated. If −1*<*SRFI*<*0.55, then the interaction is neutral.

Then Sequence and Frustration Information Content, SeqIC and FrustIC respectively, are calculated using information theory concepts (Shannon information content formulas). SeqIC is calculated from aligned residues in the MSA, based on the distribution of amino acid identities. FrustIC is calculated for aligned residues in a MSA or equivalent contacts across proteins in the MSA, based on the frustration states mapped into the residues from the structures. A reference protein must be selected to define over which residues or contacts conservation is computed. The reference structure can be defined by the user or otherwise FrustraEvo selects the protein that maximizes the sequence coverage of the MSA. All columns in which the reference protein has a gap are removed from the MSA (ungapped MSA). Selected sequences were aligned using MAFFT v7.520 *–auto option* (76) for all the ProteinMPNN runs. We used sequence alignments even though this is a structure-based model, as the algorithm designs show *≃* 40% sequence recovery with respect to the input targets. This is not the case expected for Caliby, whose objective is not focused on NSR, so we use template-based MSAs following the gap pattern of the native MSA (this one also from MAFFT). For comparative consensus analyses across different hyperparameter values (e.g., sampling temperature in our examples), or model versions, a single global MSA is built including all sequences. From this global alignment, sub-MSAs are derived, each containing the specific set of sequences used in individual frustration analyses. In this way, the global MSA is partitioned into sub-MSAs while preserving positional homology across all sets. In all sub-MSAs the sequence of the native reference is always included and setted as reference structure in FrustraEvo, ensuring comparable alignment positions across analyses. Frustration mapped from structures to an MSA results in an MSFA (Multiple Sequence Frustration Alignment). Positions in the MSFA with high conservation of their frustration states will display FrustIC high values (maximum theoretical height is log2(3)=1.58, as three frustration states are described), while those with no conservation will be closer to 0 (or even negative values if the distribution of states do not follow what is expected according to the background frequencies used by the algorithm (26). We consider a position to be energetically conserved if its associated FrustIC *>* 0.5. A very detailed description of the method can be found in previous works (24, 29, 77).

### Data visualization

We have used the Multiple Sequence Frustration Alignment (MSFA) visualizations developed in previous frustration analyses to compare FrustratrometeR results across multiple protein sequences. This type of plot consists of a heatmap for which each cell contains the residues in the MSA. Each cell is colored according to its SRFI in the corresponding structures, i.e., minimally frustrated residues are colored in green, neutral in gray, and highly frustrated in red. We also considered the Consensus MSFA (29) to summarize and visually compare the FrustIC and SeqIC, from multiple MSFAs from multiple sets, just with few modifications. In this case, each cell depicts the consensus sequence of the MSA of each run and the letter size is proportional to its SeqIC value. The cells background color corresponds to the frustration value of that residue across all structures contained in the set only when FrustIC*>*0.5. The background of the cell is white when FrustIC *≤* 0.5. If no letter is included in the cell, it is a gap conserved in all the members of the set except in the reference protein.

## Supporting Information Appendix (SI)

Supplementary text, Figures S1-S31, and Tables S1-S4 are provided in a separate accompanying PDF.

## Data, Materials, and Software Availability

Data mostly generated and processed using the MareNostrum5 supercomputer at the BSC-CNS. We used the Frustratrometer R package (23) to calculate frustration on individual structures and to model energy-minimized mutants; [FrustraEvo source code](https://github.com/proteinphysiologylab/FrustraEvo) or [FrustraEvo Docker container](https://hub.docker.com/r/proteinphysiologylab/frustraevo) (24, 29) to compute the conservation of local energetic frustration patterns for all the presented family analyses. Plots were produced with ggplot2 and ggpubr R packages, others with seaborn and matplotlib Python libraries, combined with networkx to visualize the contacts network. The custom datasets used for retraining ProteinMPNN and the resulting model weights are available at [repository-weights](https://zenodo.org/records/18957473). All input/output data needed to reproduce the main results of this article as well as the intermediate analyses are available at [repository-data](https://zenodo.org/records/18922047). Related code and documentation available at [repository-code](https://github.com/miriampol2c/architectural-constraints).

## ACKNOWLEDGMENTS

We thank Christine Orengo, Nicola Bordin, Noelia Ferruz, Peter Wolynes, and Diego Ferreiro for scientific discussions, and Ignacio Sarasua (NVIDIA) for technical support. R.G.P. was supported by the Ramon y Cajal program (RYC2023-043825-I) and the MEGAFrustratEDS grant (PID2024-159128OA-I00). M.P.G. was funded by the BSC-NVIDIA HPC RD collaboration contract (15/12/2023). M.F.M. was supported by MICIU/AEI/10.13039/501100011033 (Grant PRE2022-101718) and “ESF+” (Grant CEX2021-001148-S-20-2). R.G.P. dedicates this manuscript to the memory of Amos Bairoch, Reinhard Schneider and Peer Bork for their many significant contributions to our discipline.

## AUTHOR CONTRIBUTIONS

M.P.G: Conceptualization, Formal Analysis, Investigation, Visualization, Writing-original draft, Writing-review & editing. M.F.M: Writing-review & editing. A.B: Code Provider. M.H: Writing-review. B.R: Writing-review. A.V: Conceptualization, Funding Acquisition, Supervision, Writing-review & editing. R.G.P: Conceptualization, Investigation, Supervision, Writing-original draft, Writing-review & editing.

## References

1. CB. Anfinsen, Principles that govern the folding of protein chains. Science 181, 223–230 (1973).

2. C. Chothia, AM. Lesk, The relation between the divergence of sequence and structure in proteins. The EMBO journal 5, 823–826 (1986).

3. LN. Kinch, NV. Grishin, Evolution of protein structures and functions. Curr. Opin. Struct. Biol. 12, 400–408 (2002).

4. J. Maynard Smith, Natural selection and the concept of a protein space. Nature 225, 563–564 (1970).

5. BK. Shoichet, WA. Baase, R. Kuroki, BW. Matthews, A relationship between protein stability and protein function. Proc. Natl. Acad. Sci. United States Am. 92, 452–456 (1995).

6. N. Tokuriki, F. Stricher, L. Serrano, DS. Tawfik, How Protein Stability and New Functions Trade Off. PLOS Comput. Biol. 4, e1000002 (2008).

7. OB. Ptitsyn, AV. Finkelstein, Similarities of protein topologies: evolutionary divergence, functional convergence or principles of folding? Q. Rev. Biophys. 13, 339–386 (1980).

8. AV. Finkelstein, OB. Ptitsyn, Why do globular proteins fit the limited set of foldin patterns? Prog. Biophys. Mol. Biol. 50, 171–190 (1987).

9. B. Honig, Protein folding: from the levinthal paradox to structure prediction. J. Mol. Biol. 293, 283–293 (1999).

10. MJ. Denton, CJ. Marshall, M. Legge, The Protein Folds as Platonic Forms: New Support for the Pre-Darwinian Conception of Evolution by Natural Law. J. Theor. Biol. 219, 325–342 (2002).

11. E. Noor, et al., Uniform binding and negative catalysis at the origin of enzymes. Protein Sci. : A Publ. Protein Soc. 31, e4381 (2022).

12. J. Skolnick, M. Gao, Interplay of physics and evolution in the likely origin of protein biochemical function. Proc. Natl. Acad. Sci. 110, 9344–9349 (2013).

13. I. Friedberg, A. Godzik, Functional Differentiation of Proteins: Implications for Structural Genomics. Structure 15, 405–415 (2007).

14. J. Jumper, et al., Highly accurate protein structure prediction with AlphaFold. Nature 596, 583–589 (2021).

15. Z. Lin, et al., Evolutionary-scale prediction of atomic-level protein structure with a language model. Science 379, 1123–1130 (2023).

16. B. Rost, C. Sander, Bridging the protein sequence-structure gap by structure predictions. Annu. review biophysics biomolecular structure 25, 113–136 (1996).

17. J. Dauparas, et al., Robust deep learning-based protein sequence design using ProteinMPNN. Sci. (New York, N.Y.) 378, 49–56 (2022).

18. RW. Shuai, T. Lu, S. Bhatti, P. Kouba, PS. Huang, Ensemble-conditioned protein sequence design with Caliby (2025) ISSN: 2692-8205 Pages: 2025.09.30.679633 Section: New Results.

19. B. Rost, Protein structures sustain evolutionary drift. Fold. Des. 2, S19–S24 (1997).

20. P. Tian, RB. Best, How Many Protein Sequences Fold to a Given Structure? A Coevolutionary Analysis. Biophys. J. 113, 1719–1730 (2017).

21. H. Frauenfelder, SG. Sligar, PG. Wolynes, The energy landscapes and motions of proteins. Sci. (New York, N.Y.) 254, 1598–1603 (1991).

22. JD. Bryngelson, PG. Wolynes, Spin glasses and the statistical mechanics of protein folding. Proc. Natl. Acad. sciences 84, 7524–7528 (1987).

23. AO. Rausch, et al., FrustratometeR: an R-package to compute local frustration in protein structures, point mutants and MD simulations. Bioinforma. (Oxford, England) 37, 3038–3040 (2021).

24. RG Parra, et al., Frustraevo: a web server to localize and quantify the conservation of local energetic frustration in protein families. Nucleic Acids Res. 52, W233–W237 (2024).

25. N. Tokuriki, DS. Tawfik, Stability effects of mutations and protein evolvability. Curr. Opin. Struct. Biol. 19, 596–604 (2009).

26. DU. Ferreiro, JA. Hegler, EA. Komives, PG. Wolynes, Localizing frustration in native proteins and protein assemblies. Proc. Natl. Acad. Sci. 104, 19819–19824 (2007).

27. DU. Ferreiro, JA. Hegler, EA. Komives, PG. Wolynes, On the role of frustration in the energy landscapes of allosteric proteins. Proc. Natl. Acad. Sci. 108, 3499–3503 (2011).

28. MI. Freiberger, AB. Guzovsky, PG. Wolynes, RG. Parra, DU. Ferreiro, Local frustration around enzyme active sites. Proc. Natl. Acad. Sci. 116, 4037–4043 (2019).

29. MI. Freiberger, et al., Local energetic frustration conservation in protein families and superfamilies. Nat. Commun. 14, 8379 (2023).

30. SJ. Gould, RC. Lewontin, The spandrels of San Marco and the Panglossian paradigm: a critique of the adaptationist programme. Proc. Royal Soc. London. Ser. B, Biol. Sci. 205, 581–598 (1979).

31. SJ. Gould, The exaptive excellence of spandrels as a term and prototype. Proc. national academy sciences 94, 10750–10755 (1997).

32. M. Manhart, AV. Morozov, Protein folding and binding can emerge as evolutionary spandrels through structural coupling. Proc. Natl. Acad. Sci. 112, 1797–1802 (2015).

33. SJ. Gould, ES. Vrba, Exaptation—a Missing Term in the Science of Form. Paleobiology 8, 4–15 (1982).

34. ML. Salverda, JAG. De Visser, M. Barlow, Natural evolution of TEM-1 -lactamase: experimental reconstruction and clinical relevance. FEMS Microbiol. Rev. 34, 1015–1036 (2010).

35. JR. Knox, PC Moews, -Lactamase of Bacillus licheniformis 749/C. J. Mol. Biol. 220, 435–455 (1991).

36. AJ. Ribeiro, JD. Tyzack, N. Borkakoti, GL. Holliday, JM. Thornton, A global analysis of function and conservation of catalytic residues in enzymes. J. Biol. Chem. 295, 314–324 (2020).

37. SK. Burley, et al., Updated resources for exploring experimentally-determined pdb structures and computed structure models at the rcsb protein data bank. Nucleic acids research 53, D564–D574 (2025).

38. T. Yu, et al., Enzyme function prediction using contrastive learning. Sci. (New York, N.Y.) 379, 1358–1363 (2023).

39. F. Jacob, B. Joris, S. Lepage, J. Dusart, JM. Frère, Role of the conserved amino acids of the ‘SDN’ loop (Ser130, Asp131 and Asn132) in a class A beta-lactamase studied by site-directed mutagenesis. Biochem. J. 271, 399–406 (1990).

40. J. Echave, M. Carpentier, On the variation of structural divergence among residues in enzyme evolution. Proteins: Struct. Funct. Bioinforma. 94, 1030–1046 (2026).

41. VG. Contessoto, et al., Analyzing the effect of homogeneous frustration in protein folding. Proteins 81, 1727–1737 (2013).

42. B. Kuhlman, et al., Design of a novel globular protein fold with atomic-level accuracy. Sci. (New York, N.Y.) 302, 1364–1368 (2003).

43. Z. Zhang, HS. Chan, Native Topology of the Designed Protein Top7 is Not Conducive to Cooperative Folding. Biophys. J. 96, L25–L27 (2009).

44. BI. Dahiyat, SL. Mayo, Probing the role of packing specificity in protein design. Proc. Natl. Acad. Sci. 94, 10172–10177 (1997).

45. ST. Walsh, H. Cheng, JW. Bryson, H. Roder, WF. DeGrado, Solution structure and dynamics of a de novo designed three-helix bundle protein. Proc. Natl. Acad. Sci. 96, 5486–5491 (1999).

46. W. Kabsch, C. Sander, Dictionary of protein secondary structure: pattern recognition of hydrogen-bonded and geometrical features. Biopolym. Orig. Res. on Biomol. 22, 2577–2637 (1983).

47. RE. Hackett, et al., Investigating enzyme function by geometric matching of catalytic motifs. bioRxiv pp. 2026–02 (2026).

48. P. Maietta, et al., Firedb: a compendium of biological and pharmacologically relevant ligands. Nucleic acids research 42, D267–D272 (2014).

49. VP. Waman, et al., CATH 2024: CATH-AlphaFlow Doubles the Number of Structures in CATH and Reveals Nearly 200 New Folds. J. Mol. Biol. 436, 168551 (2024).

50. P. Laurino, et al., An ancient fingerprint indicates the common ancestry of rossmann-fold enzymes utilizing different ribose-based cofactors. PLoS biology 14, e1002396 (2016).

51. RG. Parra, R. Espada, N. Verstraete, DU. Ferreiro, Structural and energetic characterization of the ankyrin repeat protein family. PLoS computational biology 11, e1004659 (2015).

52. KH. Sumida, et al., Improving Protein Expression, Stability, and Function with ProteinMPNN. J. Am. Chem. Soc. 146, 2054–2061 (2024).

53. HH. Truong, BL. Kim, NP. Schafer, PG. Wolynes, Funneling and frustration in the energy landscapes of some designed and simplified proteins. The J. Chem. Phys. 139, 121908 (2013).

54. V. Alva, J Söding, AN Lupas, A vocabulary of ancient peptides at the origin of folded proteins. elife 4, e09410 (2015).

55. ML. Romero Romero, et al., Simple yet functional phosphate-loop proteins. Proc. Natl. Acad. Sci. 115, E11943–E11950 (2018).

56. P. Vyas, S. Malitsky, M. Itkin, DS. Tawfik, On the origins of enzymes: phosphate-binding polypeptides mediate phosphoryl transfer to synthesize adenosine triphosphate. J. Am. Chem. Soc. 145, 8344–8354 (2023).

57. MF. Aziz, F. Mughal, G. Caetano-Anollés, Tracing the birth of structural domains from loops during protein evolution. Sci. reports 13, 14688 (2023).

58. GL. Holliday, JB. Mitchell, JM. Thornton, Understanding the functional roles of amino acid residues in enzyme catalysis. J. Mol. Biol. 390, 560–577 (2009).

59. Y. Bromberg, et al., Quantifying structural relationships of metal-binding sites suggests origins of biological electron transfer. Sci. advances 8, eabj3984 (2022).

60. M. Makarov, et al., Early selection of the amino acid alphabet was adaptively shaped by biophysical constraints of foldability. J. Am. Chem. Soc. 145, 5320–5329 (2023).

61. K. Seya, ALR. Brownless, SC. Kamerlin, LM. Longo, The borderlands of foldability: lessons from simplified proteins. Trends Chem. (2026).

62. G. Caetano-Anollés, HS. Kim, JE. Mittenthal, The origin of modern metabolic networks inferred from phylogenomic analysis of protein architecture. Proc. Natl. Acad. Sci. 104, 9358–9363 (2007).

63. G. Caetano-Anollés, M. Wang, D. Caetano-Anollés, JE. Mittenthal, The origin, evolution and structure of the protein world. Biochem. J. 417, 621–637 (2009).

64. SA. Bukhari, G. Caetano-Anollés, Origin and evolution of protein fold designs inferred from phylogenomic analysis of cath domain structures in proteomes. PLoS computational biology 9, e1003009 (2013).

65. A. Goncearenco, IN. Berezovsky, Prototypes of elementary functional loops unravel evolutionary connections between protein functions. Bioinformatics 26, i497–i503 (2010).

66. T. Corlett, HB. Smith, E. Smith, JE. Goldford, LM. Longo, The history of enzyme evolution embedded in metabolism. Proc. Natl. Acad. Sci. 123, e2609531123 (2026).

67. OB. Ptitsyn, AV. Finkelstein, Similarities of protein topologies: evolutionary divergence, functional convergence or principles of folding? Q. Rev. Biophys. 13, 339–386 (1980).

68. C. Sander, R. Schneider, Database of homology-derived protein structures and the structural meaning of sequence alignment. Proteins 9, 56–68 (1991).

69. L. Holm, C. Sander, Mapping the protein universe. Sci. (New York, N.Y.) 273, 595–603 (1996).

70. CA. Orengo, DT. Jones, JM. Thornton, Protein superfamilles and domain superfolds. Nature 372, 631–634 (1994).

71. RA. Jensen, Enzyme recruitment in evolution of new function. Annu. review microbiology 30, 409–425 (1976).

72. C. Kocher, KA. Dill, Origins of life: first came evolutionary dynamics. QRB discovery 4, e4 (2023).

73. Y. Zhang, J. Skolnick, TM-align: a protein structure alignment algorithm based on the TM-score. Nucleic Acids Res. 33, 2302–2309 (2005).

74. M. Jenik, et al., Protein frustratometer: a tool to localize energetic frustration in protein molecules. Nucleic Acids Res. 40, W348–351 (2012).

75. RG. Parra, et al., Protein Frustratometer 2: a tool to localize energetic frustration in protein molecules, now with electrostatics. Nucleic Acids Res. 44, W356–360 (2016).

76. K. Katoh, K. Misawa, Ki. Kuma, T. Miyata, MAFFT: a novel method for rapid multiple sequence alignment based on fast Fourier transform. Nucleic Acids Res. 30, 3059–3066 (2002).

77. AB. Guzovsky, NP. Schafer, PG. Wolynes, DU. Ferreiro, Localization of Energetic Frustration in Proteins. Methods Mol. Biol. (Clifton, N.J.) 2376, 387–398 (2022).

