## Supplementary Material for "Architectonic Spandrels in the Origin of Enzymatic Function"

##### This PDF file includes:

Supporting text
Figs. S1 to S31
Tables S1 to S4
SI References

### Supporting Information Text

#### Supplementary Notes

##### 1. Mutated Functional Sites in the $\beta$-lactamases Fold Revert to Preferred, Constrained Identities

We observe that the functional residues in the $\beta$-lactamases fold resist energetic minimization when using native targets (Fig. 2). We further explore the reason for the presence of high frustration at the catalytic residues introducing single and multi-point in silico mutations to change their identities in the target structures that we used to reverse fold the sequences by ProteinMPNN. We started by analysing frustration changes upon mutations on the $\beta$-lactamase from *Bacillus Licheniformis* (PDB ID: 4blm-A). We used the FrustratometeR tool (1) to evaluate how frustration would be affected by all possible amino acid variants in a given position of the structure and selected for each catalytic residue the one that minimises local frustration. In most cases, mutations to valines result in the largest frustration decrease (Fig.S6). Then, we generated single mutants for all catalytic residues to valines (S70V, K73V, S130V, E166V, K234V, A237V) as well as the multi-point variant containing all the six mutations at the same time (*AlltoVal*). The structures of all variants were predicted with AlphaFold2 and used as ProteinMPNN targets to reverse fold new sets using the Single Target approach. Surprisingly, some identities present in the native 4blm-A structure (S70, K73, S130 and K234) were retrieved in the designs, instead of the valines that were introduced as frustration-minimizing mutations. When E166V is introduced as a point mutation, the position gets remodeled to phenylalanine with no frustration conservation. Interestingly, the only case when a valine is obtained as the most common amino acid across ProteinMPNN designs, although not fully conserved in sequence and frustration, is for the A237V case. When all catalytic residues are mutated to valines (*AlltoVal*) the original S70, K73, S130 and K234 are retrieved while F166 and V237 are introduced as conserved and minimally frustrated residues showing some sort of mutual stabilizing effect between these two positions (Fig.S7A). At the contacts level we observe that the native family contains 56 highly frustrated and conserved interactions, 30 specifically involving the catalytic residues (Fig.S8) and the other 26 in more distant regions. Following the same protocol, we introduced mutations to valines in all catalytic residues of all the $\beta$-lactamases family members. Such a mutated family, compared to the native one (Fig.S7B), has a network of minimally frustrated interactions around their mutated catalytic residues that now contains valines (Fig.S7C). Despite minimising the catalytic site, the mutated family retains the other 26 interactions highly frustrated and conserved common to the native family and proposes 24 new interactions redistributed in such a way that the total amount of highly frustrated interactions remains at similar levels in both families (n contacts 56 vs. 50) (Fig.S8). Nevertheless, when using these mutated structures to reverse fold sequences with ProteinMPNN using the Family Target approach, the same trend was observed. Instead of introducing valines in the proposed designs as expected, ProteinMPNN introduces the same amino acid identities that are present in the native structures, bringing back high frustration at both the single residue level (Fig.S7A) and at the contacts level (Fig.S7D) surrounding the catalytic site. To rule out that recovering of the native catalytic identities instead of the introduced valines is due to a possible artefact related to the predicted structure models, we repeated the same analysis on 3 mutant $\beta$-lactamases with experimentally resolved structures where a catalytic residue has been replaced by an alanine (PDB IDs: 1mbl-E166A, 5hap-S70A, 7qqc-K73A). Analysis of changes in frustration upon mutation with the FrustratometeR show that, in all cases, these mutations to an alanine lead to a predicted decrease in frustration around the catalytic residue (Fig.S9A). As observed with the AlphaFold2 models, even when using experimentally resolved structures of mutants as input, the original native identities of the catalytic residues are restored in the ProteinMPNN designs. For 7qqc, the K73A mutation reduces the conservation of the highly frustrated state, though K73 remains one of the proposed identities (Fig.S9D). However, we observed an exception with 1mbl-E66A. E166 is not recovered by ProteinMPNN but K166 or F166 are introduced instead, which aligns with the results from the in silico 4blm-A E166V mutants (Fig.S7A). This is consistent with previous analysis where the same catalytic residue is varied in ProteinMPNN designs when using the reference native $\beta$-lactamase as input (Fig. 2B). Simulating these experimental mutants but in silico produces a comparable outcome (Fig.S9E).

##### 2. Functional residues in the $\alpha$-globin fold are energetically minimized

We also analysed the $\alpha$-globins family, that is part of the Hemoglobin molecule where protein-protein interfaces in its tetrameric structure are relevant for function. In a recent study, we described the similarities and differences in frustration between $\alpha$ and $\beta$ globins. We found that mostly minimally frustrated positions, i.e., the stability anchors of the fold, were conserved in both families. In contrast, there was no overlap between the set of conserved and highly frustrated residues between both families. Most of these positions were found to be related to protein-protein interaction (PPI) sites between the Hemoglobin subunits or between $\alpha$-globin and the AHSP chaperone as well as residues that participate in salt bridges (2). The energetic diversity across different $\alpha$-globins family members (non-redundant sequences, n=21) calculated by FrustraEvo (3) shows that 69% of all positions in the MSFA (n=96) are energetically conserved, from which 26% (n=36) are minimally frustrated, mostly linked to stability and foldability; 34% (n=48) are neutral, and 9% (n=12) are highly frustrated, mostly correlated with PPIs. The remaining 31% (n=44) of positions are not energetically conserved (Fig.S18A, S21D). Fig.S16A contains the sub MSFA containing the 12 conserved and highly frustrated positions (K7, K11, E27, E30, K40, Y42, Q54, K99, S124, D126, K127 and Y140) and the four involved in salt bridges or chaperone interaction (K7/D74, E27/H112, E30/H50, K99/R141). We explored how frustration varies across reverse folded ProteinMPNN sequences (4) in the $\alpha$-globin family using both the Single Target (using Human $\alpha$-globin, PDB ID: 2dn1-A, Fig.S16B) and the Family Target (Fig.S16C) approaches. Fig.S16D shows a full-sequence consensus MSFA that facilitates the comparison between the native $\alpha$-globins family with designs from both approaches. We observe that residues that are conserved and minimally frustrated or neutral tend to retain their identities after reverse folding. In contrast, positions that are conserved and highly frustrated in native $\alpha$-globins are remodeled by ProteinMPNN in such a way that the associated high frustration is decreased. Q54 and Y140 are exceptions, being the only two positions among the 12 positions that are conserved and highly frustrated in native $\alpha$-globins that are recovered by ProteinMPNN when using the Single Target approach. When applying the Family Target approach, only Q54 is recovered as conserved and highly frustrated. Interestingly, we have not been able to find functional evidence for that position. However, it is worth noting that although local frustration is minimised, no design generated by ProteinMPNN (T=0.1) is frustration free (Fig.S18B-C), retaining on average between $\simeq$ 1% (if Family Target, Fig.S21D) and $\simeq$ 4% (if Single Target, Fig.S20D) of frustration (vs. 10% on average in native and globular proteins (5). Notably, the consensus MSFA also displays a few positions where ProteinMPNN introduces conserved and highly frustrated identities in the predicted designs compared to the natives ones, e.g. A53, N115 (Single Target) or P119 (both Single and Family Target) (Fig.S16D). When designing at higher temperature (T=1) (Fig.S19B-C), there is a decrease of about $\simeq$ 30% in energetically conserved positions in Single Target (Fig.S20D), comparing the minimum and the maximum T) and Family Target (Fig.S21D), comparing the native and the maximum T). In Single Target, conserved and minimally frustrated positions decrease by 11% while conserved and neutral frustrated positions decrease by 18% (Fig.S20D). In Family Target, 6% and 16% respectively (Fig.S21D). The ProteinMPNN designs contain in both approaches highly frustrated residues when sampling at such temperature, but they are always fewer than in the native proteins and are not consistently located in alignable positions of the MSAs (Fig.S19, S19). None of the originally highly frustrated and conserved positions both in the Single Target ($\simeq$ 4%) and Family Target ($\simeq$ 9%) remains in such state at the maximum T (Fig.S20, S21). In other words, a systematic conservation of frustrated residues is not observed in non-enzymatic cases such as the $\alpha$-globins fold. As for $\beta$-lactamases, we also applied Caliby (6) to reverse-fold sequences from the $\alpha$-globin native family and computed frustration using the parametrization that better correlates with folding simulations (*Suppl. notes* 4, 5). In contrast to $\beta$-lactamases, we observed that high local frustration in the $\alpha$-globin family is not maintained at any residue across the tested strategies, suggesting that these residues more likely reflect foldability or kinetic constraints rather than architectural constraints (*Table* S3).

#### 3. Persistent Frustration in the Ras Fold

To investigate if the observations for $\beta$-lactamases hold true for other enzymatic families, we studied deeper the Ras family (n=35). It is also a class of hydrolase (E.C 3.6.5.2) involved in the GTP hydrolysis, where the GTP-binding site consists of 5 motifs (G1-G5). In a previous work we reported that from the seven residues that interact with the nucleotide substrate, K117 (from KRAS numbering, located within G4) is conserved and highly frustrated at the family level suggesting a strong energetic conflict in that site that is important for function (2). We used the Family Target approach on the Ras family using the original ProteinMPNN as well as the PDB-AllEnzymes and the CLEAN-AllEnzymes retrained models at minimum and maximum temperatures. The SRFI results show that for all models and both temperatures the K117 identity is recovered. In some cases, K117 appears as conserved and highly frustrated and in other cases it appears neutral or not energetically conserved (Fig.S16A,B). Despite these fluctuations, the conservation percentages by state and ProteinMPNN version at the minimum temperature closely resemble those of the native Ras family (Fig.S15C), as observed previously in $\beta$-lactamases (Fig.S12C). Previous MutFI results from the native Ras family also showed a network of highly conserved and frustrated interactions mainly involving residues that interact with the substrate, including the K117 (2). When comparing such networks with the ones resulting from ProteinMPNN designs (Fig.S16C-E), we observe that in most cases some conserved and highly frustrated interactions involving K117 are retrieved. The only exception are the designs that were generated by the CLEAN-AllEnzymes model, which recovers mostly neutral interactions for this residue (Fig.S16D-E). Again, in agreement with the case of $\beta$-lactamases, frustration tends to be less conserved in models trained on smaller datasets (Fig.S12C, S15C). In contrast, K117 is not conserved as highly frustrated in Caliby designs, regardless of the frustration parameterization used, and even when no alternative amino acid identities are introduced (Fig.S30). Given the structural variability observed within the Ras family and therefore subsequent designs (Fig.S15A,B), restricting reverse folding to the domain of interest (G4) may provide a more appropriate design strategy specially with Caliby.

#### 4. Persistent Frustration across Structural Ensemble-Conditioned Sequence Design with Caliby

We hypothesize that training objectives focused on recovering native sequences, as can be the case of ProteinMPNN, may inadvertently drive models to capture non-structural signals rather than learning a truly generalizable mapping between structure and sequence. In consequence, we have tried to mitigate any possible bias on ProteinMPNN when designing static proteins containing any potential architectural constraint (see Local Frustration in the $\beta$-lactamase fold section and *Suppl. note* 5). However, to provide further evidence beyond an artefact of the ProteinMPNN architecture, we also test our main examples when running Caliby, a Potts model-based sequence design method capable of conditioning on an ensemble of structures (set of conformers) generated by Protpardelle-1c (7). We used the Family Target approach (but selecting the designs based on their optimized global energy (U score), instead of based on their pLDDT ESMFold score) using the initial release of Caliby, and measuring frustration at both SeqDist possible values in the same sets. The $\beta$-lactamases protein family results reveal a similar pattern to that described to date in all ProteinMPNN executions. Positions equivalent to K73, D131 and K234 across designs mostly retain their native identities as highly frustrated, however the high frustration state is not conserved for K234 when measuring frustration at SeqDist3 (FrustIC=0.30 at SeqDist3, FrustIC=0.54 for SeqDist=12) (Fig.S27). In 5/22 cases, Caliby proposes minimal frustrated residues such as V or A at the catalytic residue at position 234 by maintaining the typical $\beta$-lactamase fold. K234 is implicated in the initial protonation of the substrate N(1) by interaction with S130 through hydrogen bonding, so V or A substitution would likely reduce acylation efficiency and therefore the stabilization of the transition state. Further studies are required to assess the behavior of these designs as functional $\beta$-lactamases. The results also remain inconclusive for residues that we define as carriers of adaptive frustration in $\beta$-lactamases (Fig. 5) with such complementary strategies. A summary of how the different conserved and highly frustrated residues in the native family behave across design strategies can be seen in (*Table* S2). For *alpha*-globins, only P37 and P119 remain highly frustrated and conserved when measuring frustration at Caliby SeqDist=3, as mentioned below for ProteinMPNN T=0.1 (*Suppl. note* 5), unlike the same sets at SeqDist=12. Because prolines can distort $\alpha$-helices, it would be tempting to see this signal as representative of a spandrel, but its dependence on parametrization renders this interpretation tentative. Q54 is not conserved for any frustration state in the Caliby executions; in fact, it does not follow at all the background distribution, as it presents a negative FrustIC value (Fig.S28). Since no residue in the family remains clearly conserved and highly constrained under all conditions, we leave open the question of whether the folding of $\alpha$-globins would have required some kind of architectural bracket, given their geometry as monomers. A summary of how the different conserved and highly frustrated residues in the native family behave across design strategies can be seen in (*Table* S3). De novo examples reverse folded by Caliby also introduces some frustration across designs. Frustrated positions are scattered across designs in a non-conserved way with a minimum of 1 position per sequence. Only a couple of positions are conserved as highly frustrated for Top7 (PDB ID: 1qys) regardless of the value of the SeqDist parameter, surprisingly coinciding with the positions proposed by ProteinMPNN at so (Fig.S29, MSFA positions 22 and 43). This underscores that intrinsic frustration is not a universal feature of all folds, with Top7 being one that may potentially exhibit it.

#### 5. Persistent Frustration is Independent of FrustratometeR Parameterization

The value of SeqDist=12, frustration default parameter for all the runs, maintains consistency with the original FrustratometeR implementation (8) facilitating comparison with the majority of published static-structure analyses. But the value SeqDist=3 reflects shorter-range interactions favored in dynamic models such as AWSEM-MD, capturing local packing effects more typical of conformational ensembles and better correlates with observed folding MD simulations. To ensure the robustness and interpretability of our frustration analyses, we have reproduced the main ProteinMPNN examples also with SeqDist=3, showing consistent results primarily for the residues of interest. The same catalytic positions described previously and considered highly frustrated by consensus in $\beta$-lactamases (Fig.S3C, S4C) are still conserved as such (K73, K234), after both approaches at all temperatures and measured at SeqDist=3 (Fig.S5C). The non-catalytic, but perhaps functionally constrained, D131 in the SDN loop is also observed in most cases. This supports our hypothesis of the presence of both exapted (K73, K234) and non-exapted (e.g. D131) architectural constraints in this protein family. However, the inconsistency of the Caliby results based on parameterization with respect to the high frustration conservation of K234 leaves unresolved whether this residue represents an architectural constraint or a foldability requirement, similar to Q54 in $\alpha$-globins (*Suppl. note* 2, 4). In the case of $\alpha$-globins, we again observed how frustration at functional sites could be reduced in the designed sequences independently of the sampling temperature, as pointed out with SeqDist=12 (Fig.S20, S21). In fact, the S124 previously described as highly frustrated in the native family would not be so with SeqDist=3. Some prolines (P37, P119) also emerge as potential architectural constraints, being highly conserved and frustrated across lower sampling temperatures (Fig.S22) and matching the Caliby results (*Suppl. notes* 4. Those are indeed residues that would require a more comprehensive study. Lastly, de novo examples reverse folded by ProteinMPNN consistently introduced frustrated residues not present in the original designs in a similar way regardless of the value of the SeqDist parameter (Fig.S23-S26). In the absence of evolutionary history and native homology information, frustrated positions may fluctuate slightly in the designs, varying the total number of frustrated residues per sequence, but in no case we observed a frustration-free design. This somewhat supports the need for minimal frustration to build a fold even if it has not been shaped by evolution. Only Top7 and PDB ID: 1fsd Single Target runs contain highly frustrated and conserved residues that could be attributed to the topology of the folds. However, only in the case of Top7 also the Caliby results are consistent (*Suppl. note* 4), and therefore could be considered real architectural constraints. These results, comparable across parameter-specific runs, demonstrate that our conclusions are not artefacts of a particular parameter choice, also highlight the methodological continuity between traditional static and more physically detailed dynamic frameworks, while providing a bridge for researchers interested in extending frustration toward simulation-based contexts.

#### Data

##### $\beta$-lactamases data/workflow.

**Natives.** To analyse the $\beta$-lactamases native family we took the same dataset previously analysed in terms of energetic conflicts (Freiberger et al., 2019), containing Class A $\beta$-lactamases structures whose catalytic residues are annotated with experimental evidence at the CSA database (version 2.0, by 2018) (n=31). These PDB files were also considered to test protein sequence design by ProteinMPNN and Caliby.

**ProteinMPNN generation.** For each PDB structure (targets) and sampling temperature we generated 20 sequences, using the pre-trained ProteinMPNN but also the four re-trained ProteinMPNN versions. A total of 6200 designs were generated using the pretrained ProteinMPNN (31 targets x 20 designs x 10 temperatures). Considering the four new retrained versions of ProteinMPNN just covering the minimum and the maximum sampling temperatures, 7440 designs were additionally generated

(31 targets x 30 designs x 2 temperatures x 4 retrained ProteinMPNN versions). We folded all the designed sequences with ESMFold and analysed frustration conservation on selected designs following the two approaches described as Single Target and Family Target.

**Caliby generation.** For each target we generated 30 sequences, using the initial Caliby release and default parameters. A total of 930 designs (31 targets x 30 designs) were generated. However, the sequences generated for 9 targets [1n9bA, 1ylpA, 1ylzA, 2qpnA, 2xr0A, 3g30A, 3gmwA, 3lezA, 3ni9A] were discarded due to missmatches in the sequence length of the designs with respect to the native sequence. We folded all the designed sequences with ESMFold and analysed frustration conservation on the selected designs following only the Family Target approach. In a first approach that we refer to as Single Target we took all the designs generated per reference target and sampling temperature and applied on top of them the same frustration analysis, but to measure frustration conservation within the designs proposed by ProteinMPNN for the same given target. The Single Target approach was only applied on the reference structure (PDB ID: 4blm chain A (9)). Each of these frustration analyses contain 20 or 30 designed sequences of 4blm-A but also the native target 4blm-A (total n=21 or n=31). In the second approach that we refer to as Family Target we took only the best design generated per target and temperature instead of all of them, but considering the full $\beta$-lactamases dataset. To select for each target the best design, we chose the one that maximises the pLDDT score when predicting the structures by ESMFold (if ProteinMPNN design runs) or the one that minimizes the global energy U predicted by a Potts model (if Caliby design runs). In this case, the frustration analyses contain 32 designed ensembles, one per each $\beta$-lactamase native family member, but also the native target 4blm-A.

**In silico $\beta$-lactamases mutants.** To evaluate what happens when we mutate the identity of native catalytic sites (S70, K73, S130, E166, K234, A237) by all the other 19 amino acids identities, we mutated in silico the reference $\beta$-lactamase (4blm-A) at all the given positions by using the mutate_res() function with Modeller mode from FrustratometeR R-package (1). We chose an alternative identity that minimises frustration at those positions, building single point mutants for each catalytic residue (only for the reference structure 4blm-A) and multi point mutants including all of them (for the entire family, n=31). Then, these in silico mutants were folded by AlphaFold2 to be used as targets for ProteinMPNN as the case of native targets, following the Single Target or Family Target approach accordingly (20 designs per target in any case). Additionally, the D131V single mutant was also tested following the Single Target approach on top of 4blm-A.

**Experimental $\beta$-lactamases mutants.** To compare also the effect of introducing naturally occurring mutations in the Protein-MPNN algorithm, we also did a manual search in the Protein Data Bank (10), selecting three cases to illustrate: (1) considering our native reference 4blm-A vs. 1mbl-A (9) (E166Ala mutant), (2) considering the native OXA-48 $\beta$-lactamase - 7jhq ((11) vs. 5hap (12) (S70A mutant) and (3) considering the native CTX-M-15 extended-spectrum $\beta$-lactamase - 4hbt (13) vs. 7qqc (14) (K73A mutant). Such mutants were used as targets for ProteinMPNN as the case of native targets, following the Single Target approach (20 designs per target).

**Ras data/workflow.**

**Natives.** To analyse the subfamily RAS we took the same grouped dataset previously analysed by FrustraEvo (2), where candidate models were generated with AlphaFold2 with a mean pLDDT score of 83.5 (n=35). These models were considered to test protein sequence design by ProteinMPNN and Caliby.

**ProteinMPNN generation.** For each model (the targets), at minimum and maximum sampling temperatures we generated 20 sequence sequences, using the pre-trained ProteinMPNN but also the two re-trained models that include the filtering of all enzyme classes. In consequence, a total of 4200 designs were generated (35 targets x 20 designs x 2 temperatures x 3 ProteinMPNN versions). We folded the designed sequences with ESMFold and analysed frustration conservation on the selected designs following only the Family Target approach.

**Caliby generation.** For each native target we generated 30 sequences, using the initial Caliby release and default parameters. A total of 1050 designs (35 targets x 30 designs) were generated. We folded all the designed sequences with ESMFold and analysed frustration conservation on the selected designs following only the Family Target approach.

**$\alpha$-globins data/workflow.**

**Natives.** To analyse the $\alpha$-globin natural family we took the same dataset previously analysed by FrustraEvo (Freiberger et al., 2023) containing all non-redundant mammalian hemoglobins (n=21) present in the PDB database (by April 2022), taking the chains A individually. These splitted PDB files were considered to test protein sequence design by ProteinMPNN and Caliby.

**ProteinMPNN generation.** For each native target and sampling temperature we generated 20 sequences, using the pre-trained ProteinMPNN. A total of 4200 designs were generated (21 targets x 20 designs x 10 temperatures x pretrained ProteinMPNN). We folded the designed sequences with ESMFold and analysed frustration conservation on the selected designs following the two approaches described above for $\beta$-lactamases. The Single Target approach was only applied on the Human reference $\alpha$-globin (PDB ID: 2dn1 chain A (15). Each Single Target frustration analysis per sampling temperature contain the 20 designed sequences of 2dn1-A but also the native target 2dn1-A (total n=21), while for the Family Target approach each frustration analysis contain 21 designed sequences, one per each $\alpha$-globin native family member, but also the native reference 2dn1-A (total n=22).

**Caliby generation.** For each native target we generated 30 sequences, using the initial Caliby release and default parameters. A total of 630 designs (21 targets x 30 designs) were generated. We folded all the designed sequences with ESMFold and analysed frustration conservation on the selected designs following only the Family Target approach.

##### *De novo* proteins data/workflow.

**Natives.** To evaluate frustration on de novo proteins, we took three prime examples from the Protein Data Bank (10): 1QYS (Top7 (16)), 1FSD (17) and 2A3D (18). These PDB files were considered to test protein sequence design by ProteinMPNN and Caliby.

**ProteinMPNN generation.** For each structure, at a default sampling temperature (T=0.1), we generated at least 50 sequences using the pre-trained ProteinMPNN. Designed sequences were folded with ESMFold, and frustration conservation was analyzed using only the Single Target approach. For 2A3D, we selected the 50 designs with the highest TM-score (as a more sensitive measure of global structural similarity) from a set of 200 generated sequences. For 1QYS and 1FSD, a single run of 50 designs was performed. Despite these controls, success thresholds varied across de novo proteins. For 1QYS, successful designs required an RMSD $\leq$ 0.91 Å and TM-score $\geq$ 0.94. For 1FSD, the thresholds were RMSD $\leq$ 2.38 Å and TM-score $\geq$ 0.47. For 2A3D, success was defined as RMSD $\leq$ 2.47 Å and TM-score $\geq$ 0.70, all measured against the corresponding reference structure.

**Caliby generation.** For each target we generated 30 sequences, using the initial Caliby release and default parameters. A total of 150 designs (3 targets x 30 designs) were generated. We folded all the designed sequences with ESMFold and analysed frustration conservation following only the Single Target approach.

##### FunFams data/workflow.

**Natives.** To evaluate frustration across enzymatic functional families, we used a curated dataset provided by the CATH authors, comprising 740 FunFams with 20-200 members each (19). These families comprised both sequences with experimentally determined structures and sequences with available AlphaFold structures. Each FunFam contained at least one experimentally characterized sequence, used as the native reference. Complete FunFams were initially considered to evaluate sequence design using ProteinMPNN and Caliby. To ensure compatibility across structural algorithms and consistency with functional annotations, only 488 FunFams were retained for downstream frustration analyses, comprising a total of 82,075 MSA positions per configuration. Because inverse-folding models can assign different amino acid identities to the same structural position across family members and our final design selection is not based explicitly on sequence recovery, SeqIC was not necessarily available or comparable across all MSA positions of all configurations. Therefore, alignment positions with zero or missing SeqIC, as reported by the sequence-logo analysis, were excluded from subsequent consensus and intersection analyses. This filtering resulted in 79,338 MSA positions for the native set, 79,672 for the ProteinMPNN 0.1 set, 79,080 for the ProteinMPNN 1.0 set, and 79,626 for the Caliby 0.01 set.

**ProteinMPNN generation.** For each FunFam, we generated 30 sequences per member at two sampling temperatures (T = 0.1 and T = 1.0) using the pre-trained ProteinMPNN model, resulting in a total of 4,429,260 designs (73,821 targets $\times$ 30 designs $\times$ 2 temperatures). The designed sequences were folded using ESMFold, and frustration conservation (SeqDist=3) was computed on the subset of designs with the highest pLDDT scores, following the Family Target approach. In this framework, each frustration conservation analysis includes all selected designed sequences corresponding to the FunFam members (n), together with the native reference (total n + 1).

**Caliby generation.** We generated 30 sequences per member at the default sampling temperature (T = 0.01) using the initial release and default parameters, yielding a total of 2,214,630 designs. These sequences were also folded with ESMFold, and frustration conservation (SeqDist=3) was computed on the subset minimizing global energy (U), using the same Family Target approach as for ProteinMPNN.

**Functional annotation.** Frustration conservation results were linked to functional annotations by mapping residues onto the native reference of each FunFam. Annotations were integrated from both EnzyMM (20) and FireDB (21). EnzyMM accepts any structure as input, but is limited to a small templates library to date (February 2026). To ensure robustness and avoid uncertain mappings, only annotations corresponding to the native references were considered. For FireDB, we queried the SQL database (last updated February 2024) to extract annotations mapped to native references, as it contains only experimentally derived structures. We considered both catalytic and ligand-binding residues specific to each query, as well as consensus ligand-binding sites derived from cluster-level information. These consensus sites correspond to "master sequence binding sites" (MBS), which aggregate binding information across all sequences within a cluster. Consensus ligands were included only if they were present in at least half of the proteins in the cluster, regardless of cluster size. Finally, per-residue functional annotations were consolidated across all sources (EnzyMM, FireDB-CSA, FireDB-CTT target-specific, and FireDB-CCT cluster-wide). A residue was considered functionally supported if at least one source provided evidence.

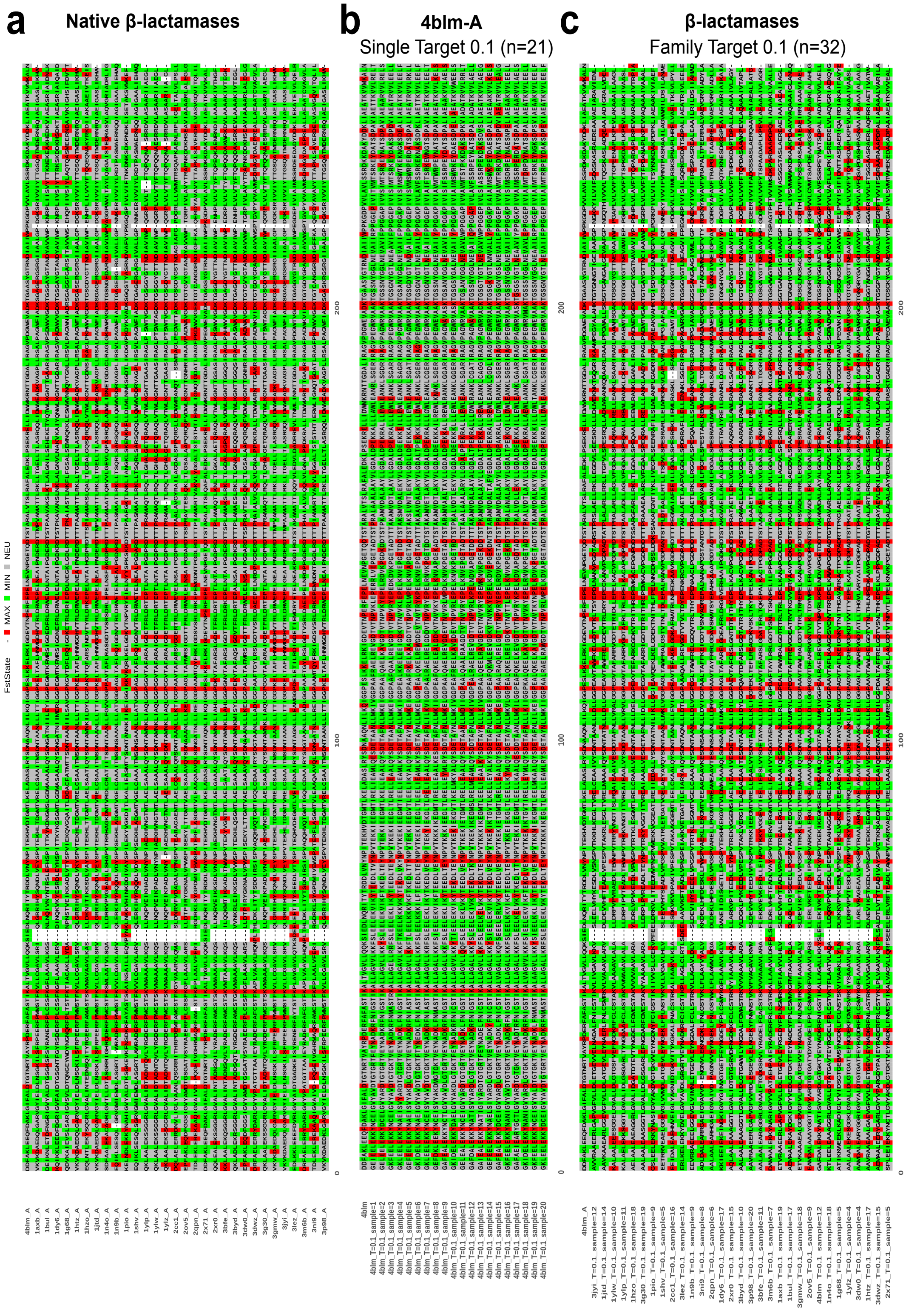


**Fig. S1.** MSFA Comparison of Native and ProteinMPNN $\beta$-lactamases (T=0.1). Full Multiple Sequence Frustration Alignment (MSFA) corresponding to the SRFI results when (a) using $\beta$-lactamases native structures (n=31), (b) the 4blm-A Single Target approach (using 20 designed protein ensembles and the reference 4blm-A, n=21) and C) the Family Target approach (using 1 ensemble per native $\beta$-lactamase and the reference 4blm-A, n=32) generated at the minimum sampling temperature (T=0.1). The minimum temperature refers to the minimum sampling temperature of the algorithm tested in our analyses, coinciding with the default. The ensembles chosen by native $\beta$-lactamase in (c) are those that maximize the ESMFold pLDDT score for the target $\beta$-lactamase in each case, among all those designed. Positions are colored according to its frustration state, except for the gaps, when SeqDist frustration remains setted by default to SeqDist=12.

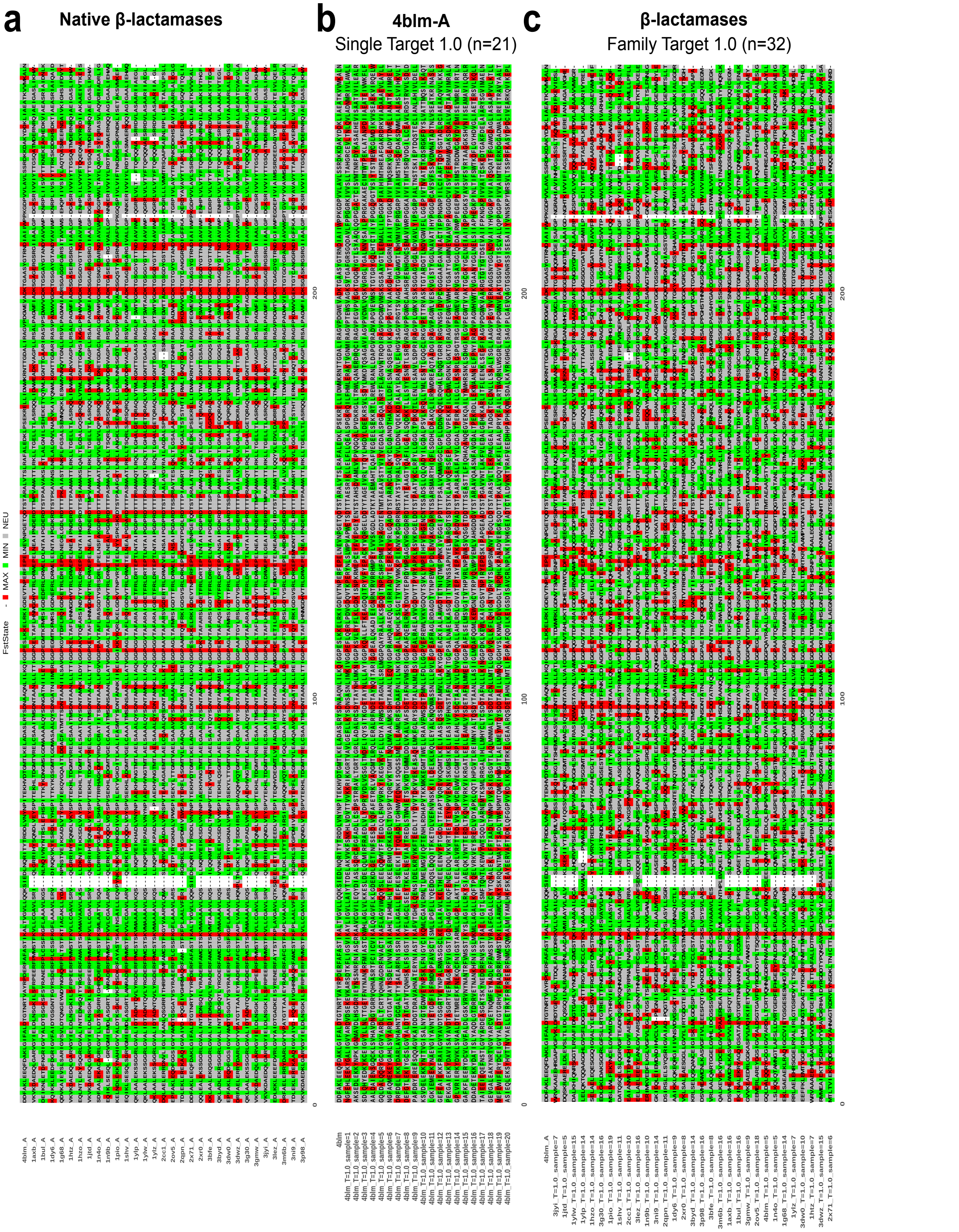


**Fig. S2.** MSFA Comparison of Native and ProteinMPNN $\beta$-lactamases (T=1.0). Full Multiple Sequence Frustration Alignment (MSFA) corresponding to the SRFI results when (a) using $\beta$-lactamases native structures (n=31), (b) the 4blm-A Single Target approach (using 20 designed protein ensembles and the reference 4blm-A, n=21) and c) the Family Target approach (using 1 ensemble per native $\beta$-lactamase and the reference 4blm-A, n=32) generated at the maximum sampling temperature (T=1.0). The maximum temperature refers to the maximum sampling temperature of the algorithm tested in our analyses. The ensembles chosen by native $\beta$-lactamase in (c) are those that maximize the ESMFold pLDDT score for the target $\beta$-lactamase in each case, among all those designed. Positions are colored according to its frustration state, except for the gaps, when SeqDist frustration remains settled by default to SeqDist=12.

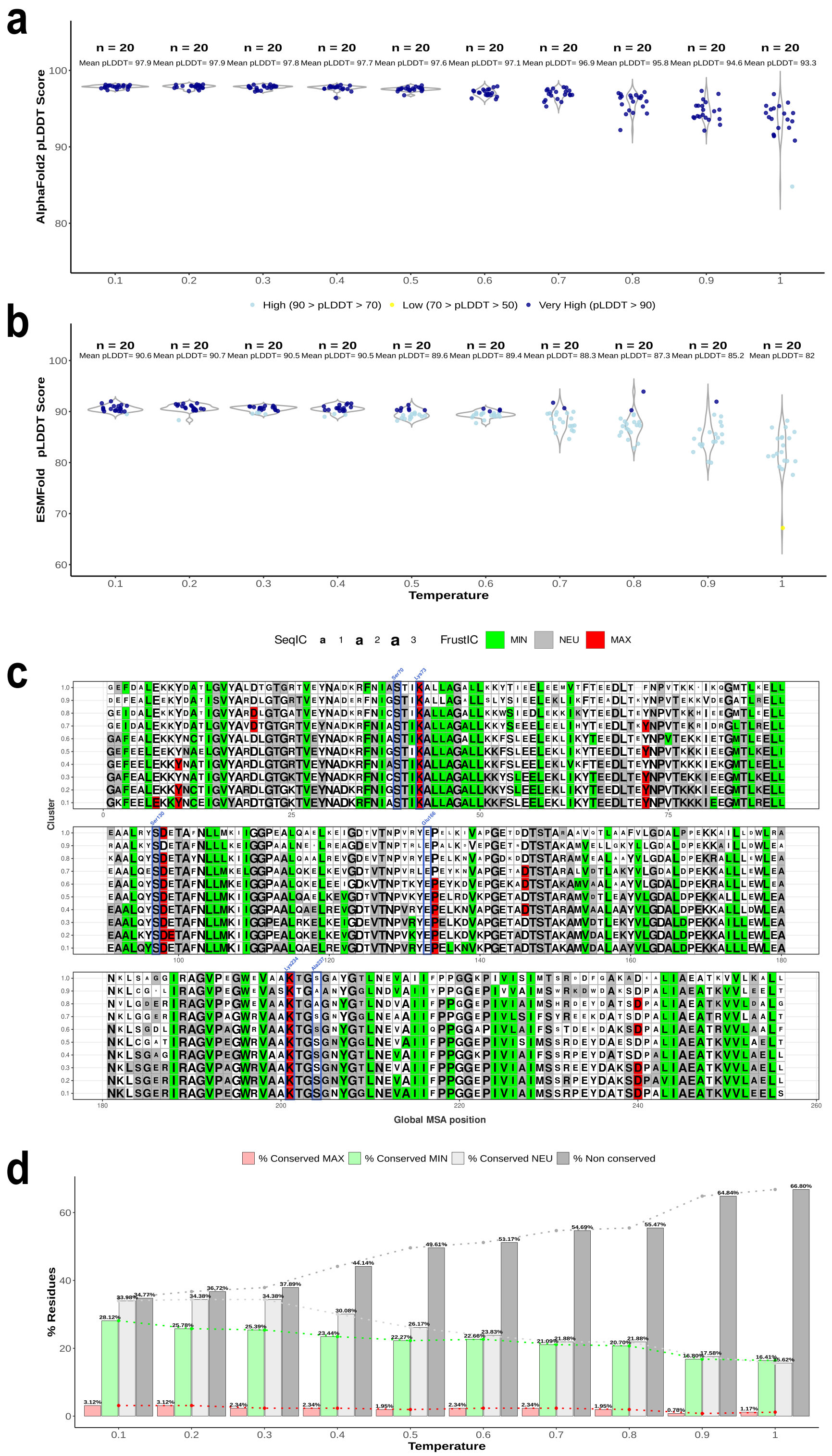


**Fig. S3.** ProteinMNN 4blm-A Single Target across temperatures. Violin plot for the pLDDT scores distribution of ProteinMPNN designs on the reference $\beta$-lactamase (PDB ID: 4blm-A) from AlphaFold2 modeling (a) and ESMFold modeling (b). 20 sequences were designed per sampling temperature, and all structures were predicted using both methods. Each point represents the pLDDT value for an individual design. For AlphaFold2, which generates five predictions per sequence, the best prediction is preselected. pLDDT scores are categorized as very high (pLDDT$>$90), high 70$<$pLDDT$<$90), or low (50$<$pLDDT$<$70). No predictions had very low scores (pLDDT$<$50) for any of the prediction methods. The means per temperature group are indicated above each violin box. (c) Comparison of the full-length sequence with consensus multiple MSFA for all sampling temperatures tested. The font size reflects the Sequence Information Content (identity conservation), and only positions with a conserved frustration state (FrustIC$>$0.5) are colored accordingly, while non-conserved positions are shown in white. The 6 catalytic residues are labeled in blue in all cases. Each cluster row summarizes the FrustraEvo results for a given MSFA, containing SRFI frustration data for an individual Single Target at a given temperature. The reference (4blm-A) is included in all cases to facilitate comparison of patterns between runs. (d) Bar chart summarizing the percentages of conserved and non-conserved residues in different temperature runs. Each set of bars represents the distribution of conservation states of residues at a given temperature. The percentages are labeled above each segment, and a dashed line highlights the trend across all cases. All results were obtained using SeqDist=12 by default.

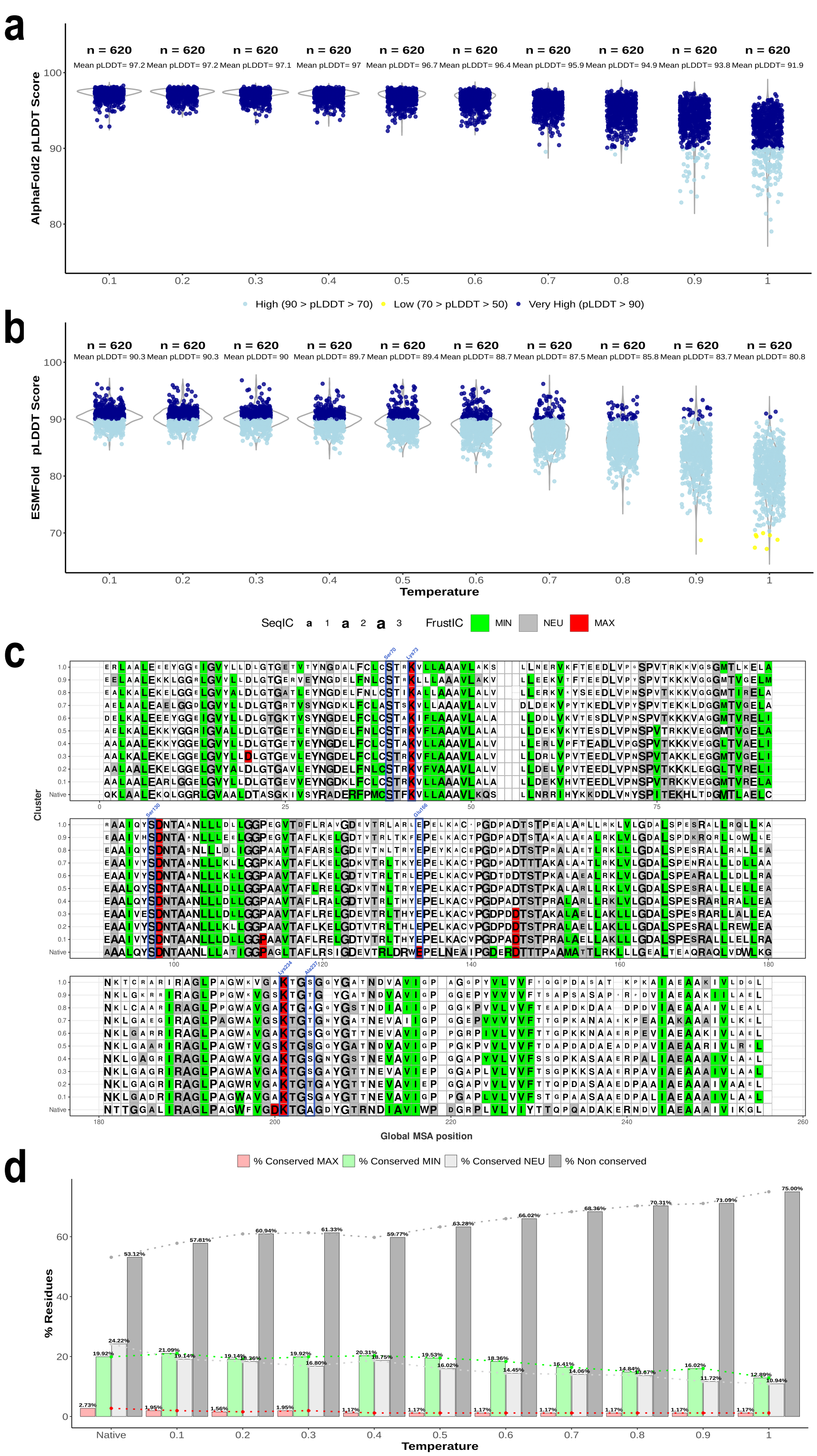


**Fig. S4.** ProteinMNN $\beta$-lactamases Family Target across temperatures. Violin plot for the pLDDT scores distribution of ProteinMPNN designs on the native $\beta$-lactamases family (n=31) from AlphaFold2 modeling (a) and ESMFold modeling (b). 20 sequences were designed per family member (individual targets) and sampling temperature, and all structures were predicted using both methods. Each point represents the pLDDT value for an individual design. For AlphaFold2, which generates five predictions per sequence, the best prediction is preselected. pLDDT scores are categorized as very high (pLDDT>90), high (70<pLDDT<90), or low (50<pLDDT<70). No predictions had very low scores (pLDDT<50) for any of the prediction methods. The means per temperature group are indicated above each violin box. (c) Comparison of the full-length sequence with consensus multiple MSFA for all sampling temperatures tested. The font size reflects the SeqIC (identity conservation), and only positions with a conserved frustration state (FrustIC>0.5) are colored accordingly, while non-conserved positions are shown in white. The 6 catalytic residues are labeled in blue in all cases. Each cluster row summarizes the FrustraEvo results for a given MSFA, containing SRFI frustration data for an individual Family Target at a given temperature. The reference (4blm-A) is included in all cases to facilitate comparison of patterns between runs. (d) Bar chart summarizing the percentages of conserved and non-conserved residues in different temperature runs. Each set of bars represents the distribution of conservation states of residues at a given temperature. The percentages are labeled above each segment, and a dashed line highlights the trend across all cases. All results were obtained using SeqDist=12 by default.

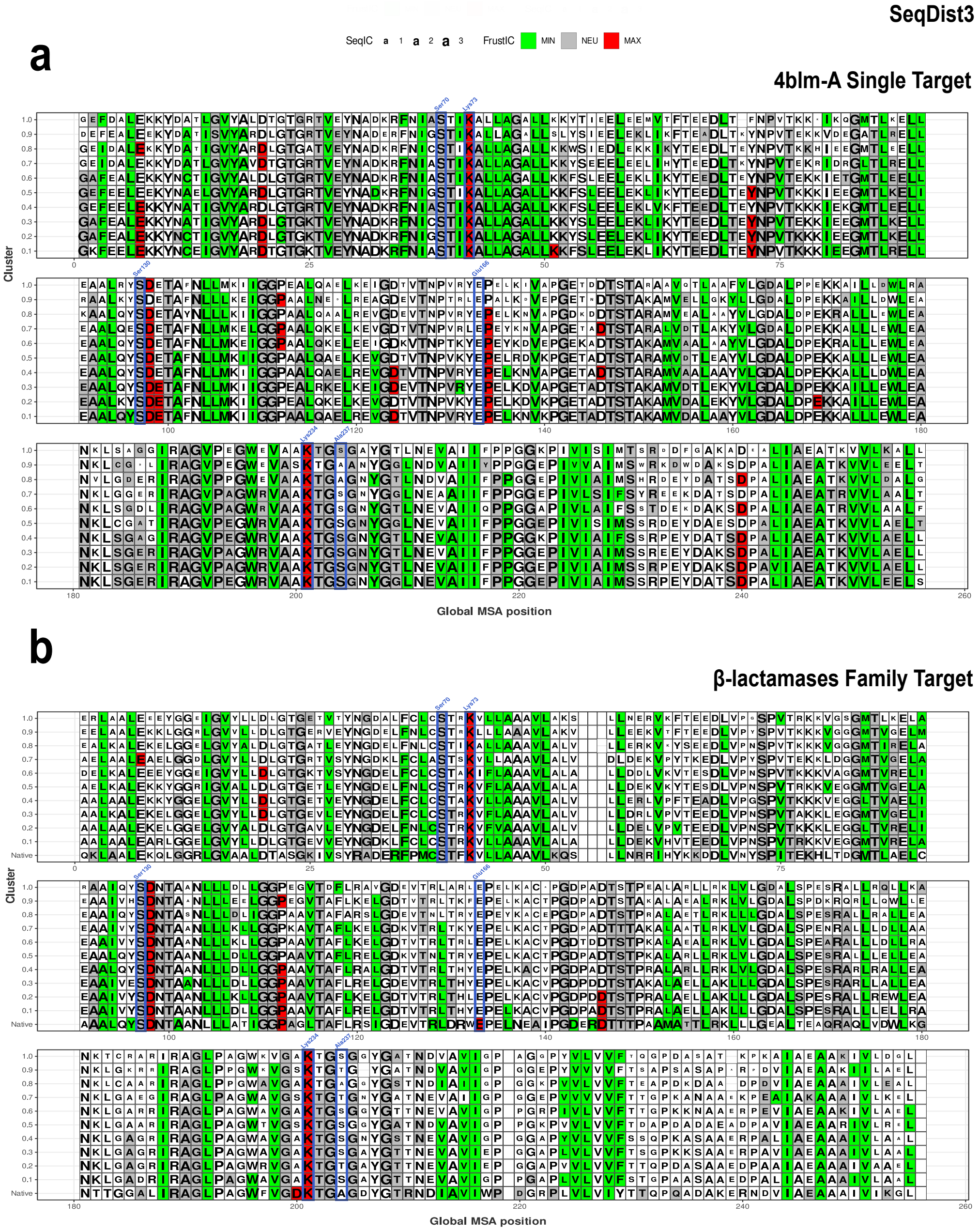


**Fig. S5.** SeqDist3 Frustratometer parameterization across ProteinMPNN $\beta$-lactamases. Comparison of the full-length sequence with consensus multiple MSFA for all sampling temperatures tested. The font size reflects the SeqIC (identity conservation), and only positions with a conserved frustration state (FrustIC$>$0.5) are colored accordingly, while non-conserved positions are shown in white. The 6 catalytic residues are labeled in blue in all cases. Each cluster row summarizes the FrustraEvo results for a given MSFA, containing SRFI frustration data for an individual Single Target (a) or Family Target (b) at a given temperature. The reference (4blm-A) is included in all cases to facilitate comparison of patterns between runs. All results were obtained using SeqDist=3, in comparison to equivalent runs (same input data across runs) in Figure S3C and Figure S4C that were obtained using SeqDist=12.

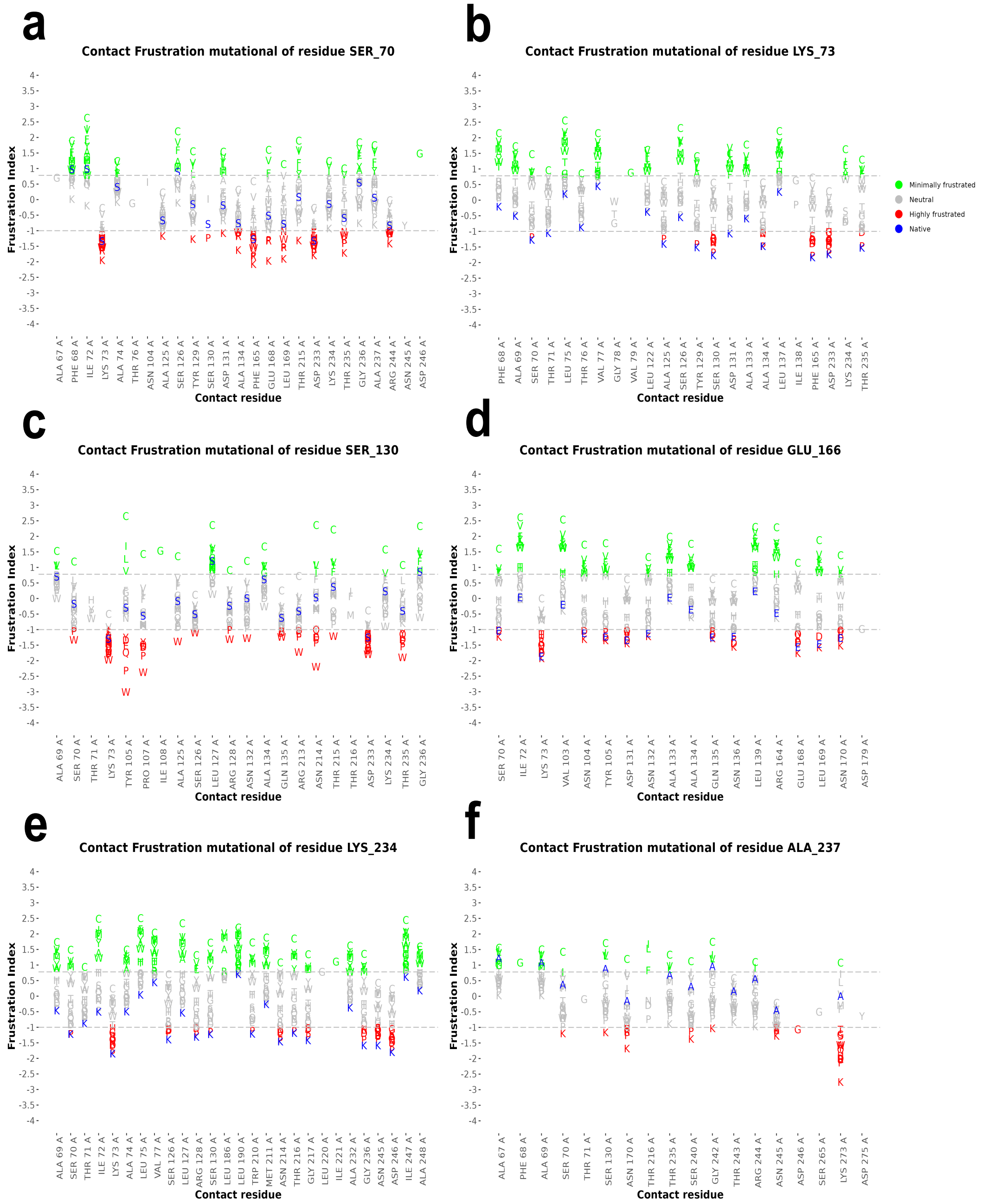


**Fig. S6.** Contact frustration changes upon mutations in 4blm-A. The mutational frustration index (MutFI) for the 6 catalytic residues annotated in native $\beta$-lactamases (a-f) for all canonical and alternative amino acids were considered. X axis: all possible contacts that form the native protein and the mutant. Y axis: frustration values, the canonical amino acids alternatives are represented in letters and coloured based on their frustration value. Native variants appear in blue. All results were obtained using SeqDist=12 by default.

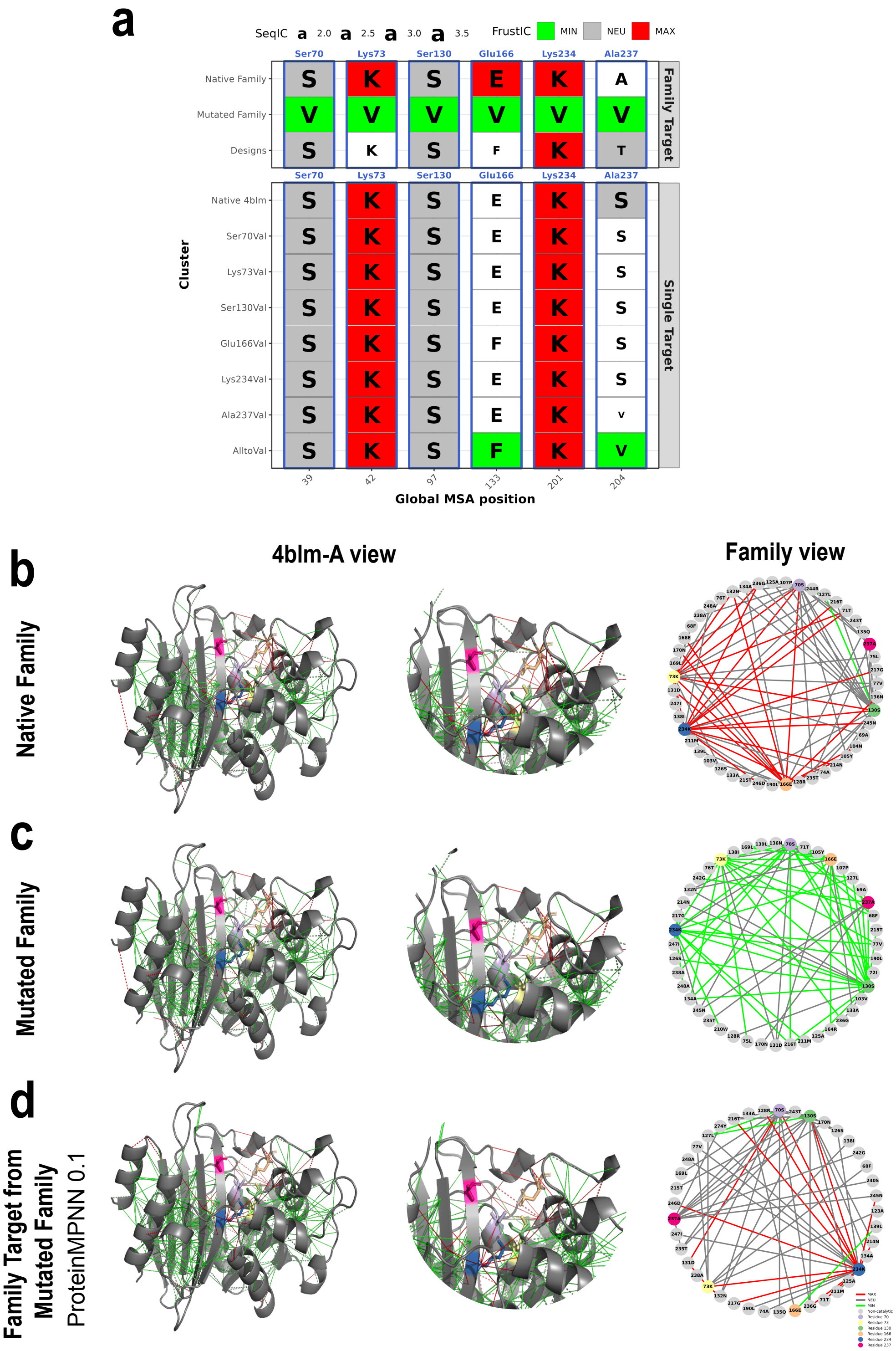


**Fig. S7.** Frustration conservation from in-silico $\beta$-lactamases mutants. (a) Consensus multiple MSFA full-sequence comparison, where the letter size reflects the Sequence Information Content (identity conservation) and only positions with a conserved frustration state (FrustIC$>$0.5) are colored accordingly, while non-conserved positions are shown in white. It contains 1) SRFI results for the Single Target approach from single (S70V, K73V, S130V, E166V, K234V) and multi-point (AlltoVal, all native identities from the catalytic residues mutated to valines) 4blm-A mutants, compared to the native pattern of 4blm-A and 2) SRFI results for the Family Target approach from multi-point mutants for the entire family (all the catalytic residues mutated to valines for all the family members and used as target (AlphaFold2 models) to ProteinMPNN, n=31), compared to the native family pattern and the one obtained when mutating the entire family. Mutational FI results from the (b) Native family, the (c) Mutated Family and the (d) Family Target approach from multi-point mutants for the entire family. Structures including zooming illustrate all the minimally and highly frustrated contacts mapped on the reference but according to family conservation, while the adjacent network displays the conserved contacts just involving the catalytic residues for the entire family, including all three frustration states. All results were obtained using SeqDist=12 by default.

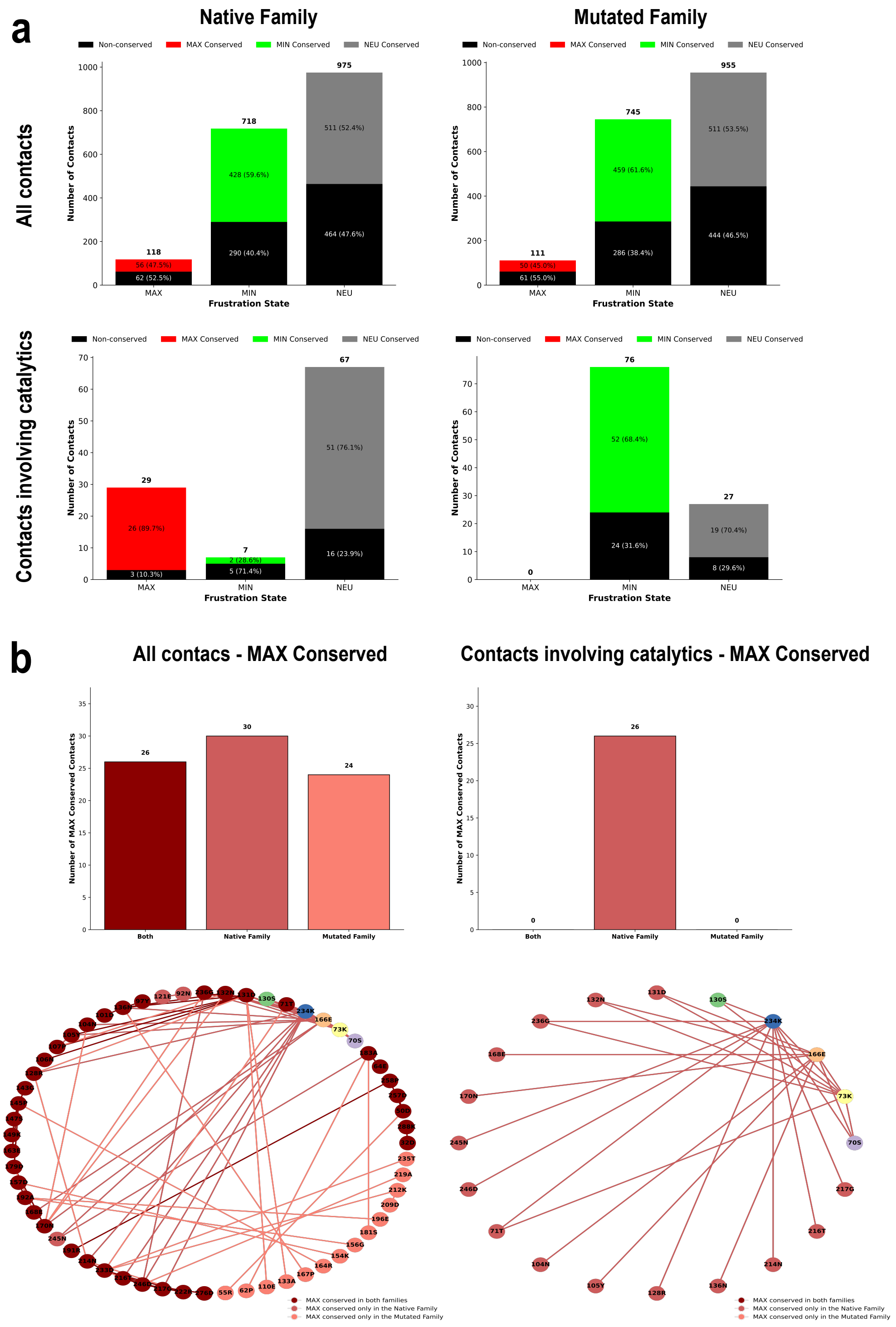


**Fig. S8.** (a) Distribution of conserved and non-conserved mutational contacts (MutFI) across frustration states in the native and the mutated $\beta$-lactamases families. Stacked bars represent the proportion of contacts that are non-conserved (black) or conserved (FrustIC$>$0.5) for the 3 frustration states: MAX conserved (red), MIN conserved (green), or NEU conserved (gray). Separate charts are shown for all contacts and only those involving catalytic residues. (b) Zoom in MAX conserved contacts shown for all contacts and only those involving catalytic residues, plotted accordingly in a contact network to visualize frustration redistribution. All results were obtained using SeqDist=12 by default.

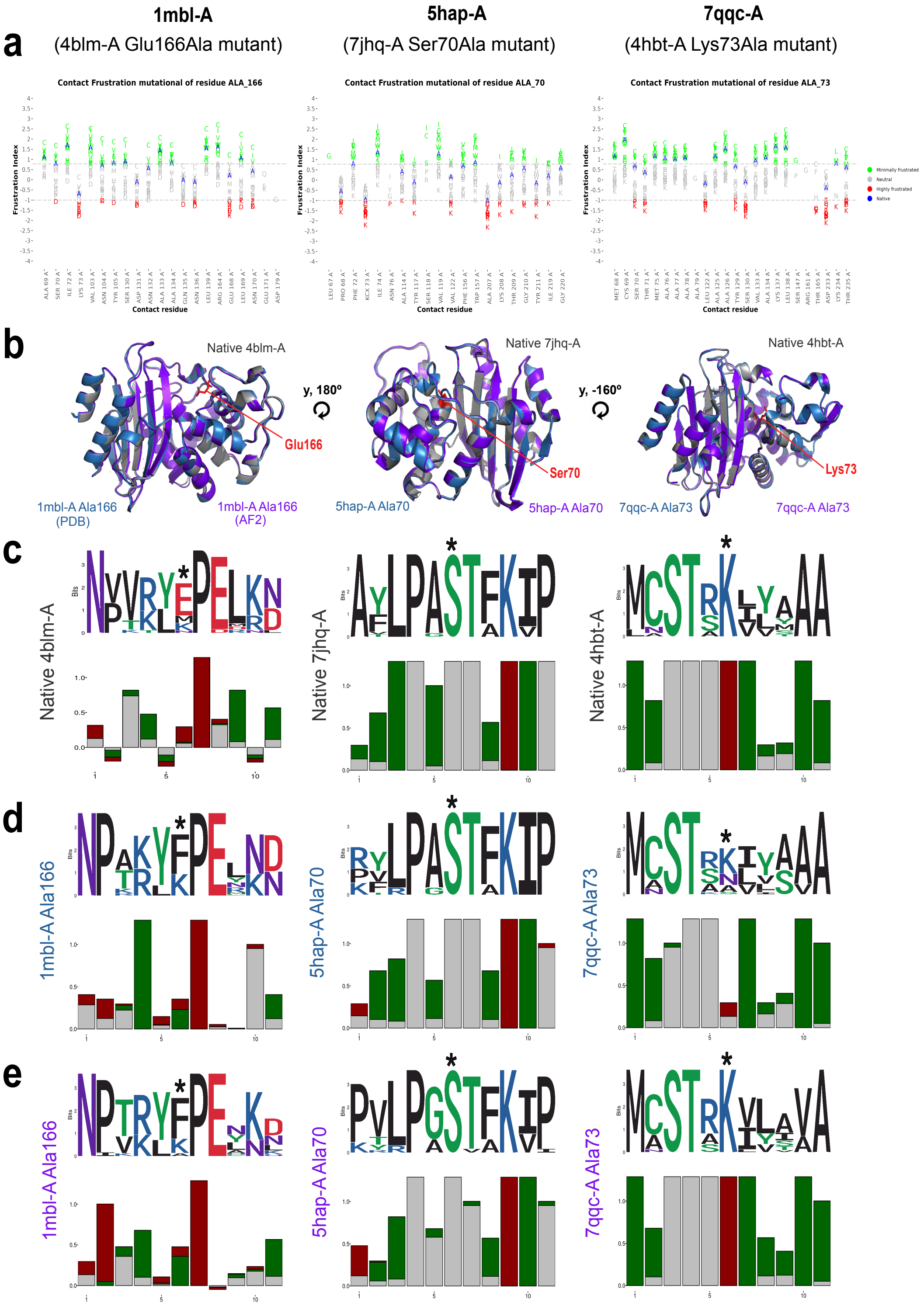


**Fig. S9.** 3 examples of experimental structures from the PDB containing mutations in a catalytic residue from $\beta$-lactamases: 1) 1mbl-A (4blm-A E166A), 2) 5hap-A (7jhq-A S70A), 3) 7qqc-A (4hbt-A K73A). For each panel is included (a) Contact frustration changes upon mutation for the mutational frustration index of the given catalytic residue for all canonical amino acids alternatives. X axis: all possible contacts that form the native protein and the mutants are shown. Y axis: frustration values, the canonical amino acids alternatives are represented in letters and coloured based on their frustration value. Native variants appear in blue. (b) Structural alignment including the wild-type protein (gray), the mutated experimental model (blue) and the mutated in silico model (purple). SRFI results are displayed summarized in partial Sequence and Frustration logos for (c) Single Target approach from the wild-type proteins (d) Single Target approach from the mutated experimental models and (e) Single Target approach from the mutated in silico models. The height of the letters and bars refers to identity (SeqIC) and frustration (FrstIC) conservation values respectively, per MSA column. The higher the height, the more conserved. Any of these runs included the input target as reference, but the best designed ensemble per target. All results were obtained using SeqDist=12 by default.

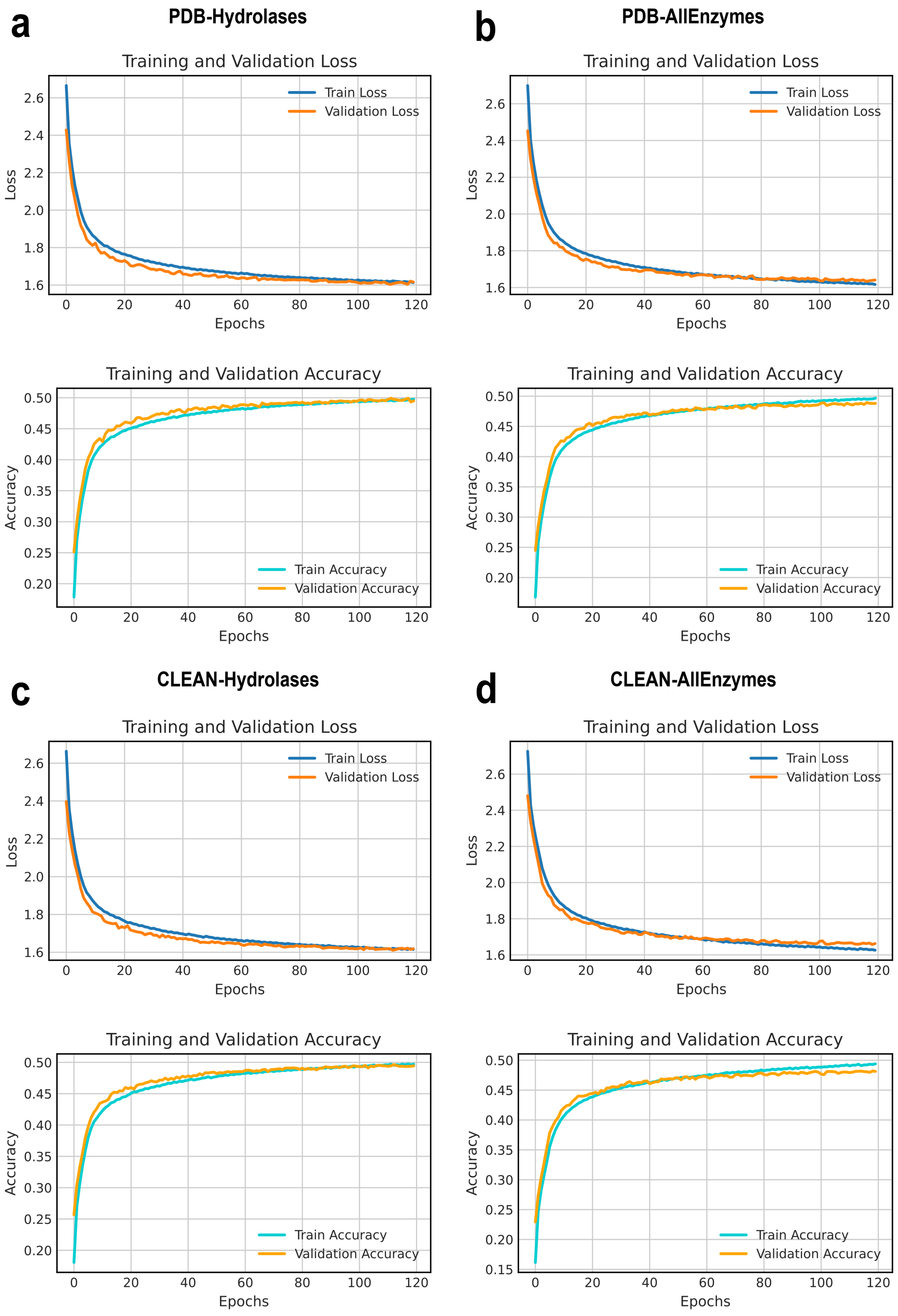


**Fig. S10.** Training and validation performance over 120 epochs for the retrained models. (a) PDB-Hydrolases (removing any protein annotated as an hydrolase in the PDB from the original ProteinMPNN training dataset). (b) PDB-AllEnzymes (removing any protein annotated as an enzyme in the PDB from the original ProteinMPNN training dataset). (c) CLEAN-Hydrolases (removing any protein predicted as an hydrolases by the CLEAN algorithm from the original ProteinMPNN training dataset). (d) CLEAN-AllEnzymes (removing any protein predicted as an enzyme by the CLEAN algorithm from the original ProteinMPNN training dataset). In each case: (Top) training and validation loss curves, showing progressive decrease and convergence as the model learns. (Bottom) Training and validation accuracy curves, indicating steady improvement and convergence toward $\simeq 50\%$ accuracy. The small gap between training and validation metrics suggests minimal overfitting and good generalization. CLEAN refers to the filtering based on the E.C numbers predicted by the CLEAN algorithm (22).

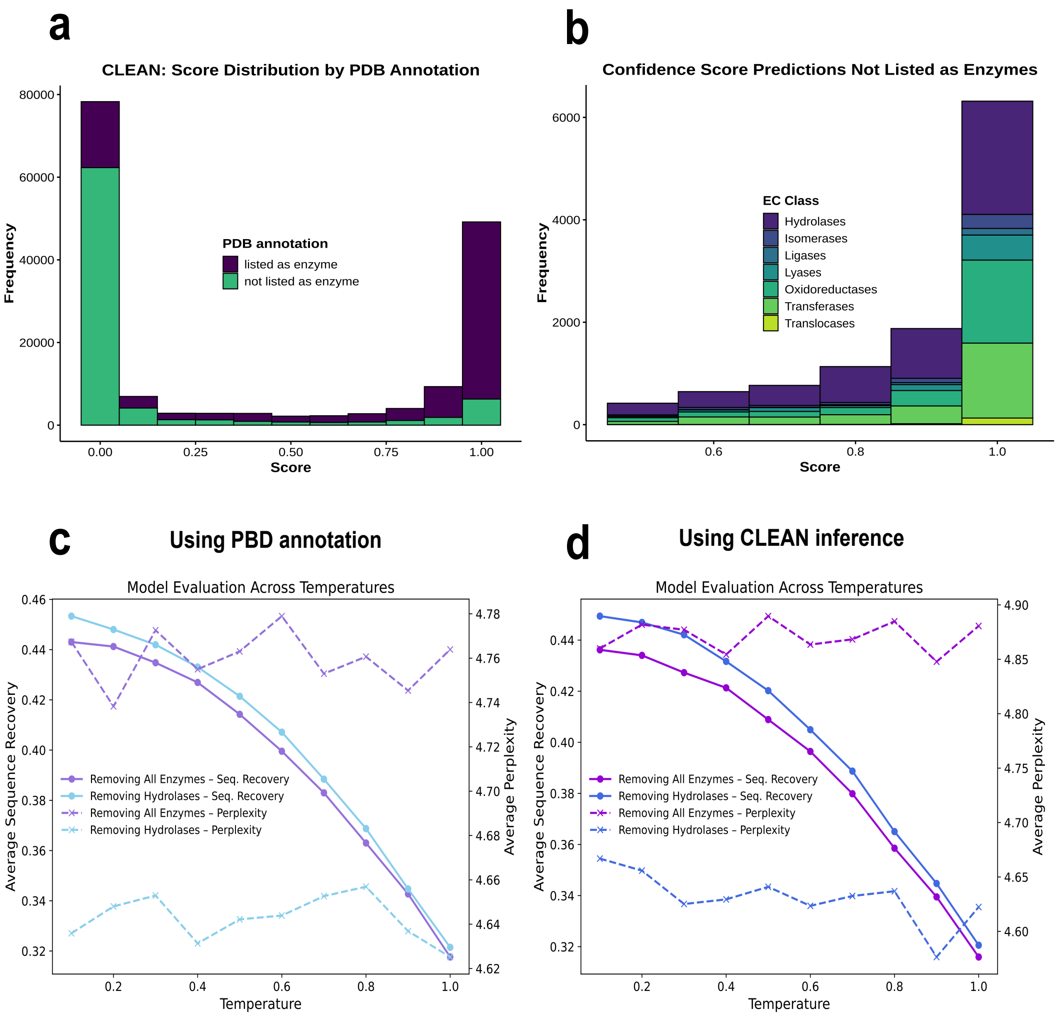


**Fig. S11.** (a) Confidence score distribution from the CLEAN algorithm (22) when inferring E.C numbers by max-separation (greedy approach that prioritizes EC numbers that have the maximum separation to other EC numbers in terms of the pairwise distance to the query sequence) for the full training dataset of ProteinMPNN, colored according to the PDB annotation. (b) Distribution subset for the entries with a confidence score higher than 0.5 and not listed as enzymes in the PDB database, colored according to the E.C inferred by CLEAN. Such classification allowed us to further filter the training dataset. (c) Re-trained models evaluation across temperatures when filtering training datasets according to the PDB annotation. (d) Re-trained models evaluation across temperatures when filtering training datasets according to the CLEAN algorithm predictions.

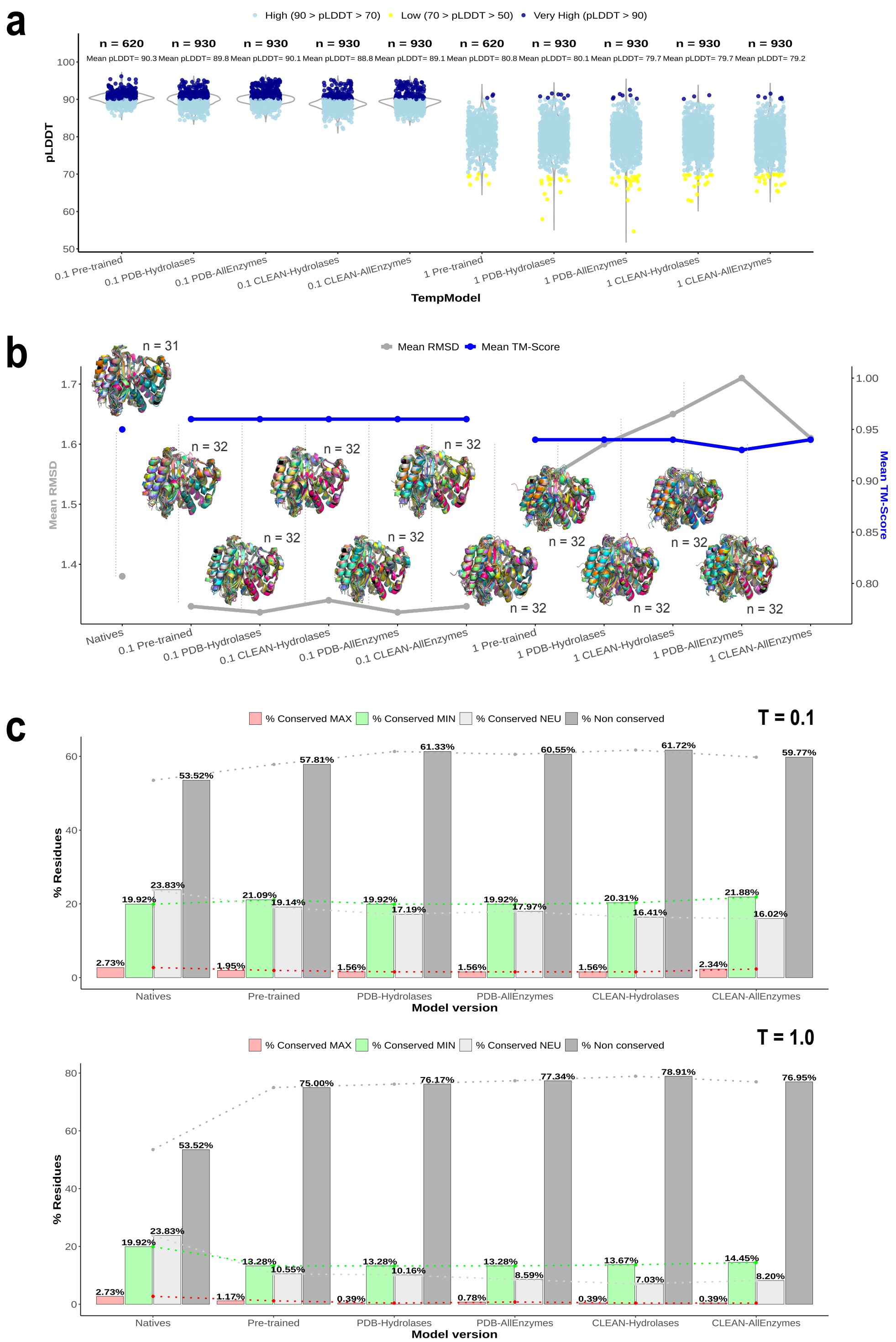


**Fig. S12.** (a) Violin plot for the ESMFold pLDDT scores distribution of ProteinMPNN designs on the native $\beta$-lactamases family (n=31), using the retrained models. 30 sequences were designed per family member (individual targets), sampling temperature (considering just the minimum and the maximum tested T) and model version. Each point represents the pLDDT value for an individual design, categorized as very high (pLDDT>90), high (70<pLDDT<90) or low (50<pLDDT<70). No predictions had very low scores (pLDDT<50). The means per temperature group are indicated above each violin box. Only the best design per target was selected to perform Family Target on top of them. (b) Structural evaluation of the selected datasets per TempModel version by the mean RMSD and the mean TMAlign score (23) when aligning designs versus the reference $\beta$-lactamase (4blm-A). (c) Bar chart summarizing the SRFI results from the different TempModel versions when performing Family Target. The native reference (4blm-A) is included in all cases to facilitate comparison between runs. Each set of bars represents the distribution of conservation states of residues at a given temperature per model version. The percentages are labeled above each segment, and a dashed line highlights the trend across all cases. All results were obtained using SeqDist=12 by default.

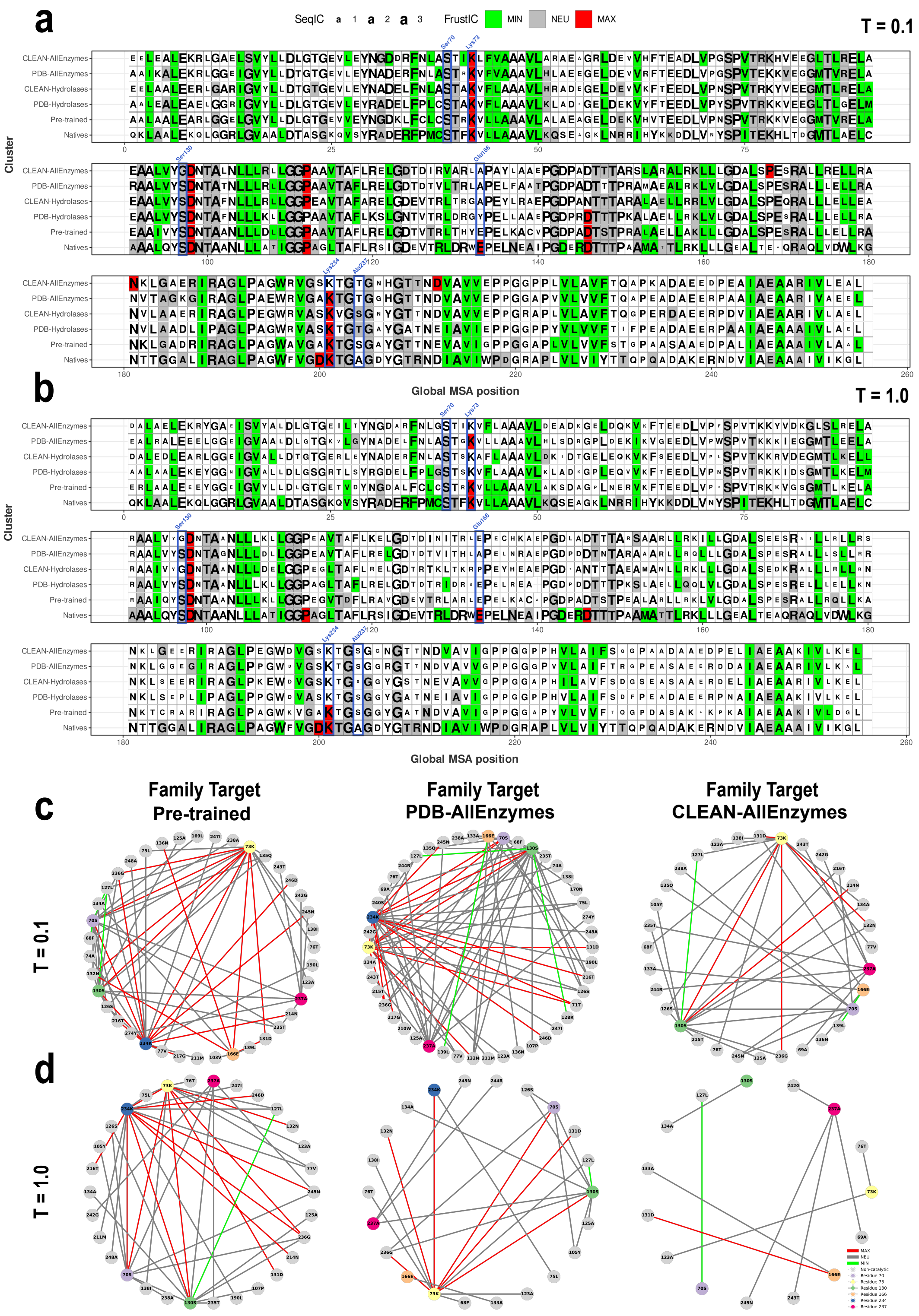


**Fig. S13.** Consensus MSFA full-sequence comparison across the pretrained and retrained ProteinMPNN versions, versus the native pattern of $\beta$-lactamases, for (a) the minimum (T=0.1) and (b) the maximum (T=1.0) sampling temperature. The letter size reflects the SeqIC (identity conservation) and only positions with a conserved frustration state (FrustIC$>$0.5) are colored accordingly, while non-conserved positions are shown in white. The 6 catalytic residues are labeled in blue in any case. Each cluster row summarises the FrustraEvo results for a given MSFA, containing SRFI frustration data from the different TempModel versions when performing Family Target using the native $\beta$-lactamases structures. The reference (4blm-A) is included in each case to facilitate pattern comparison among runs. MutFI frustration data for the network of conserved contacts involving the catalytic sites for the most restrictive retrained models versus the native network, for (c) the minimum and (d) the maximum sampling temperature as before. The 6 catalytic residues are highlighted (nodes) with distinct colors to differentiate them from the non-catalytics (gray). The contacts (edges) are colored according to the 3 frustration states. All results were obtained using SeqDist=12 by default.

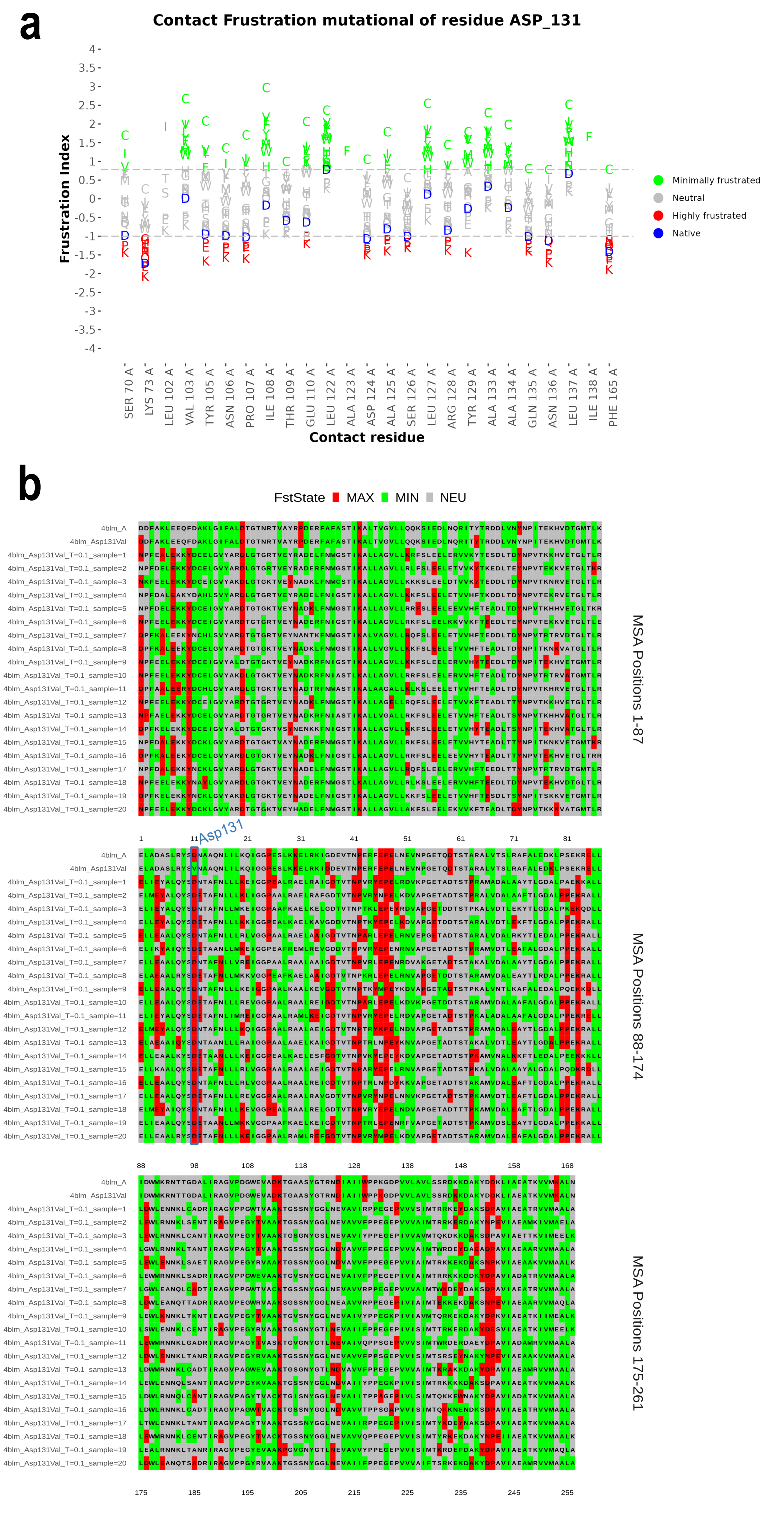


**Fig. S14.** (a) Changes in contact frustration upon mutation in the $\beta$-lactamase reference protein PDB ID: 4blm-A, for the mutational frustration index of D131 for all canonical amino acid alternatives. X-axis: all possible contacts formed by the native and mutant proteins are shown. Y-axis: frustration values; canonical amino acid alternatives are represented in letters and colored according to their frustration value. Native variants appear in blue. (b) Split Multiple Sequence Frustration Alignment (MSFA) corresponding to the SRFI results when performing Single Target analysis using the D131V mutant 4blm-A as the input target. 20 protein sequences designed by pre-trained ProteinMPNN and the native reference protein 4blm-A mutated and non-mutated are included (total n = 22). Positions are colored according to their frustration state, except for gaps. The D131 residue is marked to highlight its conservation in SeqIC and FrustIC. All results were obtained using SeqDist=12 by default.

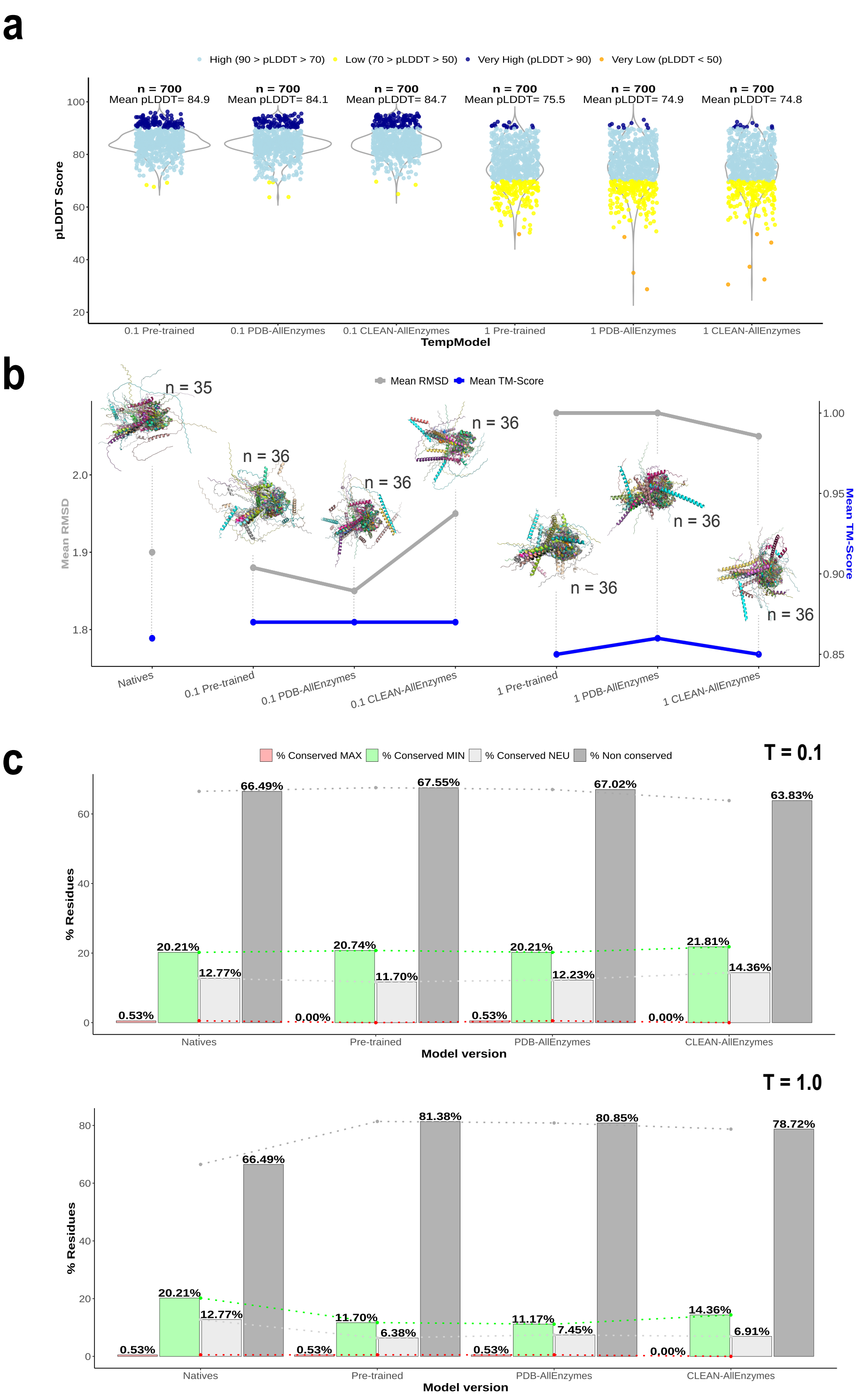


**Fig. S15.** (a) Violin plot for the ESMFold pLDDT scores distribution of ProteinMPNN designs on the native Ras family (n=35), using the more restrictive retrained models. 20 sequences were designed per family member (individual targets), sampling temperature (considering just the minimum and the maximum tested T) and model version. Each point represents the pLDDT value for an individual design, categorized as very high (pLDDT>90), high (70<pLDDT<90), low (50<pLDDT<70)or very low scores (pLDDT<50). The means per temperature group are indicated above each violin box. Only the best design per target was selected to perform Family Target on top of them. (b) Structural evaluation of the selected datasets per TempModel version by the mean RMSD and the mean TMAlign score when aligning designs versus the reference Ras (P01116-2, AlphaFold2 model). (c) Bar chart summarizing the SRFI results from the different TempModel versions when performing Family Target. The reference is included in all cases to facilitate comparison between runs. Each set of bars represents the distribution of conservation states of residues at a given temperature per model version. The percentages are labeled above each segment, and a dashed line highlights the trend across all cases. All results were performed using SeqDist=12 by default.

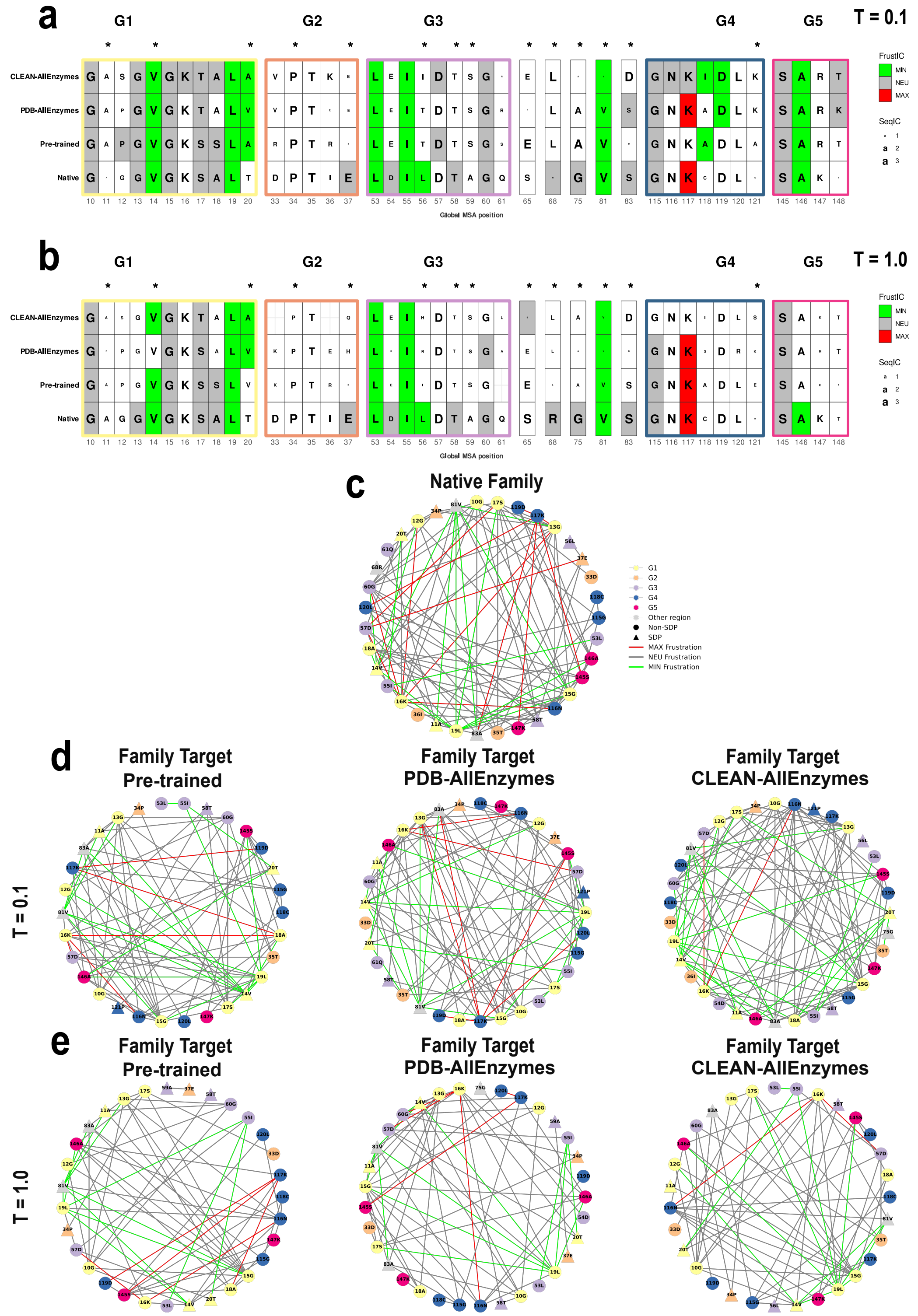


**Fig. S16.** Consensus multiple sequence frustration alignment (MSFA) comparing FrustIC and SeqIC results for the G1-G5 motifs and SDPs (marked with asterisks) (following annotation from (2)) across the pretrained and the more restrictive retrained ProteinMPNN versions, versus the native pattern, for (a) the minimum (T=0.1) and (b) the maximum (T=1.0) sampling temperature. Consensus amino acid identities are shown for each run. The letter size represents the SeqIC (identity conservation) and only positions with a conserved frustration state (FrustIC>0.5) are colored accordingly, while non-conserved positions are shown in white. Each cluster row summarises the FrustraEvo results for a given MSFA, containing SRFI frustration data from the different TempModel versions when performing Family Target using the native Ras structures. The reference (P01116-2) is included in each case to facilitate pattern comparison among runs. MutFI frustration data for the network of conserved contacts involving the residues within the G1-G5 domains and SDPs for the most restrictive retrained models versus the native network (c), for (d) the minimum and (e) the maximum sampling temperature as before. The residues (nodes) are shaped depending on whether they are SDPs or not, and are colored depending on the domain. The contacts (edges) are colored according to the 3 frustration states. All results were obtained using SeqDist=12 by default.

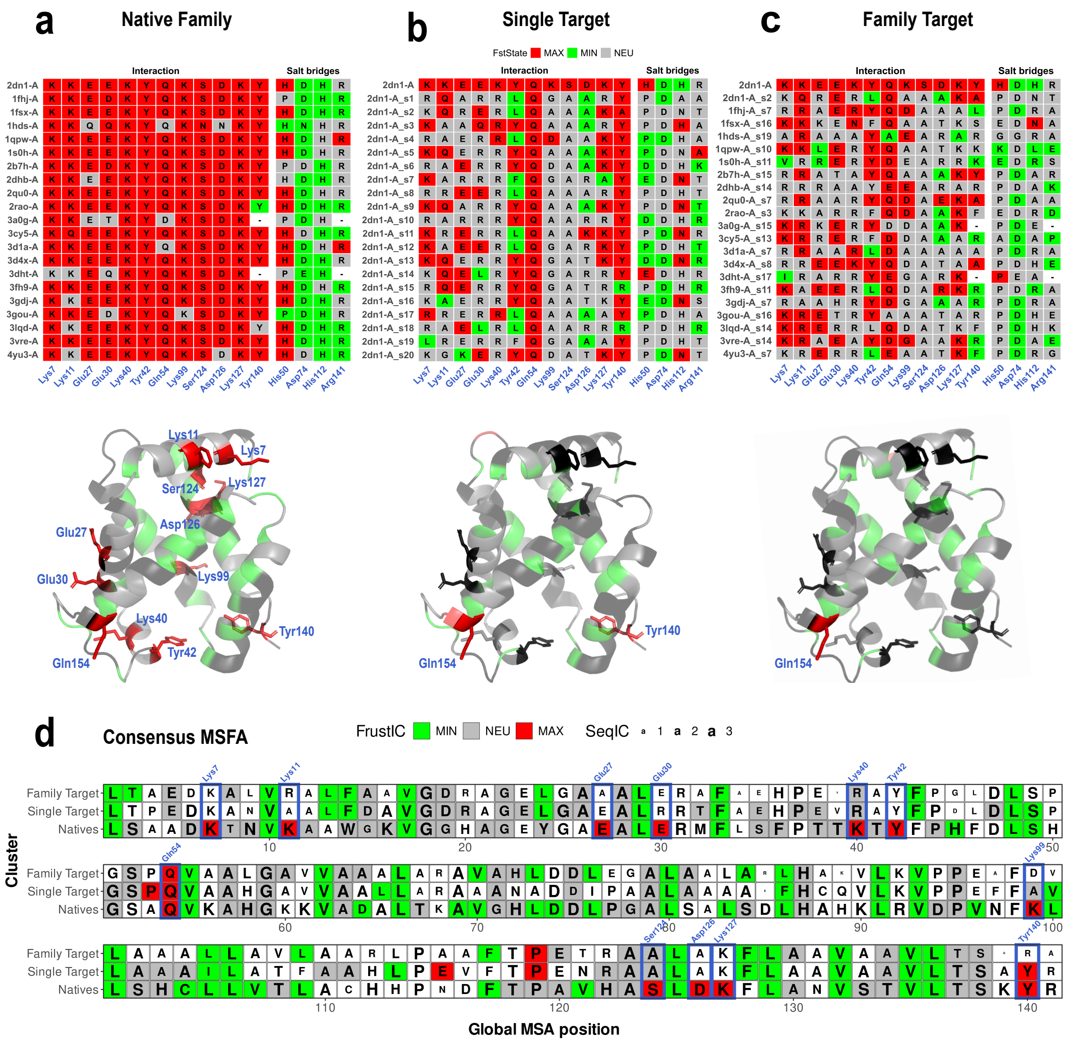


**Fig. S17.** Frustration patterns in native and ProteinMPNN $\alpha$-globins. (a) Positions subsets from the native MSFA including the 12 conserved and highly frustrated residues mostly involved in PPIs (K7, K11, E27, E30, K40, Y42, Q54, K99, S124, D126, K127 and Y140) and the four involved in salt bridges or chaperone interaction (K7/D74, E27/H112, E30/H50, K99/R141) annotated in native $\alpha$-globins (residues numbering corresponding to the reference 2dn1-A), including the SRFI results when using $\alpha$-globins native structures (n=21). The same subset is shown for both (b) the Human 2dn1-A Single Target approach (using 20 designs and the reference, n=21) and (c) the Family Target approach (using 1 design per native $\alpha$-globin and the reference, n=22) generated at the minimum sampling temperature (T=0.1). The designs chosen by native $\alpha$-globin (s number in y axis labels refers to sample number) are those that maximize the ESMFold pLDDT score for the target $\alpha$-globin in each case, among all those designed. Below, frustration is displayed for each MSFA position, mapped onto the human $alpha$-globin structure (reference), where regions in black are positions with a non-conserved frustration state (FrustIC$\leq$0.5) and the conserved ones (FrustIC$>$0.5) are coloured according to the frustration state that contributes the most to the overall FrustIC value. (d) Consensus multiple MSFA full-sequence comparison, where the letter size reflects the Sequence Information Content (identity conservation) and only positions with a conserved frustration state are colored accordingly, while non-conserved positions are shown in white. The 12 key positions mentioned earlier are labeled in blue in any case. Each cluster row summarises the FrustraEvo results for a given MSFA, containing SRFI frustration data for the previously described datasets. The reference 2dn1-A is included in each case to facilitate pattern comparison among runs.

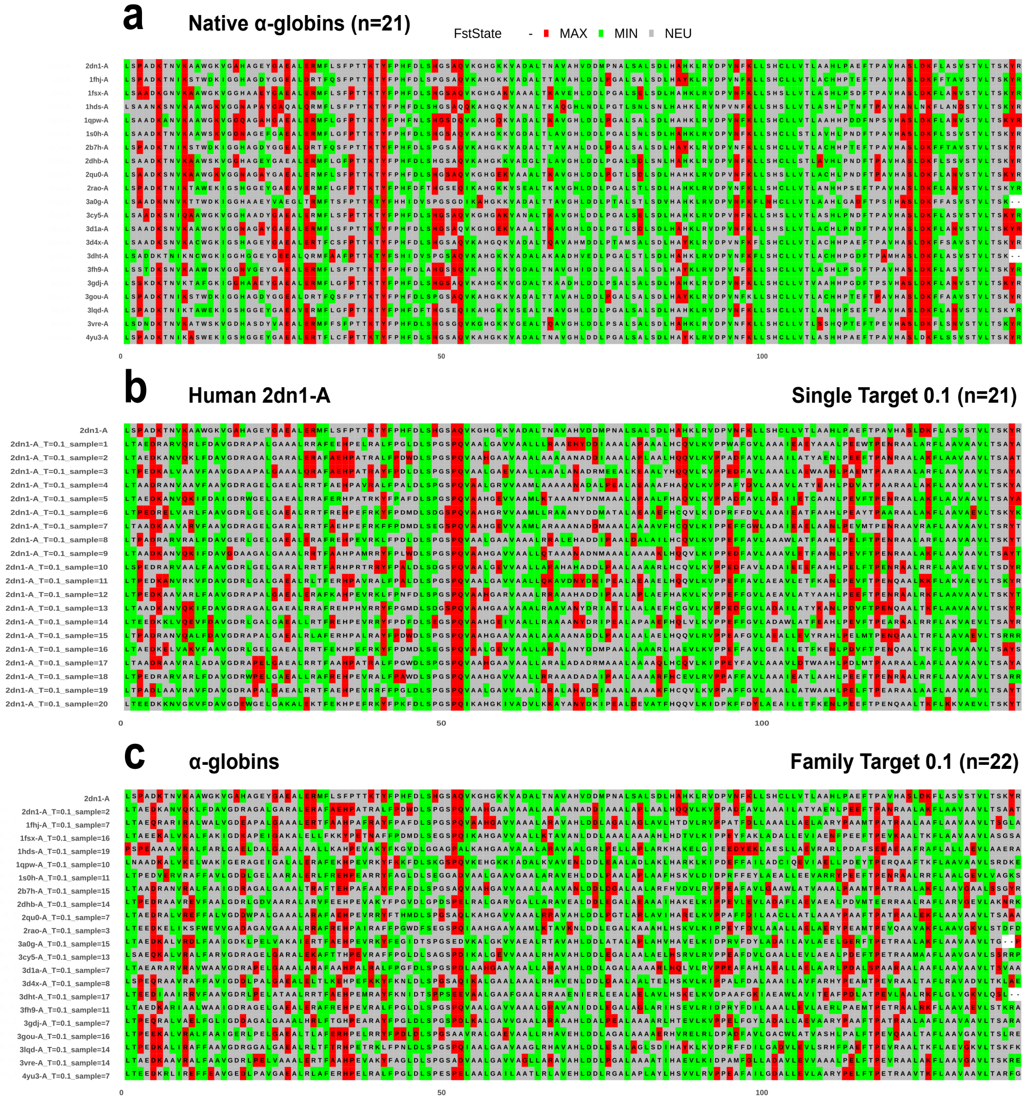


**Fig. S18.** Full Multiple Sequence Frustration Alignment (MSFA) corresponding to the SRFI results when (a) using $\alpha$-globins native structures (n=21), (b) the 2dn1-A Single Target approach (using 20 designed protein ensembles and the reference 2dn1-A, n=21) and c) the Family Target approach (using 1 ensemble per native $\alpha$-globin and the reference 2dn1-A, n=22) generated at the minimum sampling temperature (T=0.1). The minimum temperature refers to the minimum sampling temperature of the algorithm tested in our analyses, coinciding with the default. The ensembles chosen by native $\alpha$-globin in (c) are those that maximize the ESMFold pLDDT score for the target $\alpha$-globin in each case, among all those designed. Positions are colored according to its frustration state, except for the gaps, when SeqDist frustration remains setted by default to SeqDist=12.

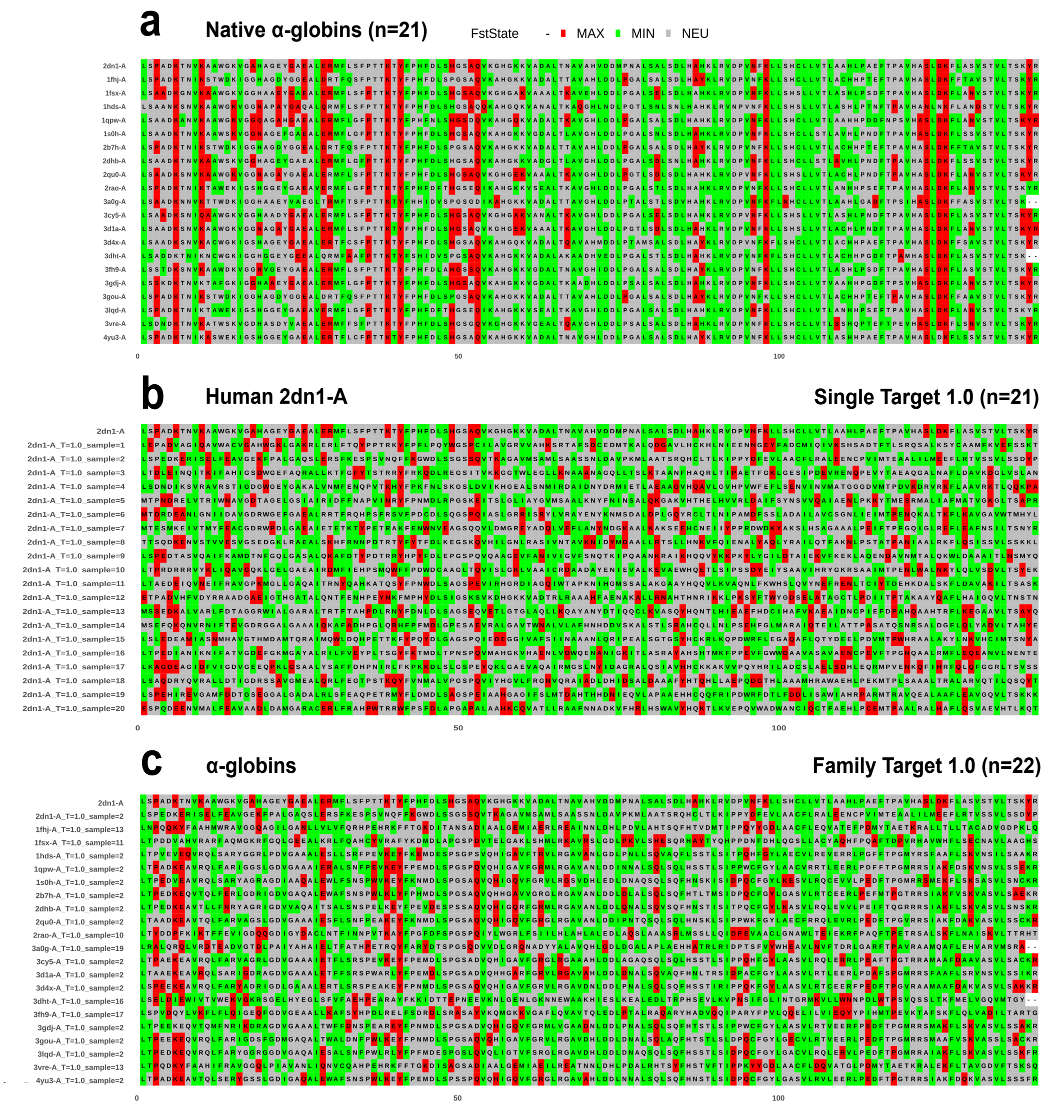


**Fig. S19.** Full Multiple Sequence Frustration Alignment (MSFA) corresponding to the SRFI results when (a) using $\alpha$-globins native structures (n=21), (b) the 2dn1-A Single Target approach (using 20 designed protein ensembles and the reference 2dn1-A, n=21) and C) the Family Target approach (using 1 ensemble per native $\alpha$-globin and the reference 2dn1-A, n=22) generated at the maximum sampling temperature (T=1.0). The maximum temperature refers to the maximum sampling temperature of the algorithm tested in our analyses. The ensembles chosen by native $\alpha$-globin in (c) are those that maximize the ESMFold pLDDT score for the target $\alpha$-globin in each case, among all those designed. Positions are colored according to its frustration state, except for the gaps, when SeqDist frustration remains settled by default to SeqDist=12.

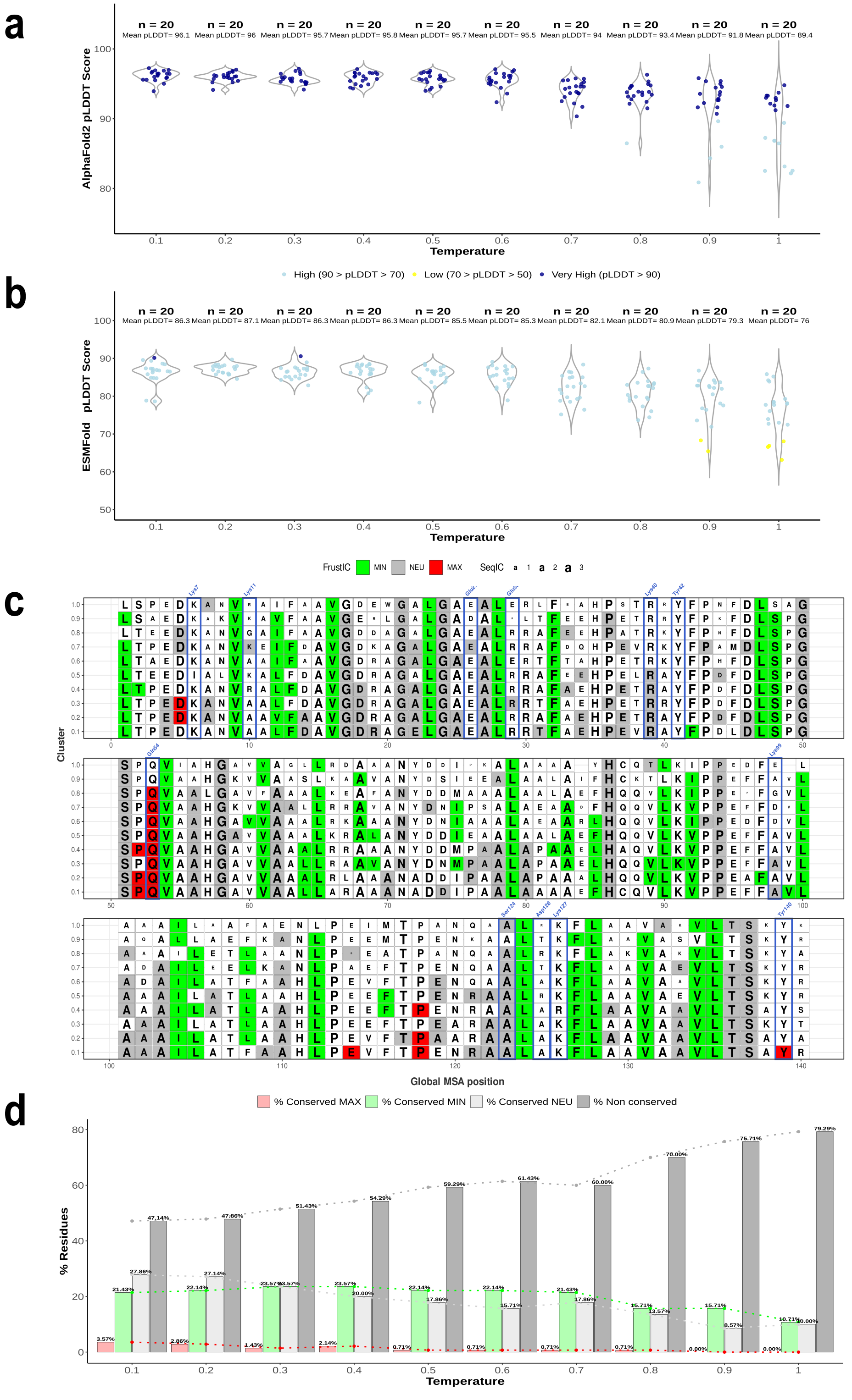


**Fig. S20.** Violin plot for the pLDDT scores distribution of ProteinMPNN designs on the reference $\alpha$-globin (PDB ID: 2dn1-A) from AlphaFold2 modeling (a) and ESMFold modeling (b). 20 sequences were designed per sampling temperature, and all structures were predicted using both methods. Each point represents the pLDDT value for an individual design. For AlphaFold2, which generates five predictions per sequence, the best prediction is preselected. pLDDT scores are categorized as very high (pLDDT$>$90), high (70$<$pLDDT$<$90), or low (50$<$pLDDT$<$70). No predictions had very low scores (pLDDT$<$50) for any of the prediction methods. The means per temperature group are indicated above each violin box. (c) Comparison of the full-length sequence with consensus multiple MSFA for all sampling temperatures tested. The font size reflects the SeqIC (sequence conservation), and only positions with a conserved frustration state (FrustIC$>$0.5) are colored accordingly, while non-conserved positions are shown in white. Residues involved in PPIs are labeled in blue in all cases. Each cluster row summarizes the FrustraEvo results for a given MSFA, containing SRFI frustration data for an individual Single Target at a given temperature. The reference (2dn1-A) is included in all cases to facilitate comparison of patterns between runs. (d) Bar chart summarizing the percentages of conserved and non-conserved residues in different temperature runs. Each set of bars represents the distribution of conservation states of residues at a given temperature. The percentages are labeled above each segment, and a dashed line highlights the trend across all cases. All results were obtained using SeqDist=12 by default.

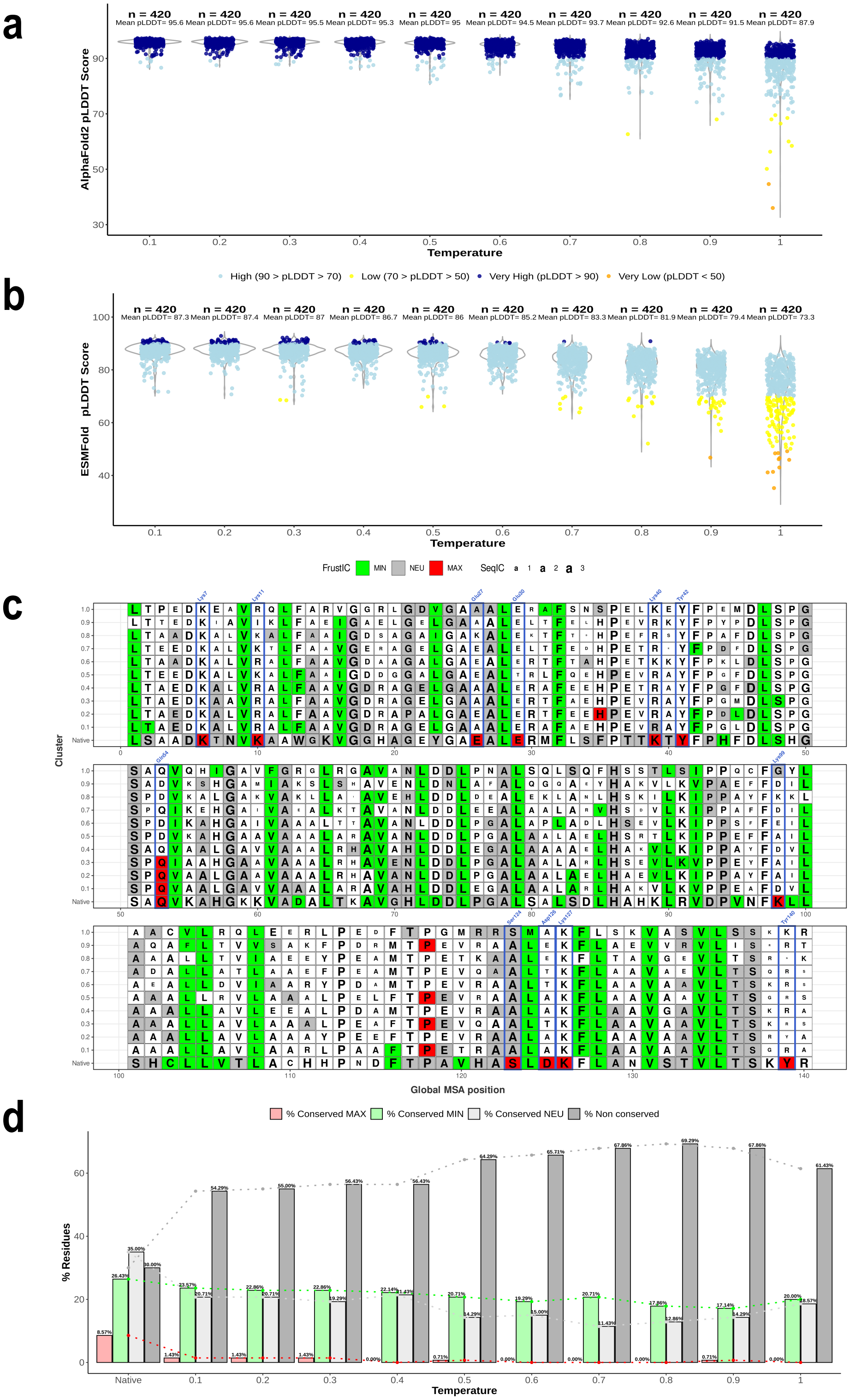


**Fig. S21.** Violin plot for the pLDDT scores distribution of ProteinMPNN designs on the native $\alpha$-globin family (n=21) from AlphaFold2 modeling (a) and ESMFold modeling (b). 20 sequences were designed per family member (individual targets) and sampling temperature, and all structures were predicted using both methods. Each point represents the pLDDT value for an individual design. For AlphaFold2, which generates five predictions per sequence, the best prediction is preselected. pLDDT scores are categorized as very high (pLDDT>90), high (70<pLDDT<90), low (50<pLDDT<70) or very low (pLDDT<50). The means per temperature group are indicated above each violin box. (d) Comparison of the full-length sequence with consensus multiple MSFA for all sampling temperatures tested. The font size reflects the SeqIC (sequence conservation), and only positions with a conserved frustration state (FrustIC>0.5) are colored accordingly, while non-conserved positions are shown in white. Residues involved in PPIs are labeled in blue in all cases. Each cluster row summarizes the FrustraEvo results for a given MSFA, containing SRFI frustration data for an individual Family Target at a given temperature. The native reference (2dn1-A) is included in all cases to facilitate comparison of patterns between runs. (d) Bar chart summarizing the percentages of conserved and non-conserved residues in different temperature runs. Each set of bars represents the distribution of conservation states of residues at a given temperature. The percentages are labeled above each segment, and a dashed line highlights the trend across all cases. All results were obtained using SeqDist=12 by default.

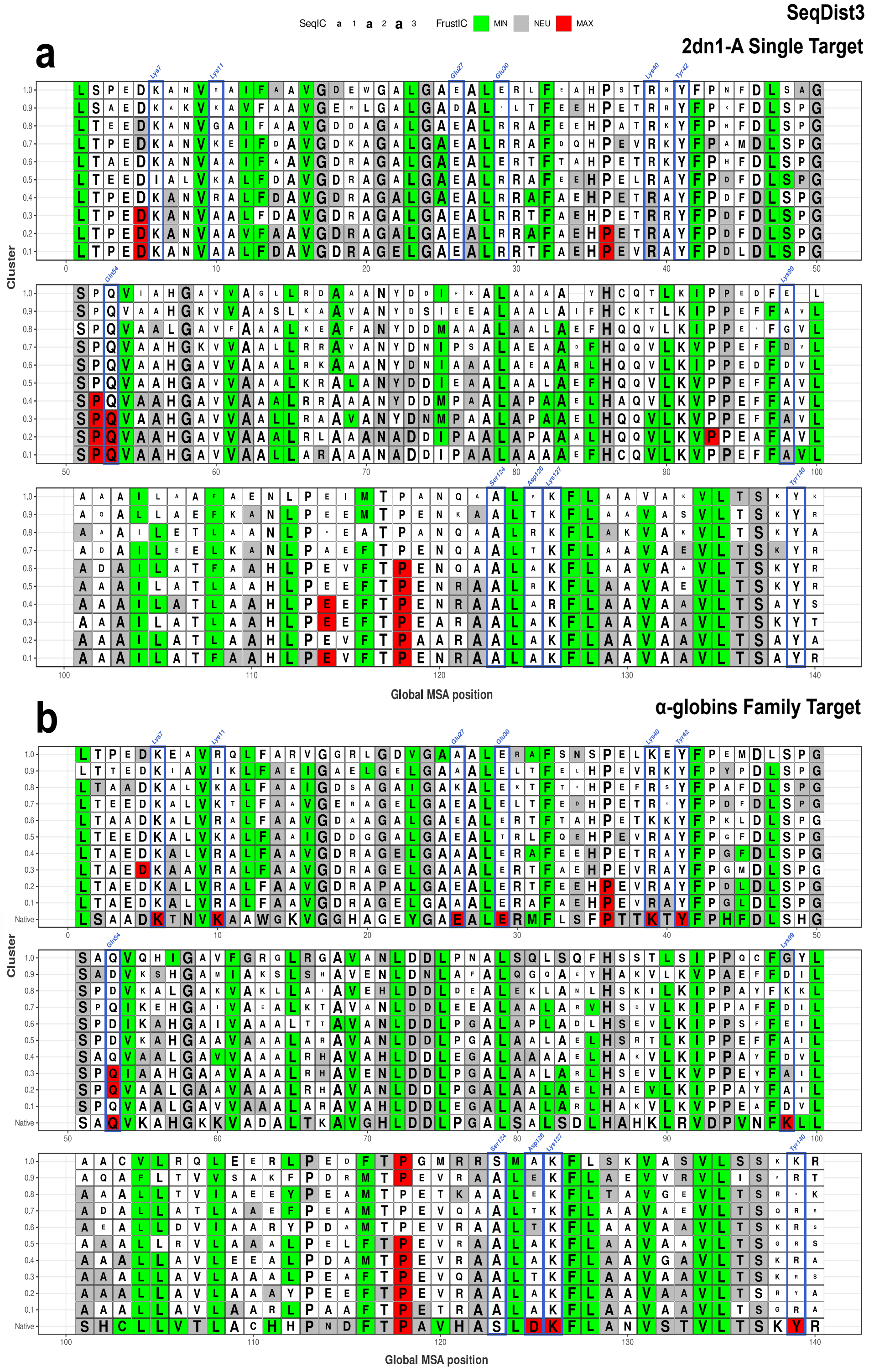


**Fig. S22.** Comparison of the full-length sequence with consensus multiple MSFA for all sampling temperatures tested. The font size reflects the SeqIC (sequence conservation), and only positions with a conserved frustration state (FrustIC>0.5) are colored accordingly, while non-conserved positions are shown in white. The interest residues are labeled in blue in all cases. Each cluster row summarizes the FrustraEvo results for a given MSFA, containing SRFI frustration data for an individual Single Target (the reference) (a) or Family Target (b) at a given temperature. The native reference (2dn1-A) is included in all cases to facilitate comparison of patterns between runs. All results were obtained using SeqDist=3, in comparison to equivalent runs (same input data across runs) in Figure S20C and Figure S21C that were obtained using SeqDist=12.

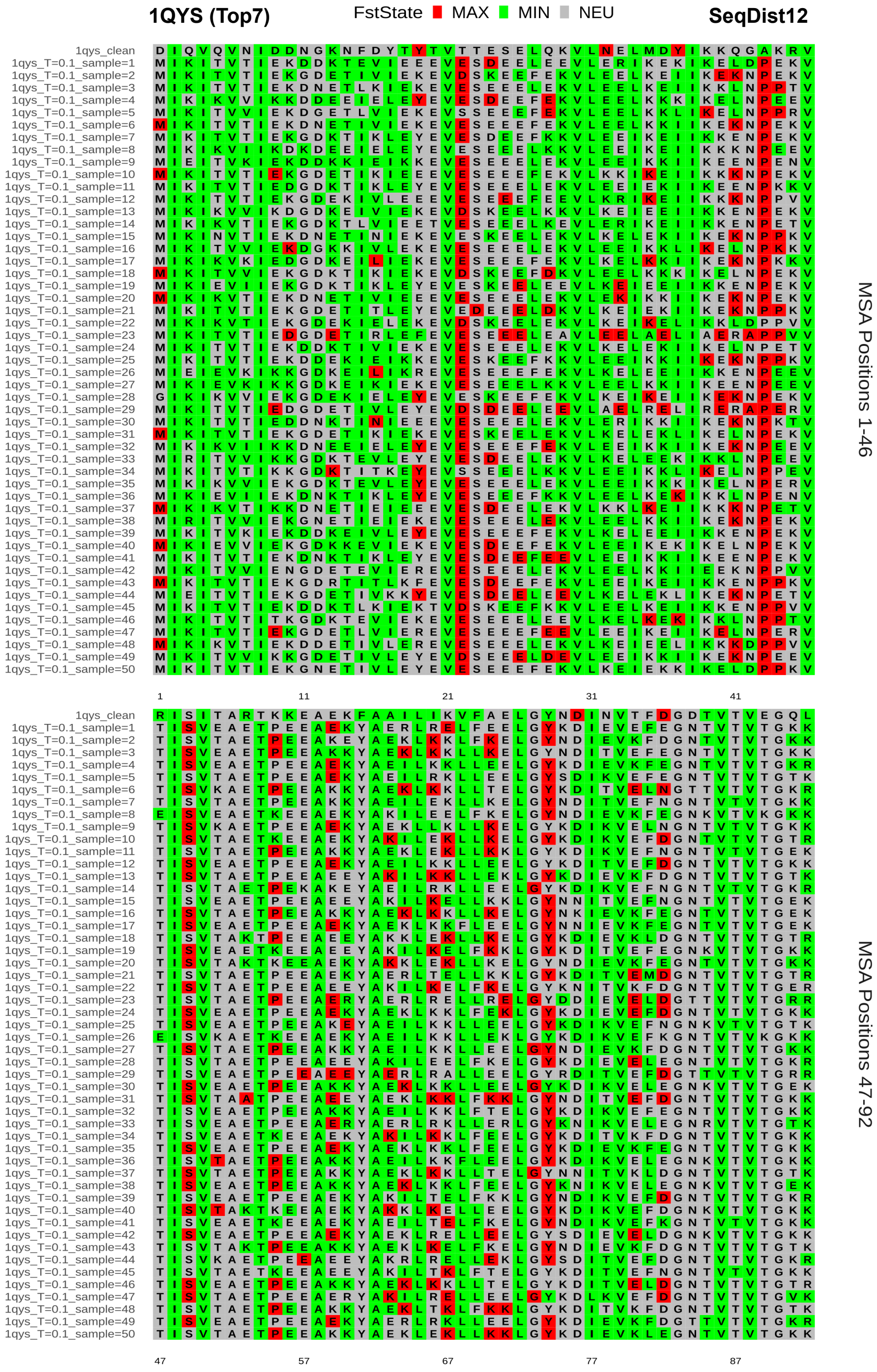


**Fig. S23.** Multiple Sequence Frustration Alignment (MSFA) mapping the SRFI results when performing Single Target analysis using the PDB ID: 1QYS de novo protein (Top7) as the input target. 50 sequences designed by the pre-trained ProteinMPNN and the reference protein are included (total n = 51). Positions are colored according to their frustration state, when SeqDist frustration parameters remain setted by default to SeqDist=12.

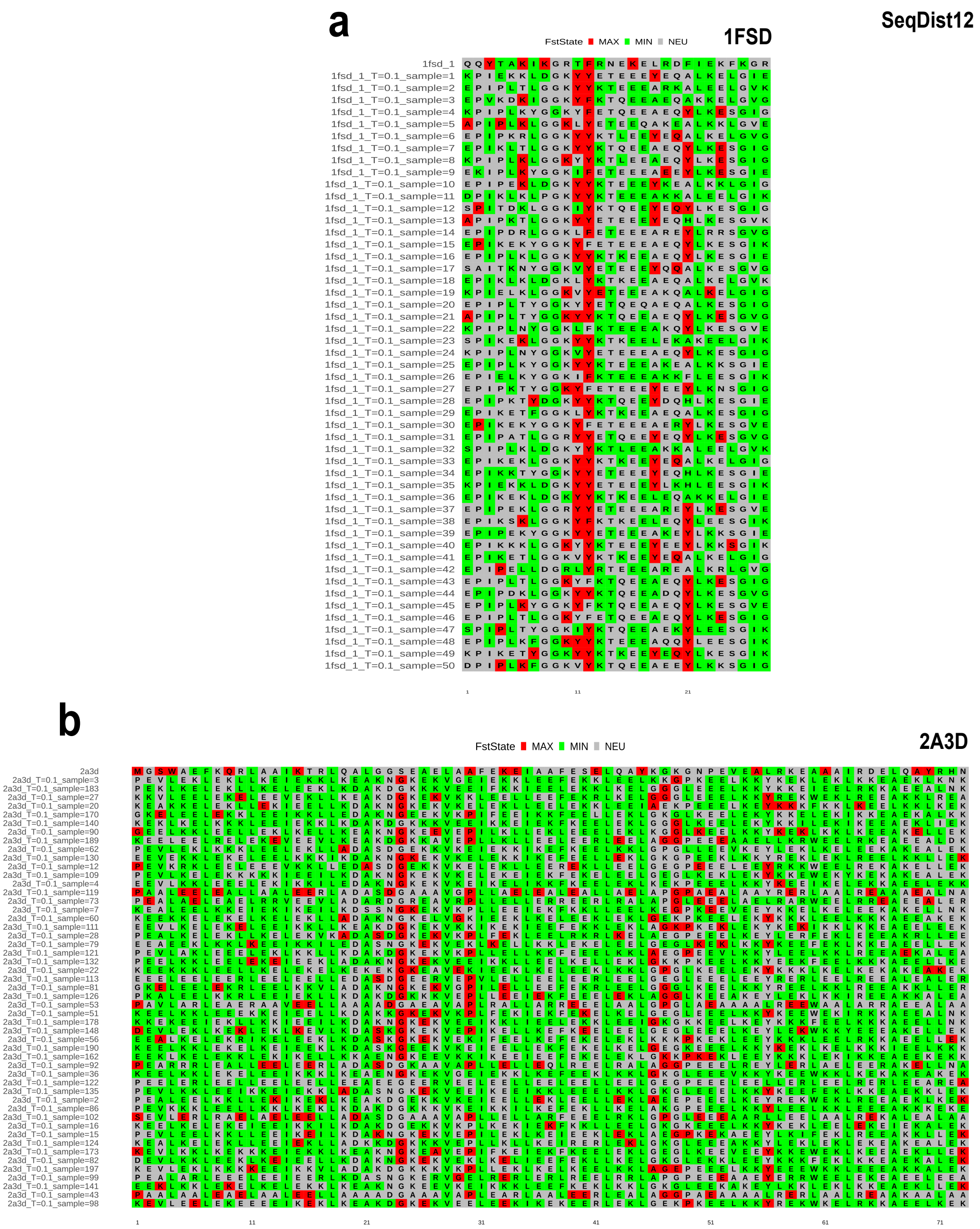


**Fig. S24.** Multiple Sequence Frustration Alignment (MSFA) corresponding to the SRFI results when performing Single Target analysis using (a) the 1FSD de novo protein and (b) the 2A3D de novo protein as input targets. 20 protein ensembles designed by the pre-trained ProteinMPNN and the reference protein are included (total n = 51). Positions are colored according to their frustration state, when SeqDist frustration parameters remain setted by default to SeqDist=12.

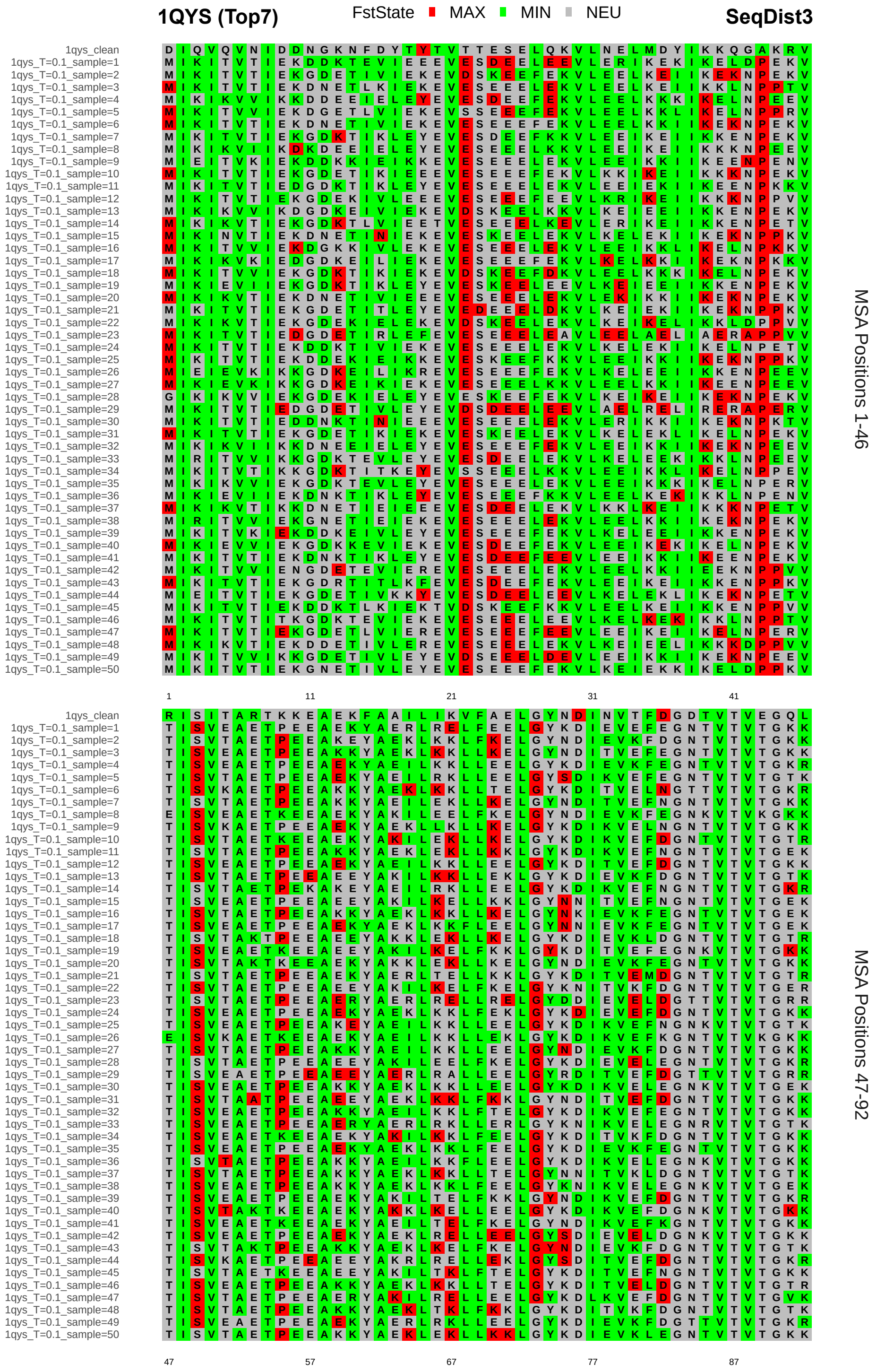


**Fig. S25.** Multiple Sequence Frustration Alignment (MSFA) mapping the SRFI results when performing Single Target analysis using the PDB ID: 1QYS de novo protein (Top7) as the input target. 50 sequences designed by the pre-trained ProteinMPNN and the reference protein are included (total n = 51). Positions are colored according to their frustration state, when SeqDist frustration parameter is setted to SeqDist=3.

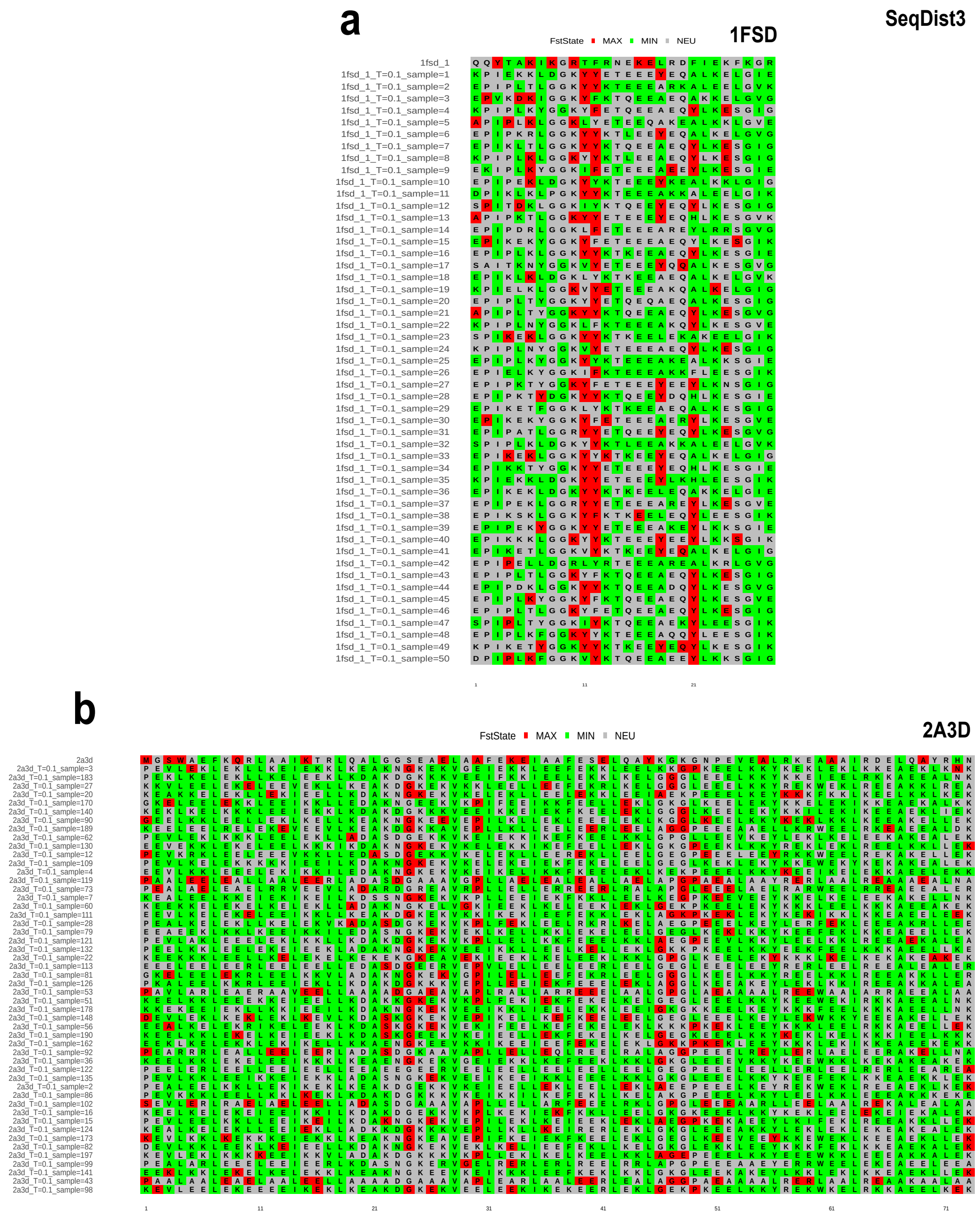


**Fig. S26.** Multiple Sequence Frustration Alignment (MSFA) corresponding to the SRFI results when performing Single Target analysis using (a) the 1FSD de novo protein and (b) the 2A3D de novo protein as input targets. 20 protein ensembles designed by the pre-trained ProteinMPNN and the reference protein are included (total n = 51). Positions are colored according to their frustration state, when SeqDist frustration parameters is setted to SeqDist=3.

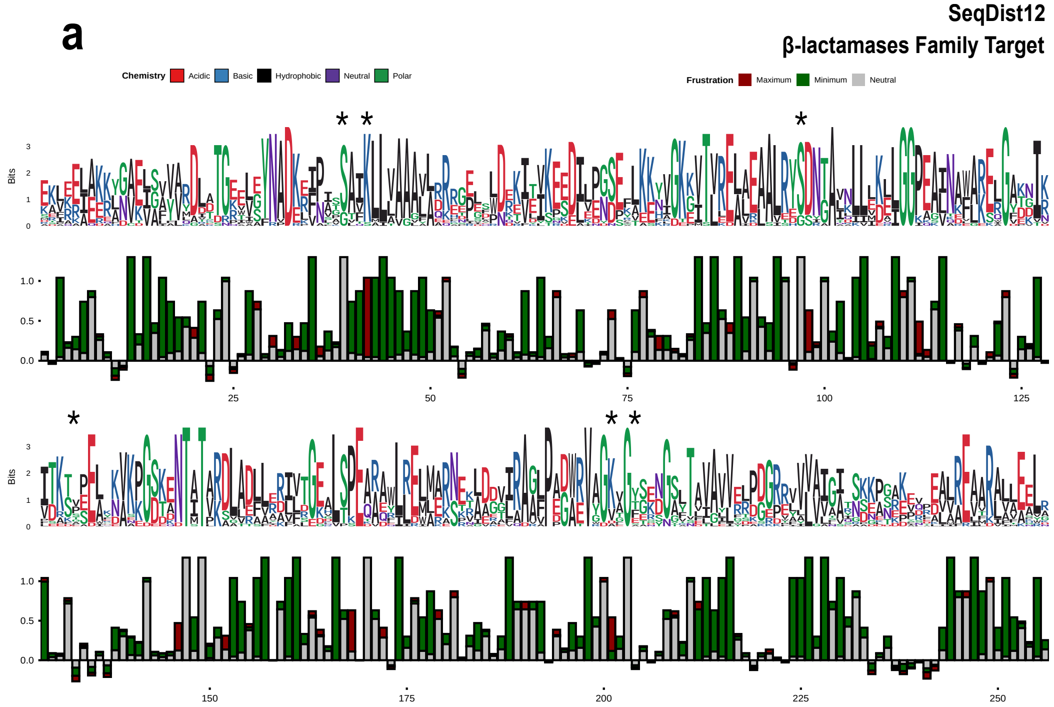


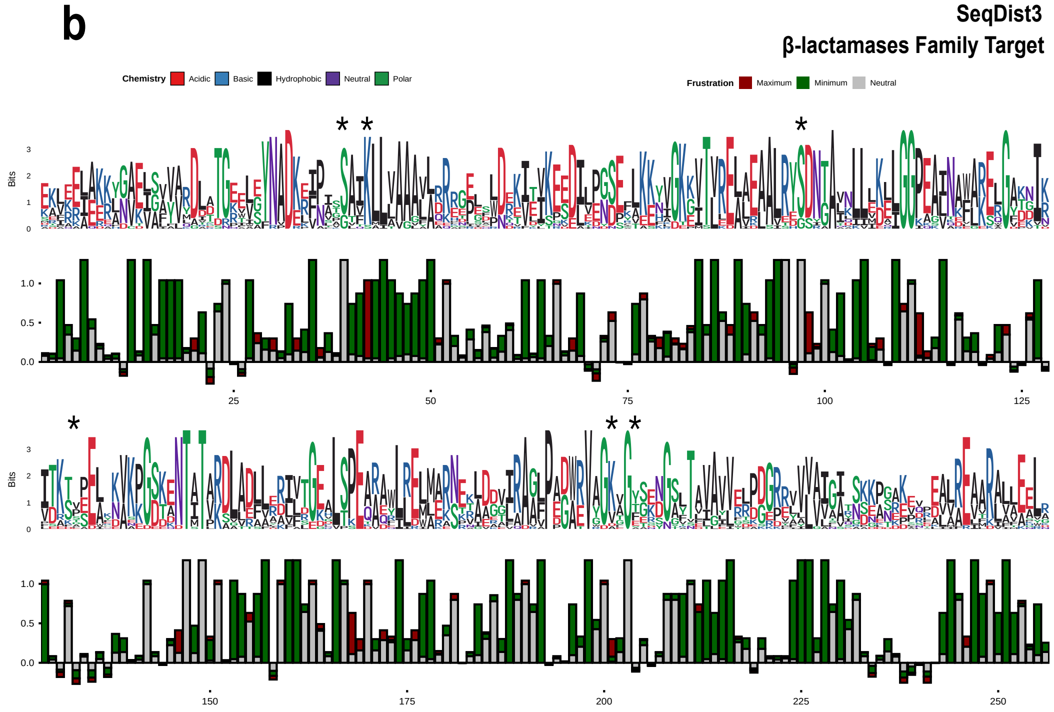


**Fig. S27.** Sequence and Frustration logo plots showing SeqIC and FrustIC values per MSA column respectively, for the Family Target approach applied to the $\beta$-lactamases native family when designing samples with Caliby. The numbering of the plots corresponds to the sequence of reference 4blm-A. Positions containing a gap in the sequence of reference are not considered in the plot. In black asterisks, we marked catalytic positions from the $\beta$-lactamases protein family. The same synthetic protein set is considered but measuring frustration conservation when SeqDist frustration parameters is setted to (a) SeqDist=12 or (b) SeqDist=3 alternatively.

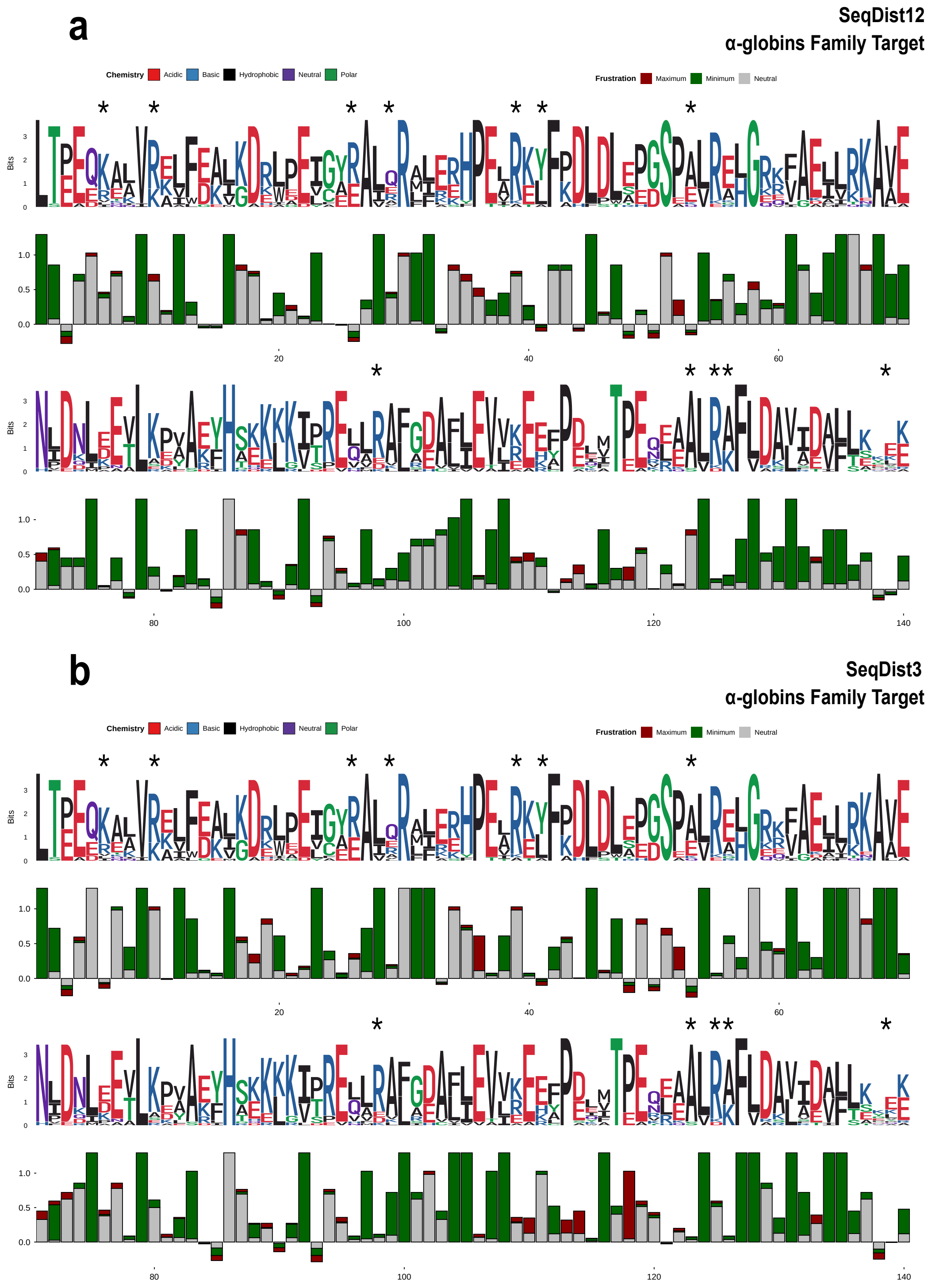


**Fig. S28.** Sequence and Frustration logo plots showing SeqIC and FrustIC values per MSA column respectively, for the Family Target approach applied to the $\alpha$-globins native family when designing samples with Caliby. The numbering of the plots corresponds to the sequence of reference 2dn1-A. Positions containing a gap in the sequence of reference are not considered in the plot. In black asterisks, we marked the functional positions of interest from the $\alpha$-globins protein family. The same synthetic protein set is considered but measuring frustration conservation when SeqDist frustration parameters is setted to (a) SeqDist=12 or (b) SeqDist=3 alternatively.

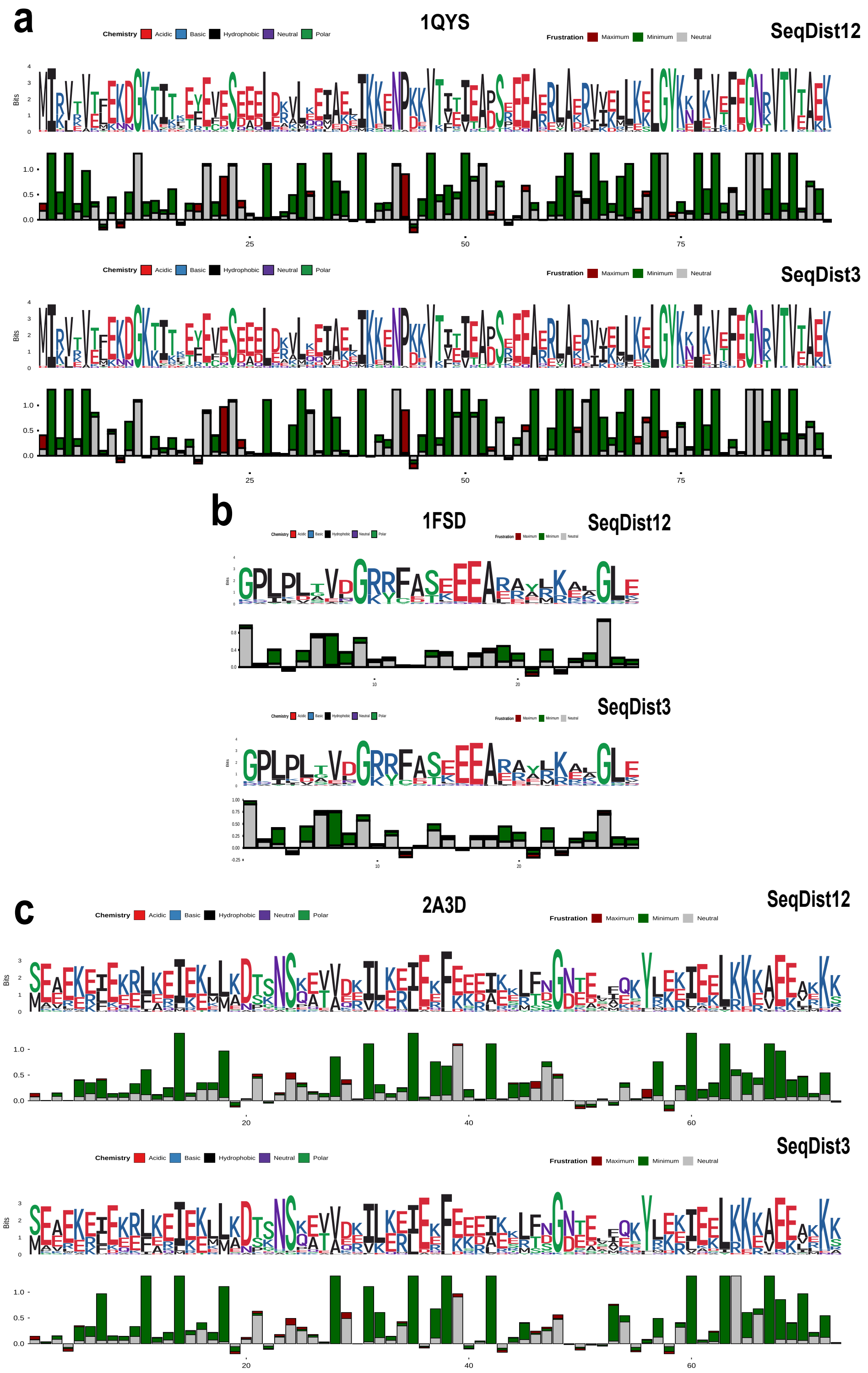


**Fig. S29.** Sequence and Frustration logo plots showing SeqIC and FrustIC values per MSA column respectively, for the Single Target approach applied to *de novo* examples: (a) 1QYS, (b) 1FSD and (c) 2A3D, when designing samples with Caliby. The numbering of the plots corresponds to the reference sequences, being these the original sequences proposed for such *de novo* proteins. Given each de novo target, the same synthetic set is considered but measuring frustration conservation when SeqDist frustration parameters is setted to SeqDist=12 or SeqDist=3 alternatively.

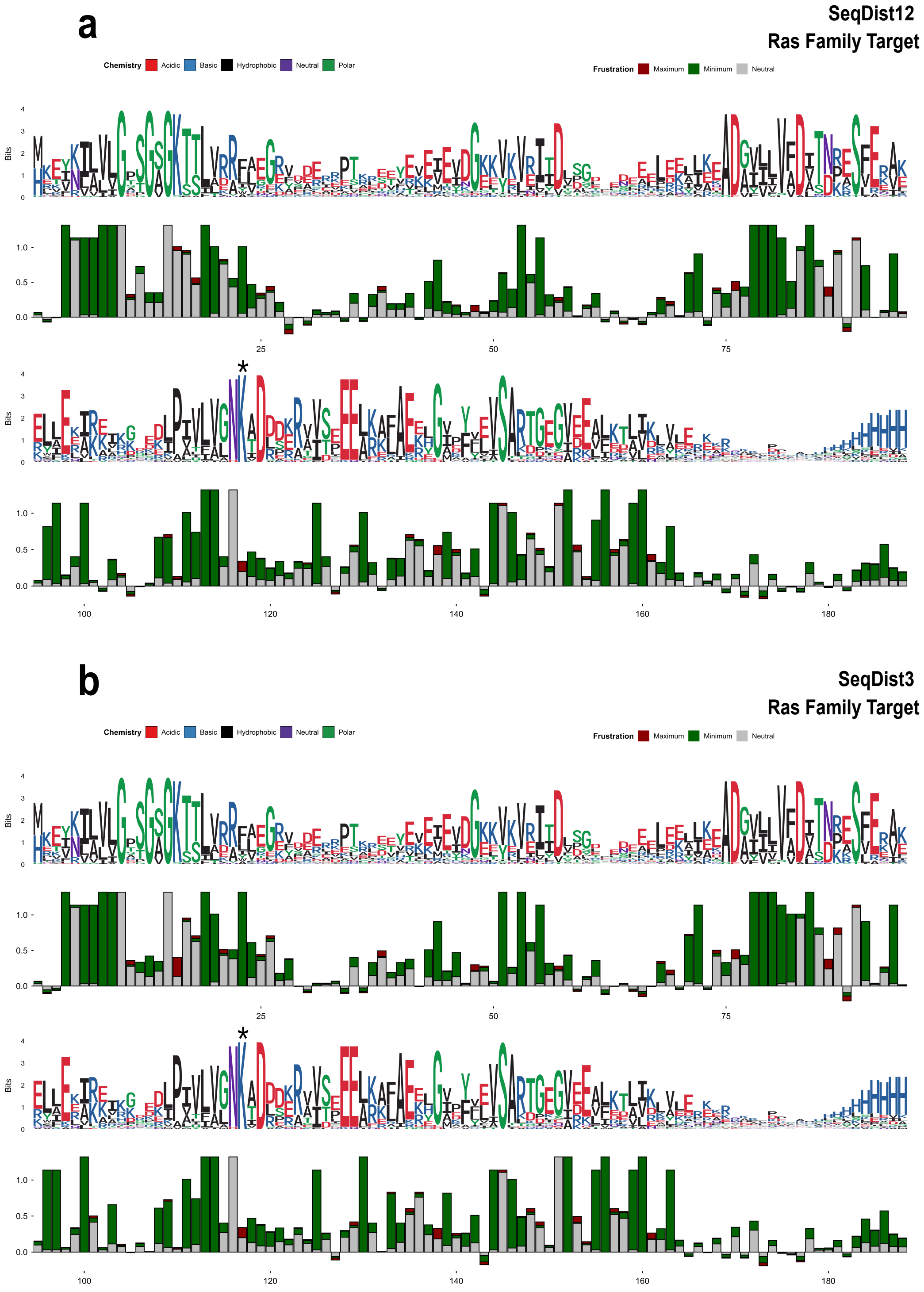


**Fig. S30.** Sequence and Frustration logo plots showing SeqIC and FrustIC values per MSA column respectively, for the Family Target approach applied to the Ras native family when designing samples with Caliby. The numbering of the plots corresponds to the sequence of reference P01116-2. Positions containing a gap in the sequence of reference are not considered in the plot. The same synthetic protein set is considered but measuring frustration conservation when SeqDist frustration parameters is setted to (a) SeqDist=12 or (b) SeqDist=3 alternatively. K117 residue is marked with *.

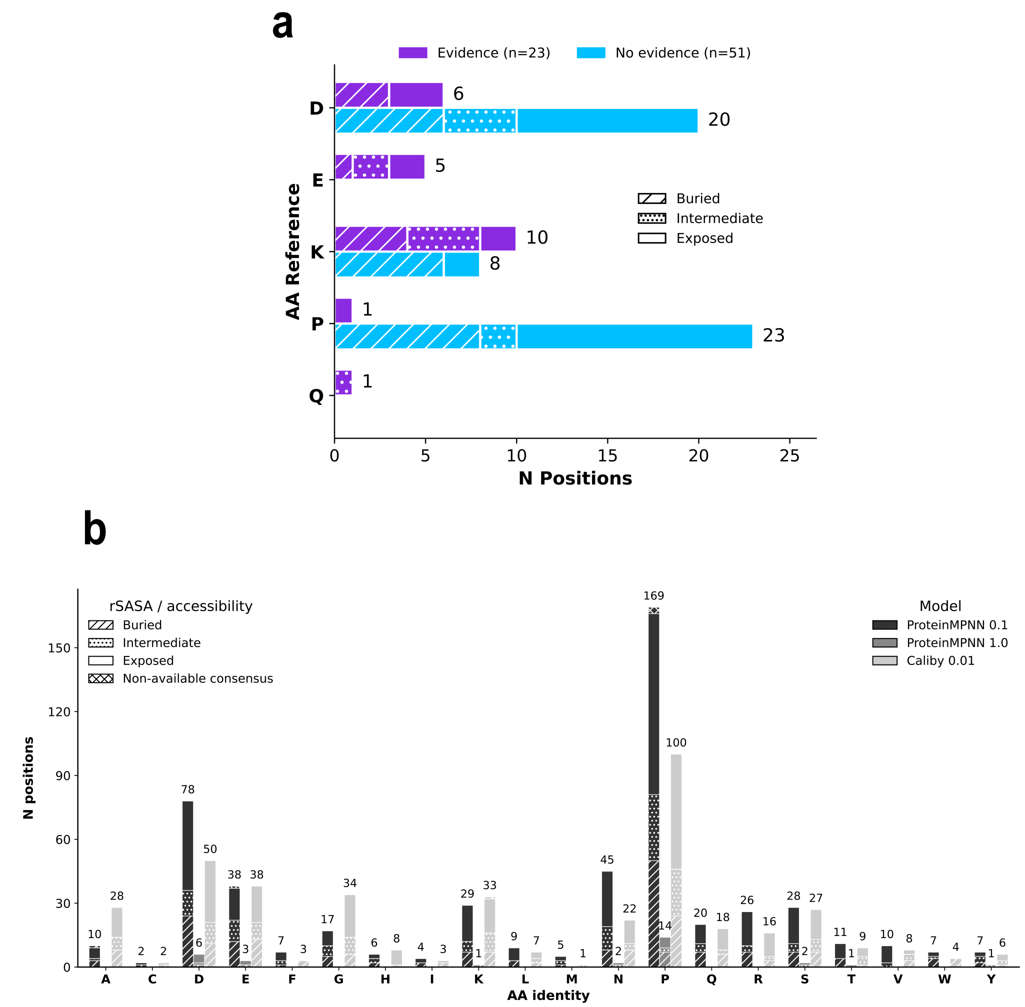


**Fig. S31.** Distribution of spandrel positions stratified by amino acid identity and relative solvent accessibility (rSASA). (a) Native spandrels, defined as positions conserved as highly frustrated across the design configurations and the native set, with (purple, $n = 23$) or without (blue, $n = 51$) functional evidence. (b) Non-native spandrels, defined as positions conserved as highly frustrated within individual design configurations but neither frustrated nor conserved in the native FunFams; intersections between design configurations are not shown. Amino acid identities correspond to those of the native reference proteins, and rSASA is the consensus relative solvent accessibility at the corresponding MSA position in the native FunFam. rSASA $\leq 0.1$ was classified as buried, $0.1 <$rSASA$< 0.3$ as intermediate, and rSASA $\geq 0.3$ as exposed.

**Table S1. Training and dataset details for ProteinMPNN re-trained versions. The first two versions were retrained using customized versions of the original ProteinMPNN dataset by (1) removing all experimentally determined entries annotated as hydrolases and (2) removing all experimentally determined entries annotated as any enzyme class, according to the E.C. number annotation in the PDB database (24). The last two versions were retrained after completing the PDB enzyme annotation using E.C. predictions from CLEAN (22) for sequences not previously categorized as enzymes in the PDB database (maximum separation score, confidence > 0.5; no hyperparameters were tuned; see Methods). Training details are provided for all four resulting models. Training num epochs indicates the number of complete passes through the training dataset used to update model weights. Backbone noise training indicates the Gaussian noise added to atomic coordinates during training to reduce memorization of crystallographic biases. Training topk indicates the number of nearest neighbors per residue used by ProteinMPNN to construct the protein structural graph during feature extraction. Best model @ epoch indicates the epoch with the lowest validation loss, whose weights were selected for subsequent sequence generation.**

| | **Re-trained ProteinMPNN versions** | | | |
|---|---|---|---|---|
| | **PDB Hydrolases** | **PDB AllEnzymes** | **CLEAN Hydrolases** | **CLEAN AllEnzymes** |
| Training num epochs | 120 | 120 | 120 | 120 |
| Backbone noise training | 0.2 | 0.2 | 0.2 | 0.2 |
| Training top_k | 48 | 48 | 48 | 48 |
| Train dataset clusters | 22517 | 19113 | 21908 | 17362 |
| Valid dataset clusters | 1349 | 1167 | 1311 | 1042 |
| Test dataset clusters | 1424 | 1212 | 1378 | 1070 |
| Best model @ epoch | 111 | 116 | 112 | 118 |

**Table S2. Residues with conserved identity and high frustration by consensus in native $\beta$-lactamases. Catalytic residues previously reported as frustrated in native $\beta$-lactamases are marked with *. Native frustration was computed using SeqDist values of 3 and 12 (sequence separation; see Methods). Single Target and Family Target design approaches were implemented using ProteinMPNN and Caliby at different sampling temperatures (model hyperparameter) and SeqDist values (Frustratometer hyperparameter). Note that Caliby was tested only at its default sampling temperature.**

| | Natives | | Single Target | | | | Family Target | | | | | |
|---|---|---|---|---|---|---|---|---|---|---|---|---|
| | SeqDist12 | SeqDist3 | SeqDist12 | | SeqDist3 | | SeqDist12 | | | SeqDist3 | | |
| | | | MPNN 0.1 | MPNN 1.0 | MPNN 0.1 | MPNN 1.0 | MPNN 0.1 | MPNN 1.0 | Caliby | MPNN 0.1 | MPNN 1.0 | Caliby |
| **K73*** | yes | yes | yes | yes | yes | yes | yes | yes | yes | yes | yes | yes |
| **D131** | yes | yes | yes | yes | yes | yes | yes | yes | yes | yes | yes | yes |
| **P145** | yes | yes | x | x | x | x | yes | x | x | yes | x | yes |
| **E166*** | yes | yes | x | x | x | x | x | x | x | x | x | x |
| **D179** | yes | yes | x | x | x | x | yes | x | x | yes | x | x |
| **D233** | yes | yes | x | x | x | x | x | x | x | x | x | x |
| **K234*** | yes | yes | yes | yes | yes | yes | yes | yes | yes | yes | yes | x |

**Table S3. Residues with conserved identity and high frustration by consensus in native $\alpha$-globins. Native frustration was computed using SeqDist values of 3 and 12 (sequence separation; see Methods). Single Target and Family Target design approaches were implemented using ProteinMPNN and Caliby at different sampling temperatures (model hyperparameter) and SeqDist values (Frustratometer hyperparameter). Note that Caliby was tested only at its default sampling temperature.**

| | Natives | | Single Target | | | | Family Target | | | | | |
|---|---|---|---|---|---|---|---|---|---|---|---|---|
| | SeqDist12 | SeqDist3 | SeqDist12 | | SeqDist3 | | SeqDist12 | | | SeqDist3 | | |
| | | | MPNN 0.1 | MPNN 1.0 | MPNN 0.1 | MPNN 1.0 | MPNN 0.1 | MPNN 1.0 | Caliby | MPNN 0.1 | MPNN 1.0 | Caliby |
| **K7** | yes | yes | x | x | x | x | x | x | x | x | x | x |
| **K11** | yes | yes | x | x | x | x | x | x | x | x | x | x |
| **E27** | yes | yes | x | x | x | x | x | x | x | x | x | x |
| **E30** | yes | yes | x | x | x | x | x | x | x | x | x | x |
| **P37** | x | yes | x | x | yes | x | x | x | x | yes | x | yes |
| **K40** | yes | yes | x | x | x | x | x | x | x | x | x | x |
| **Y42** | yes | yes | x | x | x | x | x | x | x | x | x | x |
| **Q54** | yes | yes | yes | x | yes | x | yes | x | x | x | x | x |
| **K99** | yes | yes | x | x | x | x | x | x | x | x | x | x |
| **E115** | x | x | yes | x | yes | x | x | x | x | x | x | x |
| **P119** | x | yes | yes | x | yes | x | yes | x | x | yes | yes | yes |
| **S124** | yes | x | x | x | x | x | x | x | x | x | x | x |
| **D126** | yes | yes | x | x | x | x | x | x | x | x | x | x |
| **K127** | yes | yes | x | x | x | x | x | x | x | x | x | x |
| **Y140** | yes | yes | yes | x | x | x | x | x | x | x | x | x |

**Table S4. Conservation of high frustration across protein design configurations over 488 FunFams relative to the native baseline. Positions with conserved high frustration (FrustIC > 0.5) were retained. No sequence conservation threshold (SeqIC) was applied, while MSA positions with zero or non-available SeqIC were excluded. The resulting sets were intersected across design configurations and further divided according to the presence or absence of functional evidence, with percentages calculated over frustrated native entries. Adaptive frustration refers to positions that are conserved and highly frustrated in native sequences but not in any design configuration, consistent with putative adaptive roles. Native spandrels refer to positions that are conserved and highly frustrated in native sequences and retained in other or all design configurations, consistent with fold-specific constraints. Non-native spandrels refer to positions that are conserved and highly frustrated in designed sequences (in any model) but not in native sequences, as alternative fold-specific constraints.**

| | Intersection | Evidence | No evidence | Total | % Native entries |
|---|---|---|---|---|---|
| **Adaptive frustration** | Native NOT recovered by ProteinMPNN 0.1 | 227 | 1708 | 1935 | 76.66 |
| | Native NOT recovered by ProteinMPNN 1.0 | 277 | 2020 | 2297 | 91.01 |
| | Native NOT recovered by Caliby 0.01 | 294 | 2061 | 2355 | 93.30 |
| | Native NOT recovered by any configuration | 218 | 1667 | 1885 | 74.68 |
| **Native spandrels** | Native | 332 | 2192 | 2524 | 100.00 |
| | Native & ProteinMPNN 0.1 | 105 | 484 | 589 | 23.34 |
| | Native & ProteinMPNN 1.0 | 55 | 172 | 227 | 8.99 |
| | Native & Caliby 0.01 | 38 | 131 | 169 | 6.70 |
| | Native & ProteinMPNN 0.1 & Caliby 0.01 | 34 | 103 | 137 | 5.43 |
| | Native & all 3 configurations | 23 | 51 | 74 | 2.93 |
| **Non-native spandrels** | ProteinMPNN 0.1 only | 27 | 501 | 528 | 20.92 |
| | ProteinMPNN 1.0 only | 3 | 27 | 30 | 1.19 |
| | Caliby 0.01 only | 22 | 395 | 417 | 16.52 |
| | MPNN 0.1 & MPNN 1.0, not Native | 3 | 21 | 24 | 0.95 |
| | MPNN 0.1 & Caliby, not Native | 5 | 107 | 112 | 4.44 |
| | MPNN 1.0 & Caliby, not Native | 1 | 8 | 9 | 0.36 |
| | All 3 configurations, not Native | 1 | 8 | 9 | 0.36 |

#### Data, Materials, and Software Availability

The custom datasets used for retraining ProteinMPNN and the resulting model weights are available at repository-weights. All input/output data needed to reproduce the main results of this article as well as the intermediate analyses are available at repository-data. Related code and documentation available at repository-code.

#### References


1. AO Rausch, et al., FrustratometeR: an R-package to compute local frustration in protein structures, point mutants and MD simulations. *Bioinforma. (Oxford, England)* **37**, 3038–3040 (2021).
2. MI Freiberger, et al., Local energetic frustration conservation in protein families and superfamilies. *Nat. Commun.* **14**, 8379 (2023).
3. RG Parra, et al., Frustraevo: a web server to localize and quantify the conservation of local energetic frustration in protein families. *Nucleic Acids Res.* **52**, W233–W237 (2024).
4. J Dauparas, et al., Robust deep learning-based protein sequence design using ProteinMPNN. *Sci. (New York, N.Y.)* **378**, 49–56 (2022).
5. DU Ferreiro, JA Hegler, EA Komives, PG Wolynes, Localizing frustration in native proteins and protein assemblies. *Proc. Natl. Acad. Sci.* **104**, 19819–19824 (2007).
6. RW Shuai, T Lu, S Bhatti, P Kouba, PS Huang, Ensemble-conditioned protein sequence design with Caliby (2025) ISSN: 2692-8205 Pages: 2025.09.30.679633 Section: New Results.
7. T Lu, et al., Conditional Protein Structure Generation with Protpardelle-1c. *bioRxiv: The Prepr. Serv. for Biol.* p. 2025.08.18.670959 (2025).
8. M Jenik, et al., Protein frustratometer: a tool to localize energetic frustration in protein molecules. *Nucleic Acids Res.* **40**, W348–351 (2012).
9. JR Knox, PC Moews, -Lactamase of *Bacillus licheniformis* 749/C. *J. Mol. Biol.* **220**, 435–455 (1991).
10. SK Burley, et al., Rcsb protein data bank: powerful new tools for exploring 3d structures of biological macromolecules for basic and applied research and education in fundamental biology, biomedicine, biotechnology, bioengineering and energy sciences. *Nucleic acids research* **49**, D437–D451 (2021).
11. DM Taylor, et al., Unique Diacidic Fragments Inhibit the OXA-48 Carbapenemase and Enhance the Killing of Escherichia coli Producing OXA-48. *ACS Infect. Dis.* **7**, 3345–3354 (2021).
12. V Stojanoski, et al., Removal of the Side Chain at the Active-Site Serine by a Glycine Substitution Increases the Stability of a Wide Range of Serine -Lactamases by Relieving Steric Strain. *Biochemistry* **55**, 2479–2490 (2016).
13. SD Lahiri, et al., Structural Insight into Potent Broad-Spectrum Inhibition with Reversible Recyclization Mechanism: Avibactam in Complex with CTX-M-15 and Pseudomonas aeruginosa AmpC -Lactamases. *Antimicrob. Agents Chemother.* **57**, 2496–2505 (2013).
14. P Hinchliffe, et al., Penicillanic Acid Sulfones Inactivate the Extended-Spectrum -Lactamase CTX-M-15 through Formation of a Serine-Lysine Cross-Link: an Alternative Mechanism of -Lactamase Inhibition. *mBio* **13**, e01793–21 (2022).
15. SY Park, T Yokoyama, N Shibayama, Y Shiro, JRH Tame, 1.25 Å Resolution Crystal Structures of Human Haemoglobin in the Oxy, Deoxy and Carbonmonoxy Forms. *J. Mol. Biol.* **360**, 690–701 (2006).
16. B Kuhlman, et al., Design of a novel globular protein fold with atomic-level accuracy. *Sci. (New York, N.Y.)* **302**, 1364–1368 (2003).
17. BI Dahiyat, SL Mayo, Probing the role of packing specificity in protein design. *Proc. Natl. Acad. Sci.* **94**, 10172–10177 (1997).
18. ST Walsh, H Cheng, JW Bryson, H Roder, WF DeGrado, Solution structure and dynamics of a de novo designed three-helix bundle protein. *Proc. Natl. Acad. Sci.* **96**, 5486–5491 (1999).
19. VP Waman, et al., CATH 2024: CATH-AlphaFlow Doubles the Number of Structures in CATH and Reveals Nearly 200 New Folds. *J. Mol. Biol.* **436**, 168551 (2024).
20. RE Hackett, et al., Investigating enzyme function by geometric matching of catalytic motifs. *bioRxiv* pp. 2026–02 (2026).
21. P Maietta, et al., Firedb: a compendium of biological and pharmacologically relevant ligands. *Nucleic acids research* **42**, D267–D272 (2014).
22. T Yu, et al., Enzyme function prediction using contrastive learning. *Sci. (New York, N.Y.)* **379**, 1358–1363 (2023).
23. Y Zhang, J Skolnick, TM-align: a protein structure alignment algorithm based on the TM-score. *Nucleic Acids Res.* **33**, 2302–2309 (2005).
24. SK Burley, et al., Updated resources for exploring experimentally-determined pdb structures and computed structure models at the rcsb protein data bank. *Nucleic acids research* **53**, D564–D574 (2025).
